# Mismatch repair dissection by in vivo RNAi reveals dose-dependent modulators of somatic instability and proteome remodeling in Huntington’s disease

**DOI:** 10.64898/2026.06.19.733435

**Authors:** Jillian Belgrad, Todd M. Greco, Ellen Sapp, Ashley Summers, Daniel O’Reilly, Eric Luu, Josiah E Hutton, Nozomi Yamada, Hassan H. Fakih, Raymond Furgal, Dimas Echeverria, Nicholas McHugh, Brianna Bramato, Chantal Furguson, Samuel Hildebrand, Sarah Allen, Nicholas Gaston, David Cooper, Allison Maebius, Katherine Y. Gross, Thomas F. Vogt, Michael Finley, Brinda Prasad, Marian DiFiglia, Ileana M. Cristea, Neil Aronin, Anastasia Khvorova

## Abstract

Human and mouse genetics have established mismatch repair (MMR) as a central mediator of somatic repeat expansion, a key pathogenic process in Huntington’s disease (HD) and related disorders. How individual MMR components function within the intact mammalian brain and interact with broader cellular networks remains poorly understood. We screened more than 500 chemically stabilized siRNAs targeting 10 MMR genes and used interventional RNAi in the Q111 HD mouse model to systematically dissect MMR function in vivo. MSH3 and PMS1 emerged as the most dose-sensitive regulators of somatic expansion but displayed markedly different effects on proteome stability. Quantitative proteomics generated an in vivo atlas of MMR component abundance and cross-regulation in the mammalian CNS, uncovering extensive connectivity between DNA repair, transcriptional regulation, chromatin remodeling, and mitochondrial biology. Together, these findings establish a systems-level framework linking MMR biology to neuronal function and offer mechanistic insight into selective neuronal vulnerability in HD.

## INTRODUCTION

Huntington’s disease (HD) is a fatal, autosomal dominant neurodegenerative disorder caused by 36 or more consecutive CAG repeats in exon 1 of the huntingtin (*HTT)* gene.^1–4^ Somatic expansion of the inherited CAG repeats occurs over the lifetime of the HD patient and drives disease progression and neuron cell death.^5–8^ Genome-wide association studies have implicated DNA mismatch repair (MMR) genes as key modulators of this process, with variants in MutS homolog 3 (*MSH3)*, MutL homolog 1 (*MLH1)*, and others correlating with altered age of onset.^9–12^ Subsequent studies using knockout models, induced pluripotent stem cell-derived neurons, and CRISPR interference have genetically validated MMR as a key driver of somatic expansion *in vitro* and *in vivo*.^11,13–26^

Somatic expansion driven by MMR occurs when DNA slip-outs arise from strand misalignment in R-loops that form during transcription, particularly in the repetitive and self-complementary CAG region of *HTT*.^17,27–29^ These DNA slip outs are recognized by the heterodimeric MutSβ complex composed of MutS homolog 2 (MSH2) and MSH3. Upon binding, MutSβ recruits one or more heterodimeric MutL complexes, all derived through gene duplication and divergence from a common ancestral ATPase.^30,31^ These include MutLα composed of MLH1 and PMS2, MutLβ composed of MLH1 and PMS1, and MutLγ composed of MLH1 and MLH3, which together initiate downstream repair processes.^28,29,32^ Emerging evidence shows that blocking somatic expansion combined with lowering mutant HTT expression more effectively slows HD progression than reducing mutant HTT alone, positioning MMR as a key therapeutic axis for HD.^6,8,33^

Although mismatch repair (MMR) is a known driver of somatic repeat expansion^11,13–26^, how individual MMR components shape CAG repeat expansion in an interventional context, and whether they represent viable therapeutic targets, remains poorly understood. By using small interfering RNAs (siRNAs) to target MMR genes we could evaluate in a graded fashion dose-dependent modulation and functional interactions within the MMR network. Recent advances in therapeutic siRNAs, including divalent siRNA scaffolds that provide broad CNS distribution, efficient neuronal uptake, and sustained target silencing for up to six months after a single administration,^34,35^ have enabled long-term modulation of gene expression in the brain. These advances provide the technical foundation necessary to evaluate the role of individual mismatch repair (MMR) factors in somatic repeat expansion and disease progression in Huntington’s disease models.

Here, we systematically investigated MMR genes using therapeutic siRNAs in a HD mouse model that exhibits somatic expansion in the brain. Over 500 fully chemically stabilized siRNAs to ten core MMR genes were designed, screened, and validated for efficacy in both human cell and mouse models. The two most efficacious siRNAs for each target MMR were synthesized in a divalent scaffold and evaluated in an HD expansion mouse model where quantitative relationships were established between the degree of acute MMR gene knockdown and the extent of somatic expansion. Global quantitative proteomics of striatum from treated mice provided a map of MMR protein abundance in the mouse brain and revealed extensive interconnectivity between distinct MMR complexes and transcriptional machinery, which likely contributes to the regional vulnerability observed in somatic expansion. Results showed that divalent siRNA targeting either PMS1 or MSH3 effectively blocked somatic expansion, even with partial target silencing. However, silencing of PMS1, but not MSH3, led to greater proteome remodeling, suggesting a broader biological role for PMS1 and highlighting MSH3 as a potentially more selective interventional target.

## RESULTS

### *In vitro* screening identifies top-performing siRNAs for MMR target knockdown

To systematically modulate the MMR pathway using siRNAs, we screened a panel of fully chemically modified siRNAs for the following MMR targets: *MSH2/Msh2*, *MLH1/Mlh1*, MutL homolog 3 (MLH3/Mlh3), MutS homolog 6 (*MSH6/Msh6*), polymerase delta 1 (*POLD1/Pold1*), polymerase delta 3 (*POLD3/Pold3*), exonuclease 1 (*EXO1/ Exo1*), *PMS1/Pms1*, *PMS2/Pms2*, and Fanconi anemia-associated nuclease 1 (*FAN1/Fan1*). These targets were selected based on their association with the age of HD onset in patients and their roles in MutSβ, MutL, and other known MMR complexes.^8,10–14,17,18,23,25,26,29,36–42^ The siRNA sequences were designed using a bioinformatic algorithm that predicts highly specific and functional siRNAs.^43^ For each target gene, three panels (∼48 siRNAs per target) were generated: for the human transcript, the mouse transcript, and for human and mouse using conserved regions between the two species (Figure S1B-K).

All siRNAs were fully chemically stabilized with 2′-O-methyl and 2′-fluoro modifications along with a partially phosphorothioated backbone^44,45^ to promote efficacy, potency, and durability *in vitro* and *in vivo* (Figure S1A).^44,46^ For in vitro screening siRNAs were conjugated to cholesterol to enable passive cellular uptake and mimic *in vivo* trafficking pathways. Dual-homology siRNAs were tested in human (HeLa) and mouse (N2a) cell lines. If a dual-homology siRNA demonstrated strong and consistent knockdown in both cell types, it was prioritized for further evaluation to facilitate cross-species translation. When dual-homology siRNAs were not sufficiently potent, mouse-specific siRNAs were selected for further evaluation. To identify siRNA leads for in vivo evaluation in mice, potent compounds were assessed in a 7-point dose-response assay in N2a cells. Top performers achieved >50% target mRNA knockdown and an IC50 < 500 nM in passive uptake (Figure S2). Human-targeting siRNAs were similarly validated in HeLa cells to establish a corresponding toolset for modulating MMR in human systems (Figure S3).

This screening approach resulted in a validated panel of potent, fully chemically modified siRNAs targeting human and mouse MMR pathways. A summary of the *in vitro* siRNA panel, including sequences and chemical modification patterns, is presented in Table S1 generating a novel resource for systematic MMR evaluation in human and mouse models for the community.

### Divalent siRNAs significantly lower target MMR mRNA and protein *in vivo* two months post-injection

We next evaluated siRNA modulation of MMR within the context of regional vulnerability of HD somatic instability using the Q111 knock-in mouse model.^26,47^ Q111 mice carry human exon 1 with 110–120 CAG repeats in the mouse *Htt* locus and exhibit robust age-dependent somatic expansion in the caudate putamen, where the degree of expansion is measurable within a two-month timeframe.^14,22^ The two most potent siRNAs per MMR target identified in the in vitro screening were synthesized in a fully chemically modified divalent scaffold. This scaffold incorporates a 5′-(E)-vinylphosphonate (5′-VP) modification on the guide strand, enhancing phosphate stability and optimizing interaction with the MID domain of argonaute-2,^48^ together with a linked divalent sense strand. The resulting architecture enables robust, durable CNS gene silencing with improved stability and biodistribution in vivo (Figure 1A).^34^ At three months old, mice received a single bilateral intracerebroventricular injection of 10 nmol (240 µg in 10 µL) divalent siRNA. Target mRNA and protein levels, and somatic instability were assessed two months later (Figure 1B).

**Figure 1.**
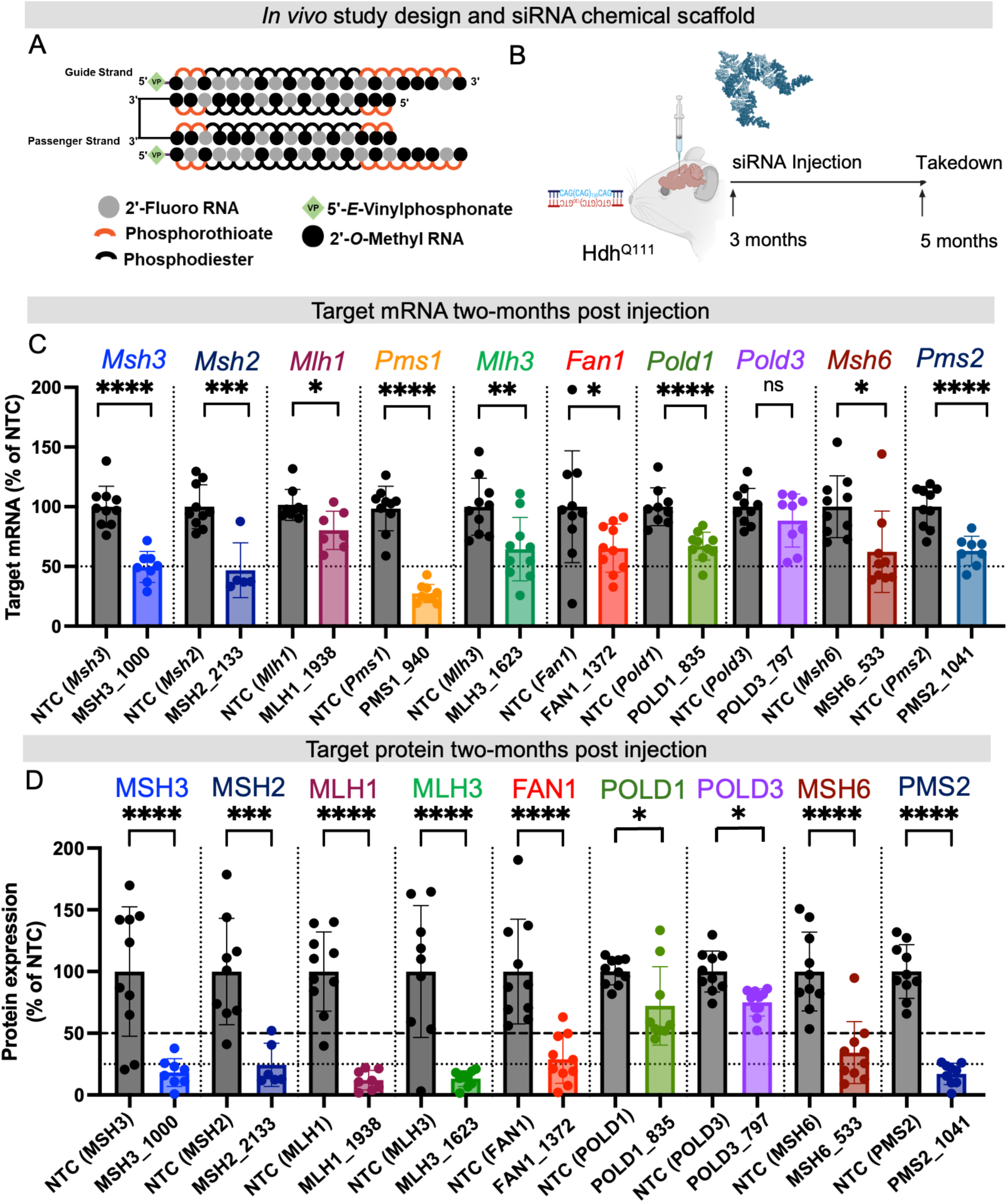
Mismatch repair pathway targets silenced by divalent siRNA two-months post injection in the Q111 striatum. (A) Divalent siRNA scaffold and chemical modification pattern. (B) *In vivo* study paradigm. 10 nmol/10 µL divalent siRNA injected intracerebroventricularly into Q111 mice at 3 months of age with outcomes measured at 5 months of age. (C) Striatum mRNA levels detected by branched DNA assay for top-performing siRNA per target. Two-tailed t-test NTC versus target. (D) Striatum protein levels quantified from western blot for top-performing siRNA per target. Two-tailed t-test NTC versus target. N=6-10 mice per group.

Mice (n = 6–10 per group) were randomized by sex, age, and inherited tail CAG length (Figure S4) to balance the groups. Controls included artificial cerebrospinal fluid (aCSF),^49^ a non-targeting scrambled divalent siRNA (NTC), and MSH3-targeting divalent siRNA previously validated to block somatic expansion in HD mouse models^22^. MMR target characterization was conducted in Q111 Huntington’s disease mice across three independent studies, each involving 100–120 animals. Appropriate control groups were included in every experimental arm to ensure direct comparison across treatment conditions with MSH3 silencing and impact on expansion serving as a positive control in each *in vivo* experimental cohort (Figure S5). For somatic expansion assays, a baseline cohort of age-matched Q111 mice was sacrificed at the time of injection to establish the starting levels of striatal CAG instability. All *in vivo* validated siRNA sequences and modification patterns are listed in Table S2.

At two months post-treatment with divalent siRNA, significant reductions in target mRNA expression were observed in the caudate putamen (Figure 1C; reported as % of NTC): *Msh3* to 49.6% ± 12 (p < 0.0001), *Msh2* to 46.0% ± 20 (p = 0.0003), *Mlh1* to 80.0% ± 14 (p = 0.0109), *Pms1* to 27.3% ± 7 (p < 0.0001), *Mlh3* to 64.5% ± 25 (p = 0.001), *Fan1* to 65.2% ± 19 (p = 0.045), *Pold1* to 67.0% ± 11 (p < 0.0001), *Msh6* to 62.3% ± 32 (p = 0.0114), and *Pms2* to 63.0% ± 11 (p < 0.0001). Reduction of *Pold3* mRNA (88.0% ± 20) was not statistically significant (p = 0.1979). Figure 1 shows the knockdown of the more potent siRNA for each target; all remaining brain regions and siRNAs investigated are shown in Figure S6.

To evaluate target protein levels, we screened most commercially available antibodies for their specificity against mouse MMR proteins; data on both functional and non-functional antibodies are provided in Table S3. Validated antibodies used for protein detection are listed in Table S4 and Figure S7. In the caudate putamen, we observed robust protein knockdown (50–80%) across targets relative to NTC. Protein silencing generally exceeded mRNA knockdown (Figure 1D; Figures S8 and S9), consistent with published data.^50^ MSH3 protein levels were reduced to 18.2 ± 10% (p < 0.0001), MSH2 to 24.4 ± 16% (p < 0.0001), MLH1 to 12.1 ± 7% (p < 0.0001), MLH3 to 12.3 ± 6% (p < 0.0001), FAN1 to 29.7 ± 20% (p < 0.0001), POLD1 to 65.7 ± 23% (p = 0.018), POLD3 to 75.5 ± 10% (p = 0.0289), MSH6 to 37.3 ± 24% (p < 0.0001), and PMS2 to 19.1 ± 6% (p < 0.0001). While a reliable PMS1 antibody was unavailable, *Pms1* mRNA was silenced to ∼27%, suggesting at least a similar level of reduction at the protein level. In summary, we achieved robust *in vivo* modulation (>50% mRNA, >75% protein) for eight of ten MMR targets, and moderate silencing (25–35%) of POLD1 and POLD3.

### Systematic *in vivo* silencing of MMR genes reveals differential effects on striatal somatic repeat expansion

We next assessed the impact of knocking down key MMR targets on somatic CAG repeat instability (Figure 2A,B). We first focused on primary MMR machinery commonly shared across different pathways, including MSH3 and MSH2, which form MutSβ; MLH1, a subunit of all MutL complexes; MSH6; and FAN1 (Figure 2C). In canonical MMR, MSH6 pairs with MSH2 to form the MutSα complex, which recognizes base–base mismatches and small (1–2 bp) insertion–deletion loops to initiate repair. FAN1 binds and sequesters MLH1^51^, and may also compete directly with MutSβ for recognition of DNA slipouts.^52^ Somatic expansion was quantified using the somatic instability index, which describes the skew of CAG repeat lengths with increasing index number associated with greater CAG expansion.^53^ Treatment groups were compared to the NTC group. Results obtained in caudate putamen from the most potent divalent siRNA for each MMR target are shown in Figure 2. Somatic instability changes from additional siRNAs are reported in Figure S10.

**Figure 2.**
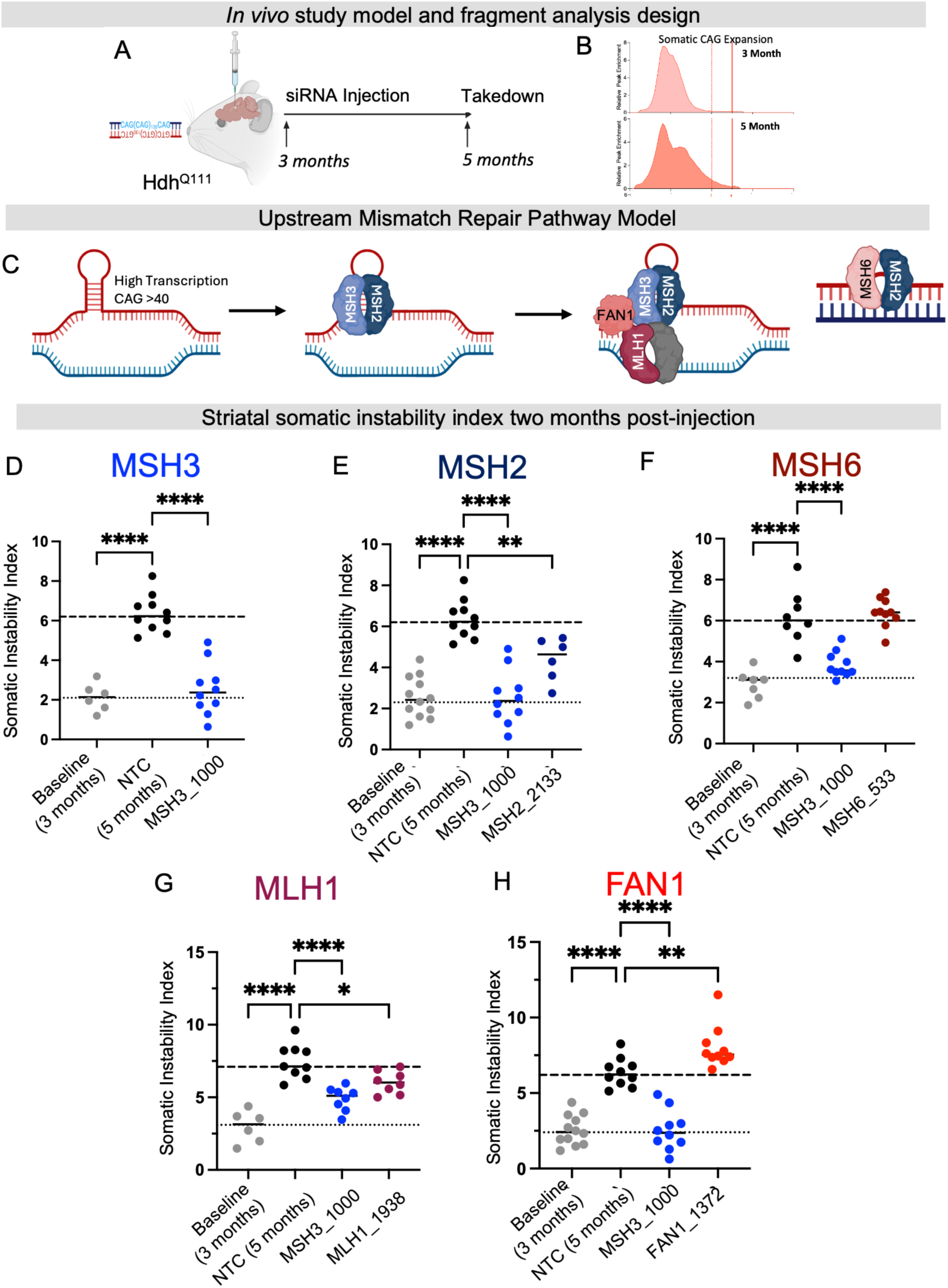
siRNA-mediated lowering of upstream mismatch repair targets alters Q111 striatal somatic instability *in vivo*. (A) *in vivo* study paradigm (B) representative fragment analysis curves used to quantify somatic instability index. (C) Schematic of upstream mismatch repair targets at repetitive CAG locus. (D-H) Baseline is striatal somatic instability of an untreated Q111 cohort at the time of injection. Non-targeting control (NTC), MSH3 controls and target siRNA are striatal somatic instability index of Q111 mice two-months post injection at age 5 months. Striatal somatic instability with top performing siRNA against (D) MSH3, MSH3_1000 (E) MSH2, MSH2_2133 (F) MSH6, MSH6_533 (G) MLH1, MLH1_1938, (H) FAN1, FAN1_1372. Fragment analysis quantified using somatic instability index. Each dot is the somatic instability index quantified from one mouse. N = 6-10/group. Statistics are one-way ANOVA vs NTC. * p < 0.05, ** p < 0.01, *** p < 0.001, **** p < 0.0001.

Consistent with our previous work,^22^ divalent siRNA targeting MSH3 completely blocked expansion (p < 0.0001; Figure 2D). MSH2 knockdown also significantly reduced expansion (p = 0.0035; Figure 2E), though less effectively than MSH3. Similarly, MLH1 knockdown significantly reduced instability (p = 0.0156; Figure 2G), but residual expansion from baseline occurred. These results differ from published homozygous genetic knockout studies of MSH2 or MLH1 that eliminate expansion, suggesting that even low levels of MSH2 or MLH1 are enough to facilitate complex formation and activity. MSH6 knockdown did not significantly impact expansion with a slide bias toward increasing instability (p = 0.9272; Figure 2F). FAN1 knockdown significantly increased expansion (p = 0.0082; Figure 2H), a well-documented phenomenon.^10–13^ Overall, these findings reveal a differential dose sensitivity of MMR components in regulating somatic repeat expansion.

We next examined PMS1, PMS2, and MLH3, each of which can bind MLH1 to form MutLβ, MutLα, and MutLγ, respectively (Figure 3A). The MutL complexes are essential for the repair and resolution of single-stranded DNA slip-outs following MutSβ binding.^17,29,37^ Previous studies suggest that individual MutL complexes may play distinct roles in repeat expansion. By systematically targeting the unique subunit of each MutL complex to selectively suppress their activity, we aimed to explore the individual role of each complex in somatic expansion *in vivo*.

**Figure 3.**
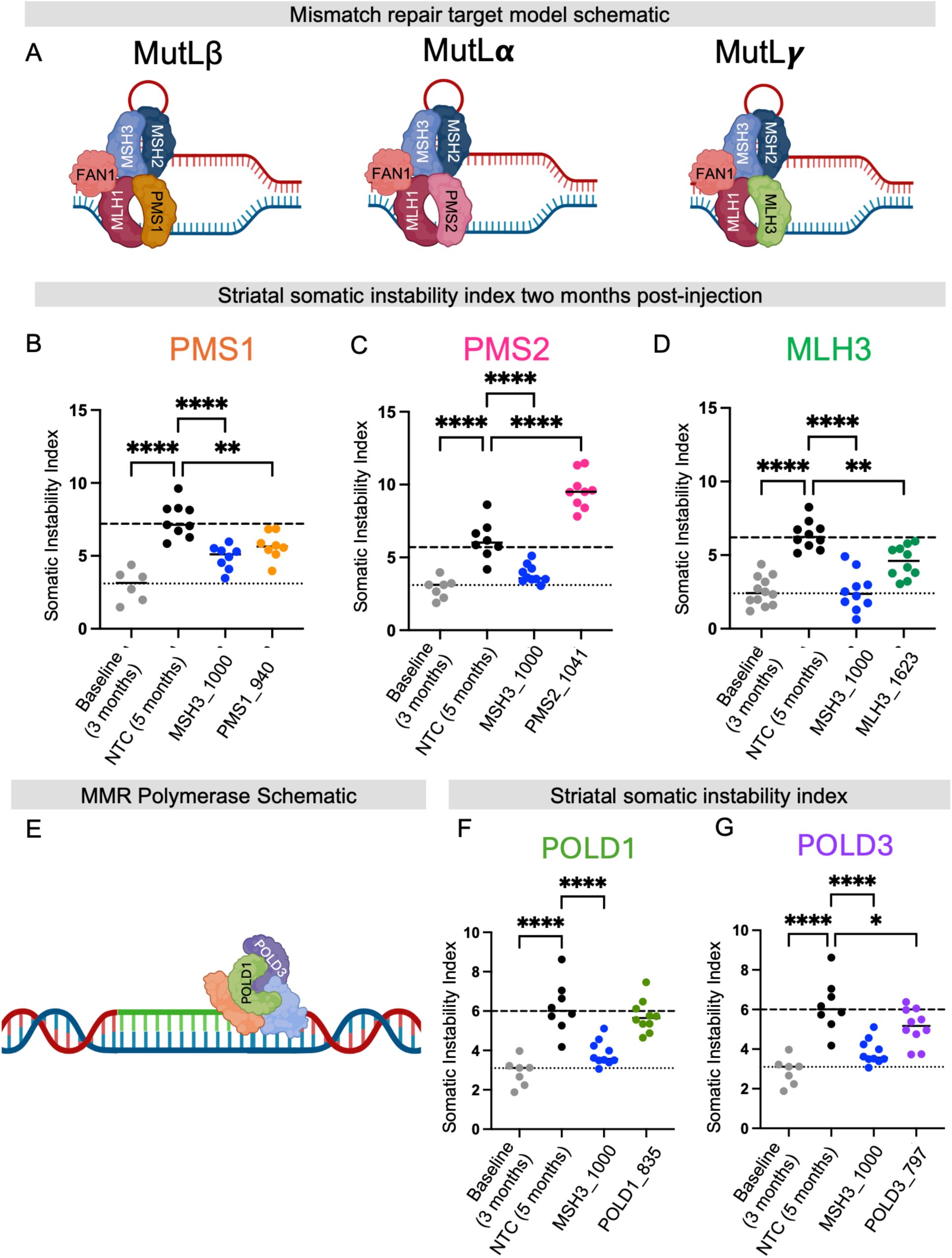
Interventional lowering of mismatch repair targets unique to MutLß, MutLα, MutLγ and polymerase. (A) Schematic of mismatch repair targets unique to MutLß, MutLα or MutLγ at a reprehensive repetitive CAG locus. (B-D) Baseline is striatal somatic instability of an untreated Q111 cohort at the time of injection. Non-targeting control (NTC), MSH3 controls and target siRNA are striatal somatic instability index of Q111 mice two-months post injection at age 5 months. Striatal somatic instability with top performing siRNA against (D) PMS1, PMS1_940, (C) PMS2, PMS2_1041, (D) MLH3, MLH3_1623. (E) Schematic of polymerase delta 1 (POLD1) or polymerase delta 3 (POLD3) during a mismatch repair expansion event. (F) Baseline is striatal somatic instability of an untreated Q111 cohort at the time of injection. Non-targeting control (NTC), MSH3 controls and target siRNA are striatal somatic instability index of Q111 mice two-months post injection at age 5 months. Striatal somatic instability with top performing siRNA against (F) POLD1, POLD1_835, (G) POLD3, POLD3_797. Fragment analysis quantified using somatic instability index. Each dot is the somatic instability index quantified from one mouse. N = 6-10/group. Statistics are one-way ANOVA vs NTC. * p < 0.05, ** p < 0.01, *** p < 0.001, **** p < 0.0001.

PMS1 knockdown significantly reduced expansion, similar to MSH3 knockdown (p = 0.0032; Figure 3B). MLH3 silencing also reduced somatic expansion (p = 0.0018; Figure 3D), although to a lesser extent than MSH3. Because the degree of target modulation was comparable across compounds, the observed differences in efficacy likely reflect biological differences among the targets rather than variations in silencing efficiency. These findings suggest that MLH3 may be particularly sensitive to the extent of target suppression. This interpretation is consistent with previous studies showing that homozygous knockout or catalytic inactivation of MLH3, as well as genetic ablation of PMS1, effectively prevents somatic expansion, highlighting the potentially nonlinear and dose-dependent contribution of these factors to repeat instability.^8,23^ PMS2 knockdown caused a robust increase in instability (p < 0.0001; Figure 3C), enhancing expansion in the striatum to a higher degree than FAN1. PMS2 as a protector from expansion has also been observed in Q111 mice via PMS2 excision,^21^ but since MutLα drives most repair in human cells, the result is surprising. This increase in somatic instability was also observed in the medial cortex, a brain region characterized by substantially slower rates of repeat expansion, where FAN1 knockdown did not produce a statistically significant effect (Figure S11). Collectively, these findings demonstrate that MutLβ and MutLγ, but not MutLα, drive *in vivo* somatic repeat expansion in the caudate putamen of Q111 mice. PMS1 appears to be a top therapeutic target, comparable to MSH3.

Finally, we evaluated the downstream replication factors, *POLD1* and *POLD3* (Figure 3E). Achieving robust *in vivo* knockdown of both genes proved challenging, despite identifying multiple highly potent siRNAs *in vitro*, suggesting a potential for tight regulatory control or presence of compensatory mechanisms for these essential polymerases. *POLD1* knockdown had no significant effect on expansion (Figure 3F, p = 0.5679), while *POLD3* knockdown led to a modest but statistically significant reduction in expansion (Figure 3G, p = 0.0486). Although the moderate knockdown of each protein complicates the interpretation of their functional impact, the fact that partial reduction of *POLD3* affected expansion supports a role for DNA polymerase activity in driving somatic instability, consistent with prior work.^21^

In summary, targeted interventional modulation of MSH3 and PMS1 effectively blocks CAG expansion in the HD mouse striatum, modulation of MSH2, MLH1, MLH3, and POLD3 slows this expansion, and modulation of FAN1 and PMS2 accelerates expansion. Compared to prior genetic excision studies,^14,15,21,23,26^ these outcomes show both concordant and discordant effects, suggesting that somatic expansion is shaped not only by the degree of MMR gene silencing—even when targets are reduced by >80%—but also by the absolute abundance of each factor and its interactions within the broader DNA repair network. Such differential sensitivity may underlie the selective vulnerability of specific neuronal subtypes in HD.

### Mapping the abundance and modulation of DNA repair and handling proteins in the mouse caudate putamen

MMR is highly dynamic and tightly regulated, mainly at the protein level.^29^ To explore these dynamics and assess how selective siRNA-mediated silencing of individual MMR components impacts broader DNA repair and handling (DNA-RH) networks at the protein level, we performed quantitative mass spectrometry (MS). MS provides an unbiased, quantitative, and high-resolution overview of the proteome, enabling the characterization not only of relative target protein levels but also of the stability and interdependence of the MMR network in the context of HD.

A custom experimental peptide spectral library was prepared as described in Methods. Deep proteome profiling was obtained using striatal tissues from untreated Q111 mice. Proteome coverage was maximized by performing basic pH reverse-phase peptide pre-fractionation and extended liquid chromatography gradients (Figure 4A and Material & Methods). The library spectra from existing proteomic datasets were used from nuclear and cytoplasmic-enriched fractions isolated from striatal tissues of Q20 knock-in mice^54^ and tryptic peptides from recombinant human MMR complexes (Greco et al, in preparation). The resulting qualitative spectral library encompassed over 180,000 unique peptides covering ∼10,000 proteins, including low-abundance siRNA targets such as FAN1, MLH1, and MSH3 (Figure 4A).

**Figure 4.**
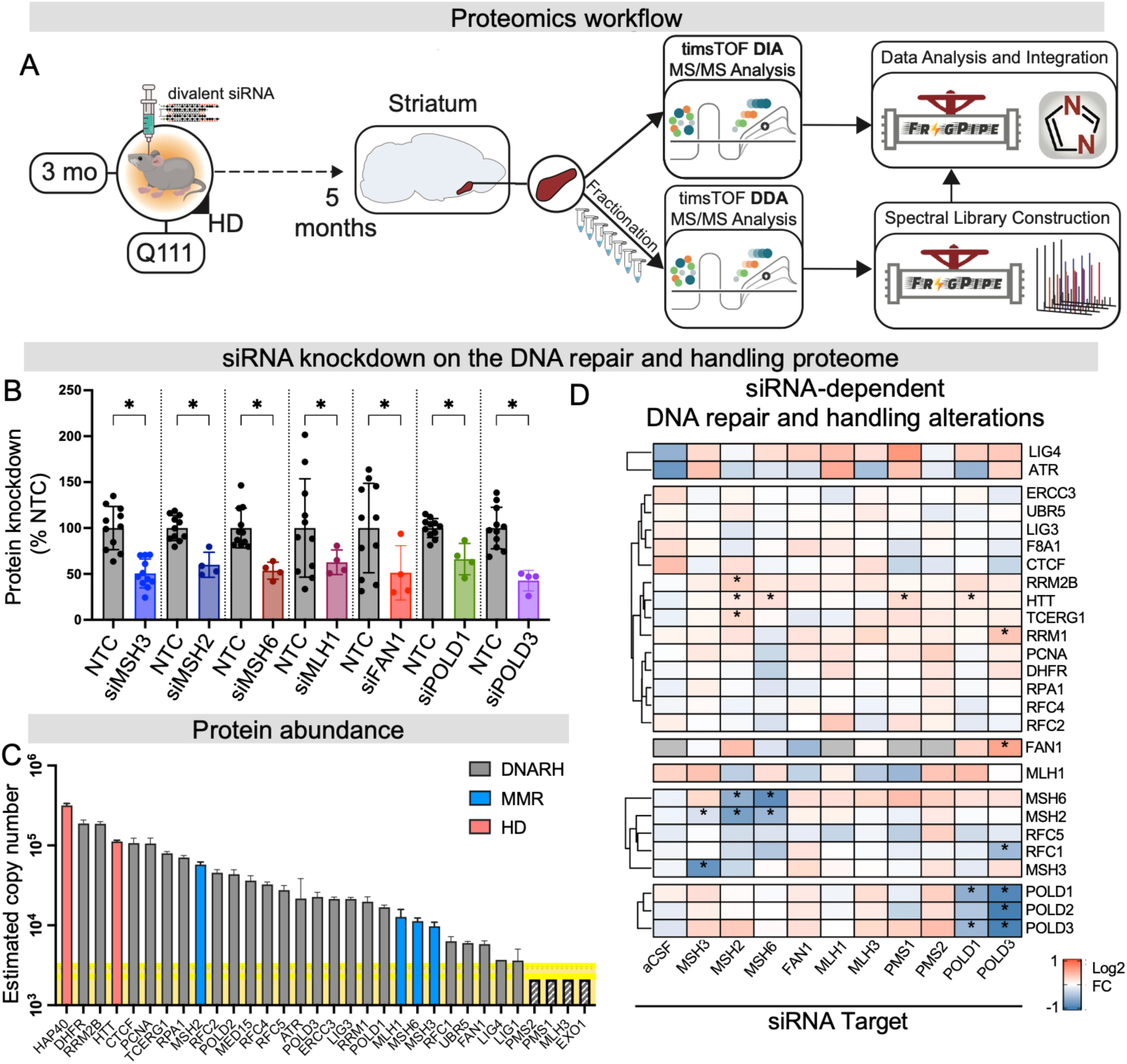
DNA repair pathway proteome response to single MMR target modulation via RNAi. (A) Tissue processing for proteomic analysis and workflow. (B) Single target knockdown validation of each target compared to non-targeting control (NTC). (C) Estimated copy number (Log_10_ scale) of HD related (red), DNA repair and handling (DNARH, gray), or mismatch repair pathway (MMR, blue), calculated by the Histone-based Proteomic Rule approach. Highlighted yellow indicates confidence level of detection confidence interval, with proteins below the level of detection highlighted with striped shading. (D) Heatmap of single target silencing (siRNA target, x-axis) for DNA repair and handling proteins, expressed as log_2_ fold-change, with red and blue shading indicating increased or decreased abundance, respectively, compared to NTC. Significance was indicated by * (adjusted p-value < 0.1).

The experimental library consisted of proteins isolated from unfractionated striatal tissue of control Q111 (NTC) and MMR siRNA treated Q111 mice. NTC samples were present within each treatment group (Figure S12A) and were used in relative quantification of target knockdown and to assess the impact of siRNA treatments on the striatal proteome. Evaluation of variance within siRNA treatment and control groups flagged one (out of 12) of the NTC replicates (NTC_03) for removal from the downstream analysis (Data S2). Principal component analysis (PCA) across siRNA and control conditions confirmed a minor but consistent batch effect (Figure S12B), which could be modeled and mitigated by limma (Materials and Methods) through inclusion as covariates during differential protein analysis.

Aided by the deep mouse spectral library, the majority of siRNA targets were quantified, showing significant knockdown levels (Figure 4B) supporting the RNA and western blotting analyses. For several targets (FAN1, MSH3, MLH1) their absolute abundance and relative knockdown levels were sufficiently robust that the proteins were no longer quantified in most replicates. For example, FAN1 was quantified by a single peptide in six out of 11 controls but only in one out of four targeting siRNA, supporting its knockdown at the protein level. Due to this effect, to more accurately reflect target knockdown levels in Figure 4B, missing values in both control and siRNA groups were imputed. In contrast, we did not use imputation for global proteome analyses (Figures 4C and D, 5 and 6) to avoid imputation-related false discoveries, with the acceptable trade-off that visualization of target knockdown underestimates actual knockdown levels. Interestingly, statistically significant reductions in POLD1 and POLD3 were observed using quantitative proteomics, despite only marginal changes detected by western blot (Figure 4B, Figures S8, 9). In contrast, three MutL targets (PMS1, PMS2, and MLH3) and EXO1 were below the limit of mass spectrometry detection, however, knockdown remains confident based on RNA and western blotting quantification. This highlights a value of using orthogonal methodology in evaluation of target modulation levels.

Next, we performed an independent computational analysis of NTCs using the “proteomic ruler” method to estimate absolute protein copy numbers (Figure 4C, Table S5).^55^ This method estimates absolute protein abundances by scaling MS intensities according to the number of theoretical tryptic peptides,^55^ providing more interpretable quantitative comparisons between different proteins, with the caveat that it has lower accuracy for very large and small proteins, e.g. HTT and HAP40. From the NTC condition (n = 11 biological replicates), 9,464 proteins were quantified with a median CV of 12%, spanning five orders of magnitude in abundance. The 100 least abundant proteins had a median value of ∼2,000 copies per cell, suggesting that the siRNA targets which were not detected (PMS1, PMS2, MLH3, and EXO1) are less abundant (Table S5, Data S2). We rank-ordered the quantified siRNA targets along with proteins that participate in DNA-RH and/or are genetic modifiers of HD (Figure 4C, Data S2 and S3). The proliferating cell nuclear antigen (PCNA) protein was highly expressed (∼10^5^ copies). Moreover, we estimated up to an order of magnitude difference in the amounts of MMR/DNA repair proteins, ranging from 5.84 ± 0.43 x 10^3^ (FAN1) to 5.78 ± 0.37 x 10^4^ (MSH2) copy numbers (Figure 4C). This dataset represents one of the first quantitative characterizations of the relative abundance of mismatch repair (MMR) proteins in the CNS of Huntington’s disease models. The resulting framework facilitates interpretation of the differential dose sensitivity and target-specific contributions of individual MMR factors to somatic instability, providing important context for the development of MMR-directed therapeutic strategies.

### Single-gene MMR silencing produces distinct proteomic signatures in the DNA maintenance network

We next investigated the consequences of single-gene MMR silencing on steady-state proteome abundances (Figure 4D). Based on mRNA, western blot, and MS-based quantification of siRNA targets, we selected the single most potent siRNA treatment for each target for comparison. Non-targeted mass spectrometry analysis quantified similar numbers of proteins (∼8, 200) with uniform variance distributions (∼15% median) among siRNA groups (Figure S13A & B), further supporting the robustness and consistency of protein quantification. Among these proteins, we detected differential protein regulation using DEqMS, which employs well-established linear models (limma) and Bayesian methods (eBayes) with a variance correction based on the number of quantified peptides.^56^ Differential proteins were filtered at FDR < 10% (adjusted p-value < 0.1) and for some comparisons, an effect size threshold was used. Comparison of NTC and artificial CSF (sham) treated animals showed minimal proteomic variation (Figure S13C), indicating that the observed changes following siRNA treatment are not driven by the oligonucleotide scaffold or administration procedure. We have previously established by RNA-seq that siRNA-mediated modulation of MMR targets in the CNS is highly selective.^6^ Notably, long-term MSH3 silencing significantly reduced expression of the intended target while producing minimal off-target transcriptomic perturbations. These data support the conclusion that the proteomic alterations identified here predominantly reflect biologically relevant downstream consequences of target modulation.

Within siRNA targets, at baseline levels, MutSα/β components MSH2, MSH6, and MSH3 had different cellular abundances at 5.8 ± 0.4 x 10^4^, 1.1 ± 0.1 x 10^4^, and 0.98 ± 0.09 x 10^4^ cell copies, respectively (Figure 4D, Data S3). The wide differences in abundance may be mechanistically relevant, as MSH2 is about five times more abundant than MSH3. Upon knockdown, coordinated effects between MutSα/β components were observed. For example, silencing of either MutSα subunits (MSH2 or MSH6), resulted in 40-50% target knockdown (Figure 4B and D) and a similar level of reduction in the other subunit (Figure 4D), consistent with the known role of MSH2 in stabilizing MSH6.^57,58^ In contrast, MSH3 levels remained unchanged following MSH2 silencing, while MSH2 was slightly (10%) but significantly lower with MSH3 silencing.

MSH3 silencing showed a small reduction in MSH2 levels and elevation in MSH6, though neither were statistically significant (Figure 4D). Conversely, MSH6 silencing did not impact MSH3 levels. This suggests that MSH3 is more tightly transcriptionally regulated and that MSH2 availability is not limiting for MutSβ complex formation. In contrast, MSH6 upregulation may outcompete MSH3 for MSH2 binding, potentially limiting MutSβ complex formation, a finding that may open a new therapeutic avenue for modulating MMR pathway activity. Interestingly, in addition to its interconnectivity with the MutS subunits, silencing MSH2 led to upregulation of RRM2B and TCERG1, both of which were previously identified in GWAS studies as modifiers of age of onset, with TCERG1 also reported as an HTT protein interaction^54,59^. This indicates a direct link between MMR locus occupancy, transcriptional elongation, and dNTP production.

The modulation of both MSH2/MSH6 and PMS1 resulted in a statistically significant upregulation of HTT expression by 10 - 17% (Figure 4D, Figure S13D, Data S4). Since HTT expression decreases as the level of expansion increases,^60^ this may serve as a readout of reduced expansion rates. However, realistically, over two months the degree of expansion though measurable is unlikely sufficient to drive a change in HTT expression levels. In addition, MSH6 silencing produces a similar effect but does not materially alter expansion. Thus, this relationship is more likely to reflect the impact of the MMR machinery occupancy at the locus on transcriptional elongation. MLH1 silencing eliminated MLH1 protein signal in all but one replicate, but did not significantly affect levels of other measurable MMR partners (Figure 4D), a surprising result given that MLH1 is the core component of MutL complexes and prior findings that in silico mutations to MLH1 can disrupt PMS1 and PMS2 levels.^61^ This may suggest alternative interaction models or compensatory mechanisms to the *in vivo* MMR complex interactions. FAN1 knockdown did not have overt effects on other DNA-RH partners (Figure 4D), though a mild 15% up-regulation across the panel was observed (Data S3).

To assess whether our PMS1 knockdown findings were plausible, we compared differentially expressed (DE) proteins in striatum of Q111 PMS1 siRNA-treated mice (5m) (Data S3) with DE genes from striatum of PMS1 homozygous knockout Q140 mice (6m) reported in Table S1 of Wang et al.^8^ Overall, at the same FDR threshold, the total numbers of DE proteins/genes was in a similar range (1663 genes vs. 1274 proteins, Figure S14A). Moreover, for the 150 shared targets between the two datasets, STRING network analysis revealed significant connectivity (92 proteins). It is notable that alterations of transcripts and proteins due to PMS1 silencing both connect to prominent pathways pathways involved in HD pathobiology, supported by the largest cluster in the network (Figure S14B, red nodes) containing HTT and other proteins involved in synaptic transmission and cytoskeletal organization. Overall this analysis supports a conserved role for PMS1 in modulating the HD transcriptional landscape across independent models and modalities (Figure S14B).

POLD1- and POLD3-targeting siRNAs shows high interconnectedness within the transcriptional machinery confirming current understanding of the polymerase-multi-subunit complex involvements (Figure 4D).^9,15,21,41,62^ POLD3 silencing strongly decreased the levels of POLD1 (p < 0.001), POLD2 (p < 0.001), and RFC1 (p < 0.01). POLD3 silencing also significantly increased FAN1 a protein associated with reduced expansion rates.^10–13,40,51,63^ This is again likely related to the link between transcriptional elongation and MMR locus occupancy. It also raises an interesting question as to whether the negative impact of POLD3 silencing on expansion is a direct effect or is mediated indirectly through FAN1 upregulation.

Silencing of POLD3 increases Rrm1, a key player in dNTP production.^64^ It is unclear whether this is a direct effect or a consequence of POLD3 silencing or related to positive feedback to enable nucleotide availability.

Together, these observations suggest a functional connection between FAN1-based protection against expansion, transcriptional regulation and genome stability, potentially explaining the impact of POLD3 silencing on reduced somatic instability (Figure 3F, G).

Overall, this analysis provides the first systematic proteomic dataset characterizing siRNA-mediated MMR perturbation and its impact on the mouse DNA repair proteome. These data offer new insights into potential functional protein–protein interactions and regulatory hierarchies within the pathway.

### MSH3 silencing minimally affects the global proteome, while PMS1, MLH1, and MLH3 silencing induces broad proteomics remodeling

Over fifty repeat-associated disorders involve somatic instability as a key driver.^65–67^ In HD models, targeting somatic expansion reduces protein aggregates and reverses neurodegenerative transcriptional signatures.^6,8,33^ These findings establish the MMR pathway as a compelling therapeutic target.^7,9,10,12^ Nevertheless, MMR is essential for maintaining genomic integrity. Therefore, safety remains a critical consideration for therapeutic modulation. Here, our DIA-PASEF-based quantitative MS dataset enables sensitive and unbiased evaluation of divalent siRNA-induced effects on the tissue-level proteome. This approach captures both direct on-target and downstream network-level changes at depth and breadth.

We found differences in proteome remodeling depending on the silenced MMR component (Figure 5, Figure 6). To break down these changes, we first examined the impact of single-target knockdown on the global proteome (Figure 5A-J, Data S2, Data S3) and then characterized patterns between the knockdown proteome signatures and if these signatures aligned with shared or unique biological pathways (Figure 6). MSH3 and PMS1 have emerged as top therapeutic candidates based on their dose-sensitive effects on somatic instability. We compared their impact on proteome remodeling in striata from Q111 mice treated with siRNAs targeting either MSH3 or PMS1. Remarkably, MSH3 silencing, despite fully blocking somatic expansion (Figure 2D), caused minimal disruption of the global proteome (Figure 5A,K). Very few proteins exhibited >1.5-fold changes (Figure 5A), with only one protein, PLIN4 (p < 0.05), showing statistically significant upregulation. PLIN4 is involved in lipid storage and turnover and contains ∼30 repeats of a 99-bp region prone to expansion, raising the possibility of a mechanistic link between MSH3 silencing and PLIN4 regulation.^68,69^ Importantly, the same MSH3 siRNAs administered for 10 months in wild-type mice induced no significant transcriptomic changes, supporting both the specificity of the target and the inertness of the siRNA sequence in the fully modified scaffold.^6^

**Figure 5.**
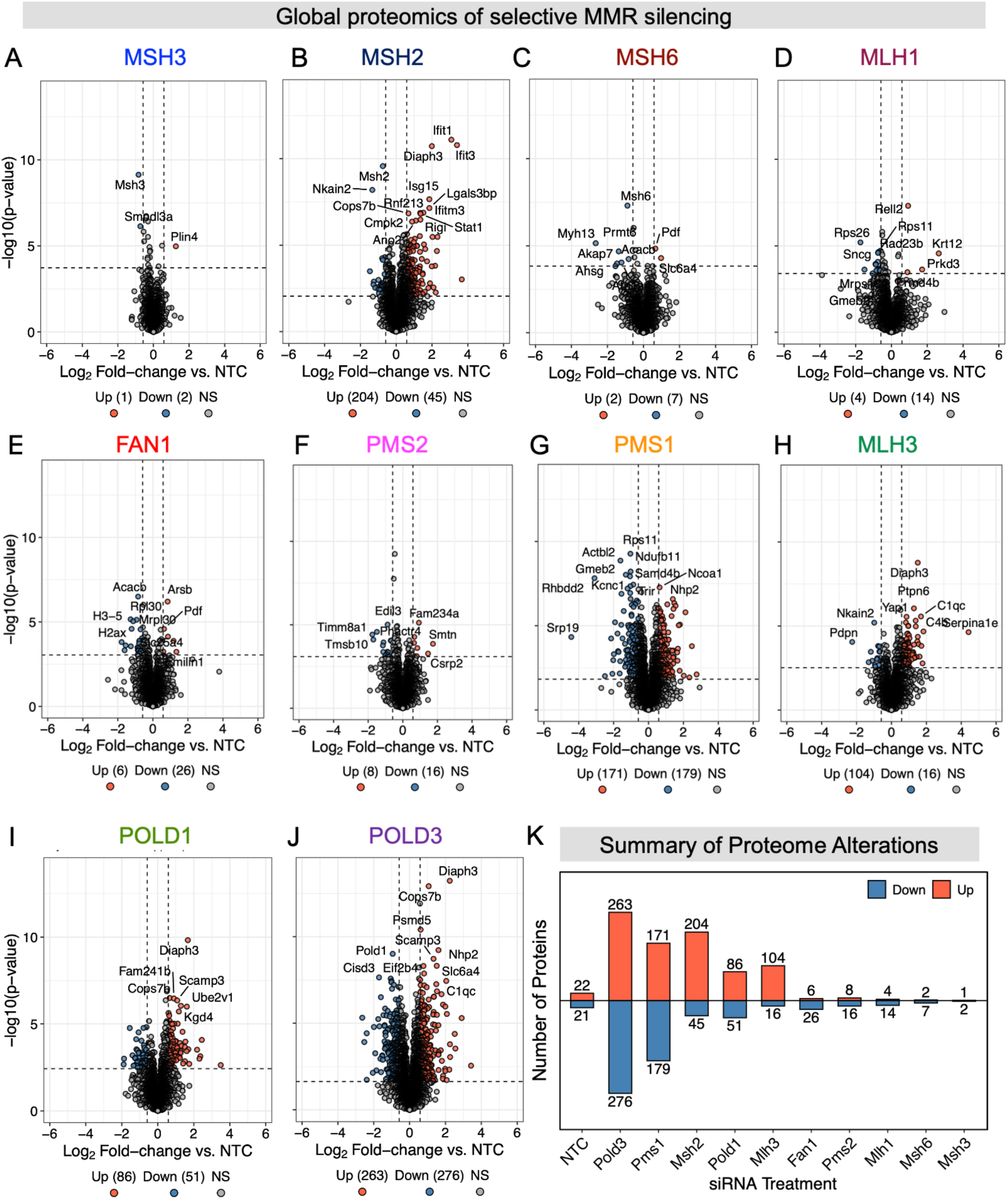
MSH3 silencing has minimal effect on global proteome. Volcano plot of proteins within the DNA repair pathway levels following siRNA silencing of (A) MSH3 (B) MSH2 (C) MSH6 (D) MLH1 (E) FAN1 (F) PMS2 (G) PMS1 (H) MLH3 (I) POLD1 or (J) POLD3. Significant proteins and thresholds are indicated in red and blue circles. X-axis is log_2_ fold change versus NTC. Y-axis is the −log_10_(p-value) of each protein. Horizontal threshold reflects p-value at FDR = 0.1. (K) Number of proteins differentially expressed across the global proteome with respective siRNA treatment. Red and blue are up and down-regulated groups, respectively.

**Figure 6.**
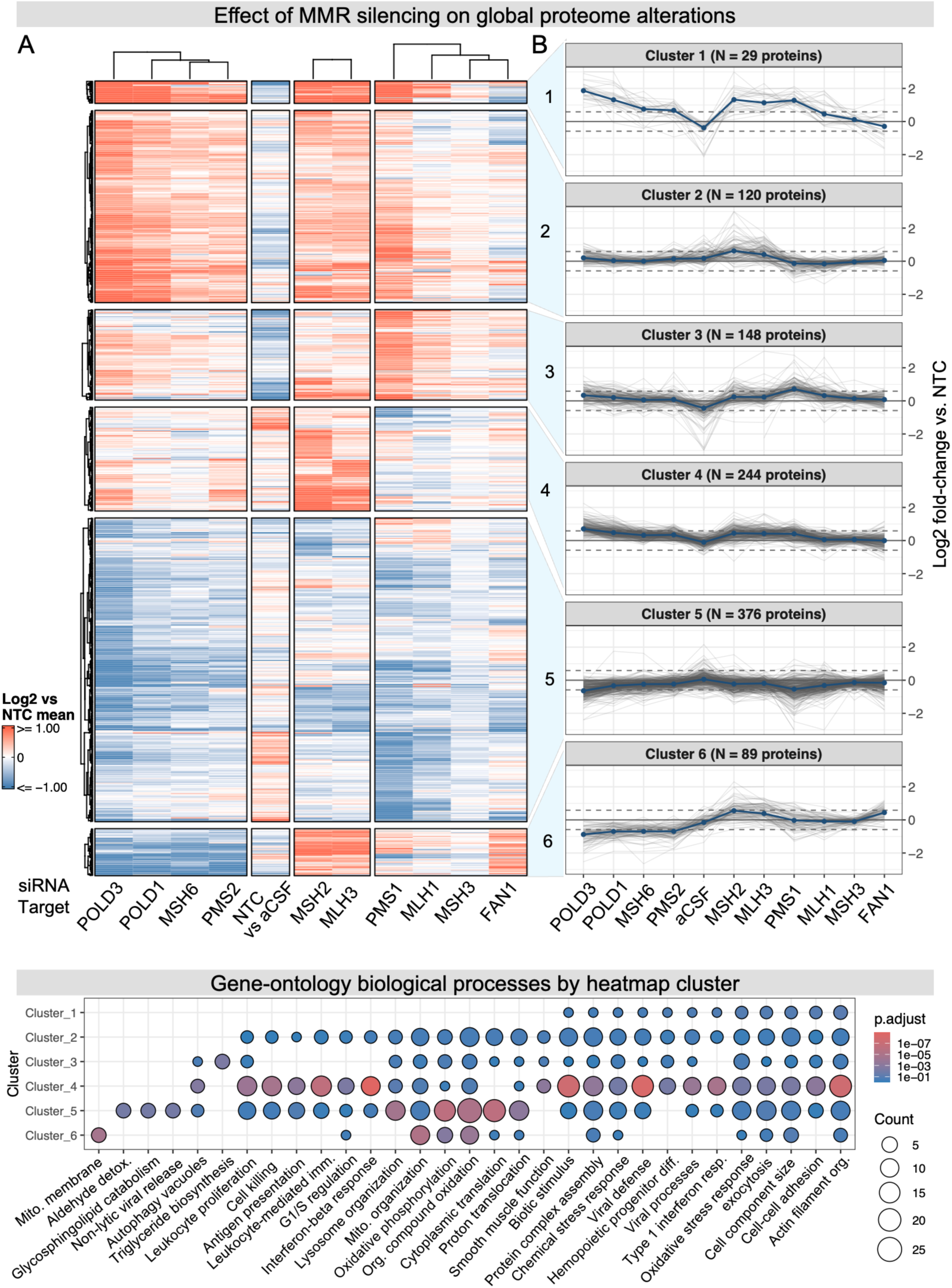
Silencing MMR pathway implicates mitochondrial and immune response biological pathways. (A) Heatmap of global proteome changes following single MMR target silencing. Proteins that were differentially regulated in at least one silenced target group were clustered by their log2 fold-change versus NTC. Note, the effect of NTC treatment on these proteins was measured versus aCSF and included for reference to the target ratios (B) Spaghetti cluster profile plots showing the median log2 fold-changes of the cluster across all siRNA treatments. Data used to generate graph is shown is Data S4. (C) Gene ontology over representation analysis of biological processes (BP) by heatmap cluster. Color indicates adjusted p-value. BP terms were significant (p < 0.01) in at least one cluster; non-significant clusters with annotated proteins were shown for transparency. Size of the circle indicates the number of proteins associated with the designated pathway.

PMS1 silencing, which showed a similar blocked expansion phenotype, triggered a greater degree of proteome remodeling, with 350 proteins having abundance differences, mostly in the range of 1.5 – 2-fold (Figure 5G, PMS1). PMS1, along with MLH1 and PMS2, has been shown to support a large interactome in whole-cell extracts.^70^ These results highlight the potential involvement of PMS1 in diverse biological functions, consistent with its involvement in DNA repair, transcription, and chromatin architecture.

MLH3 silencing led to alteration of 116 proteins, the majority of which were up-regulated (n = 100) (Figure 5F, MLH3). These protein alterations pointed to cellular factors involved in chromatin organization/DNA-binding and Stat1-mediated interferon signaling. Interestingly, MLH3 knockdown induced strong upregulation of interferon-inducible GTPases (e.g., IRGM2, IGTP, IIGP1).

Overall, the effect of single MMR modulation on proteome dysregulation was target-dependent (Figure 5A-H), with POLD3 and PMS1 producing the broadest changes and MSH3 the fewest (Figure 5I).

### Biological characterization of global proteome changes associated with silencing of individual MMR members

We next asked whether there was any commonality in the proteome signatures and pathways affected by selective modulation of MMR components. Overall, 1016 proteins were differentially regulated (|log2FC| > 0.585, FDR < 10%) following silencing of at least one MMR component (Data S4). Given the average number of differential proteins per MMR target was ∼140, the relatively large number of unique differential proteins (n=1016) suggests that each target knockdown may induce uniquely pleiotropic effects. Further, while the siRNA constructs were designed to minimize off-target effects,^43^ off-target effects of siRNA are a potential confounding factor. Hierarchical clustering of log2FC values from the differential proteins was performed to identify coordinated and divergent proteome alterations (Figure 6A,B, Data S4). The signature of the non-targeting construct (NTC) vs. control (aCSF) is unique and largely opposing that of the other target siRNA vs. control (NTC). As expected, POLD3 and POLD1 differential signatures were highly similar. MSH6 and PMS2 silencing shared similar patterns as POLD silencing. Interestingly, MSH6 and PMS2 both function to counter somatic expansion. This shared signature again points to strong interconnectivity between transcriptional elongation and MMR-mediated repeat regulation and strongly supports that the observed proteomics characteristics are functionally relevant rather than driven by off-target effects of the individual sequences.

To explore the specific pathways that underlie these differential signatures, the average differential patterns from six row clusters were assembled (Figure 6B) and the protein clusters were tested for significant enrichment in biological processes versus the genome-level GO annotation (Figure 6C, Data S5). The largest cluster, cluster 5, (n = 406 proteins) represented proteins that were largely downregulated across all target siRNAs, with varying magnitudes, while cluster 6 reflected shared downregulation from POLD/MSH6/PMS2 silencing but opposing upregulation under MSH2/MLH3 silencing. This dichotomy is interesting as the latter behaved similar to MSH6 and PMS2 with respect to expansion, but have opposite proteome regulation within cluster 6, which includes strong enrichment for mitochondria-related processes (Figure 6C). FAN1 silencing had a similar upregulation effect within cluster, even though it has the opposite impact on somatic instability.

MSH2 and MLH3 silencing formed a distinct column cluster, with proteome upregulation reflected most strongly in cluster 4, which was functionally enriched in immune responses processes (Figure 6C and Data S5). The relative proteome quiescence of MSH3 and FAN1 silencing likely drives their similar clustering, however, MSH3’s pattern also shares features with PMS1 silencing, which is interesting given PMS1 and MSH3 are on opposite ends of the differential spectrum (see Figure 5K). Evaluating PMS1 in more detail, we observed that cluster 2 contained proteins up-regulated by PMS1 silencing, but this signature was also shared with most of the other MMR components. In contrast, the proteins within cluster 3 (N = 121 proteins, Figure S15) were most strongly upregulated in PMS1 silencing. Given the low number of cluster members, none of the functional terms were strongly enriched, though oxidative phosphorylation, cell stress responses, and actin cytoskeleton organization were the most prominent (Figure 6C, Figure S15, Data S5).

Two major processes emerge from the proteomics analysis of MMR silencing: mitochondrial dysfunction and response to oxidative stress, with specific pathways showing both up- and downregulation depending on the silenced targets, and immune system modulation or stability. These prominent mitochondrial signals have been described in HD for decades and may reflect not only the presence of mutant HTT but also changes in MMR, dNTP availability, and transcriptional machinery.^71–73^ Another highly significant but selectively enriched cellular process associated with upregulation under MSH2/MLH1 silencing, involved in various immune responses, such as interferon stimulation and regulation of leukocytes. Together, these findings provide an added dimension to our understanding of the interplay between MMR, transcription, mitochondrial and immune system response to stress.

Collectively, these data show that while multiple MMR components suppress somatic expansion, MSH3 uniquely does so with minimal off-target effects, making it a leading therapeutic candidate for HD and other repeat expansion disorders. More broadly, our findings uncover a stoichiometry-sensitive proteomic network linking DNA repair, replication, transcription and mitochondrial function, where even modest MMR knockdown disrupts distant partners. This dosage sensitivity may underlie striatal neuron vulnerability in HD and underscores the need for system-wide evaluation in therapy development.

## DISCUSSION

We investigated the MMR pathway *in vivo* using divalent siRNAs to modulate somatic repeat expansion in the CNS of an HD mouse model. Therapeutic silencing of MSH3, MSH2, PMS1, MLH1, MLH3, and POLD3 effectively blocked or slowed expansion, whereas knockdown of FAN1 and PMS2 accelerated it. Proteomic analyses revealed key interdependencies within the DNA repair network, including the impact of MSH2 knockdown on the HD modifiers TCERG1 and RRM2B, interconnectedness between the MMR and transcription machinery, and MMR single-target modulation on mitochondrial stress and immune system reactivity. Importantly, MSH3 knockdown left a minimal proteomic footprint despite its potent suppression of somatic expansion, supporting it as a high-priority therapeutic candidate. More broadly, this work expands the framework for manipulating somatic expansion and provides new insights into MMR network dynamics in post-mitotic mammalian tissue.

In HD, the selective vulnerability of medium spiny neurons and Purkinje cells to somatic expansion^7,41^ is thought to stem from the absolute and relative expression of MMR components—particularly MSH3, MSH2, MSH6, MLH1, and FAN1—as well as the local chromatin structure and transcriptional activity at the *HTT* locus. Here, dose sensitivity of mismatch repair components revealed mechanisms of CAG repeat expansion and therapeutic target feasibility in HD. Divalent siRNAs achieved 80–90% knockdown of core MMR machinery (MutSα and β and MutLα, β, and γ), enabling systematic assessment of dose sensitivity and therapeutic feasibility. Our finding that MSH3 knockdown robustly blocks expansion while comparable knockdown of MSH2 only partially reduces it likely reflects higher baseline MSH2 expression: even with reduced MSH2 levels, sufficient MutSβ can still form, whereas reducing the lower-abundance MSH3 allows MSH6 to outcompete it for MSH2 binding, disrupting MutSβ formation. This result contrasts with earlier data from MSH2 knockout cell lines, where MSH3 protein was reduced despite stable mRNA levels, implicating post-translational destabilization.^74^ Importantly, prior work from the Yang lab has shown that heterozygous knockout of MSH2 or MLH1 can enable residual expansion to occur, supporting that protein abundance,^8^ even minor, can influence the entire system dynamic. MSH6 silencing trended toward increased expansion, consistent with competition between MutSα and MutSβ. Collectively these findings emphasize the importance of studying biological processes in both genetic and interventional paradigms, specifically in biological systems where ratio interactions may be functionally significant.

Dose sensitivity was also observed for PMS2. Whereas a 50% reduction (with PMS2_600 siRNA) resulted in a slight, non-significant increase in expansion, an 80% reduction (with PMS2_1041 siRNA) significantly increased somatic instability, suggesting the stabilizing impact of PMS2 may not be region-specific and that varying PMS2 expression levels may contribute to regional vulnerability to expansion. Interestingly, the protective role of PMS2 has been masked in different cellular contexts.^15^ For example, PMS2 knockout reduces expansion in yeast and induced pluripotent stem cells. CRISPRi studies further demonstrate that the mouse liver is significantly less sensitive to PMS2 silencing compared to the brain,^21^ underscoring the importance of cellular context. Our finding that PMS2 knockdown increases expansion in the medial cortex suggests that PMS2-targeting siRNAs could serve as a tool to accelerate expansion in regions with inherently slower rates in mouse models.

Somatic repeat expansion is a rare event that depends on transcription and is limited to non-dividing cells. The average initial expansion rate in humans is approximately one CAG repeat per year, with high selectivity for medium spiny neurons and progenitor cells.^7,41^. While the exact number of HTT transcriptional events per day is not well defined, estimates range from 2 to 50 based on observable mRNAs, suggesting only one in 500–2,000 transcriptional events results in productive CAG expansion. Slippage likely occurs more frequently, but the efficiency of MMR recognition and processing of slipped loops constrains the expansion rate. Our data support that MMR proteins function as potent positive and negative regulators of expansion, reinforcing their multimodal role in repeat expansion disorders.^75^ Silencing PMS2 had the most profound accelerating impact, suggesting it competes with other complexes by binding slipped loops in a dominant but non-expansionary manner. The ratio of MMR machinery to HTT transcriptional events may determine selective cellular vulnerability.

These data support multiple models of somatic repeat expansion. The primary event driving somatic expansion is likely the recognition of slipped CAG loop structures, generated during *HTT* transcription, by the MutSβ sliding clamp complex.^24,27^ Our data strongly support this model, with MSH3 acting as the concentration-limiting factor. The significant impact of MLH1, MLH3, PMS1, and PMS2 modulation further supports downstream recruitment of MutL complexes^76–78^. FAN1 acts protectively against expansion, consistent with prior findings^10–13,40^.

The pathway of MutL recruitment following MutSβ binding is less clear. MutLα, the predominant form in MMR, is highly protective against expansion. MutLγ also has nuclease activity^17^, while MutLβ (MLH1-PMS1 in humans, MLH1-MLH2 in yeast)^79^ lacks known nuclease activity. In this case, one could speculate that somatic expansion occurs only when the slippage loop is recognized by a MutSβ that then recruits a complex containing both MutLβ and MutLγ.^37^ The higher affinity of MutLβ may be necessary to increase the complex’s half-life,^80,81^ supporting productive MutLγ-induced cleavage and expansion. Indeed, MutLγ has been shown to preferentially cleave opposite the slipped loop, whereas MutLα can cleave on both strands, though it has been shown to engage in strand discrimination depending on nicks^17,77,82^. Our finding that PMS1 knockdown suppresses expansion suggests that MutLβ is essential in facilitating expansion. Without a nuclease domain, one could speculate PMS1 functions to coordinate and position the nuclease-competent MutL complexes to optimize strand cleavage. PMS2 may act similarly to FAN1, competing with PMS1 and MLH3 to bind MLH1 and prevent MutLγ and MutLβ formation.^38^ While baseline levels of PMS2 are relatively low, the profound stabilization effect of PMS2 on repeats observed in this study indicates high-affinity binding and preferential complex formation, supporting this competition model.

Nevertheless, two key questions remain unresolved: (1) Why do both MutLβ and MutLγ play a role in productive expansion? This hypothesized interdependence is supported by RNAi-mediated, splice modulator, and genetic CRISPRi experiments.^15,19,21^ (2) Why does the absence of MutLα profoundly enhance expansion rates? Based on our results, we propose three potential models: the Proximity Model, the Multi-Subunit Filament Model, and the Holliday Junction Model (Figure 7).

**Figure 7.**
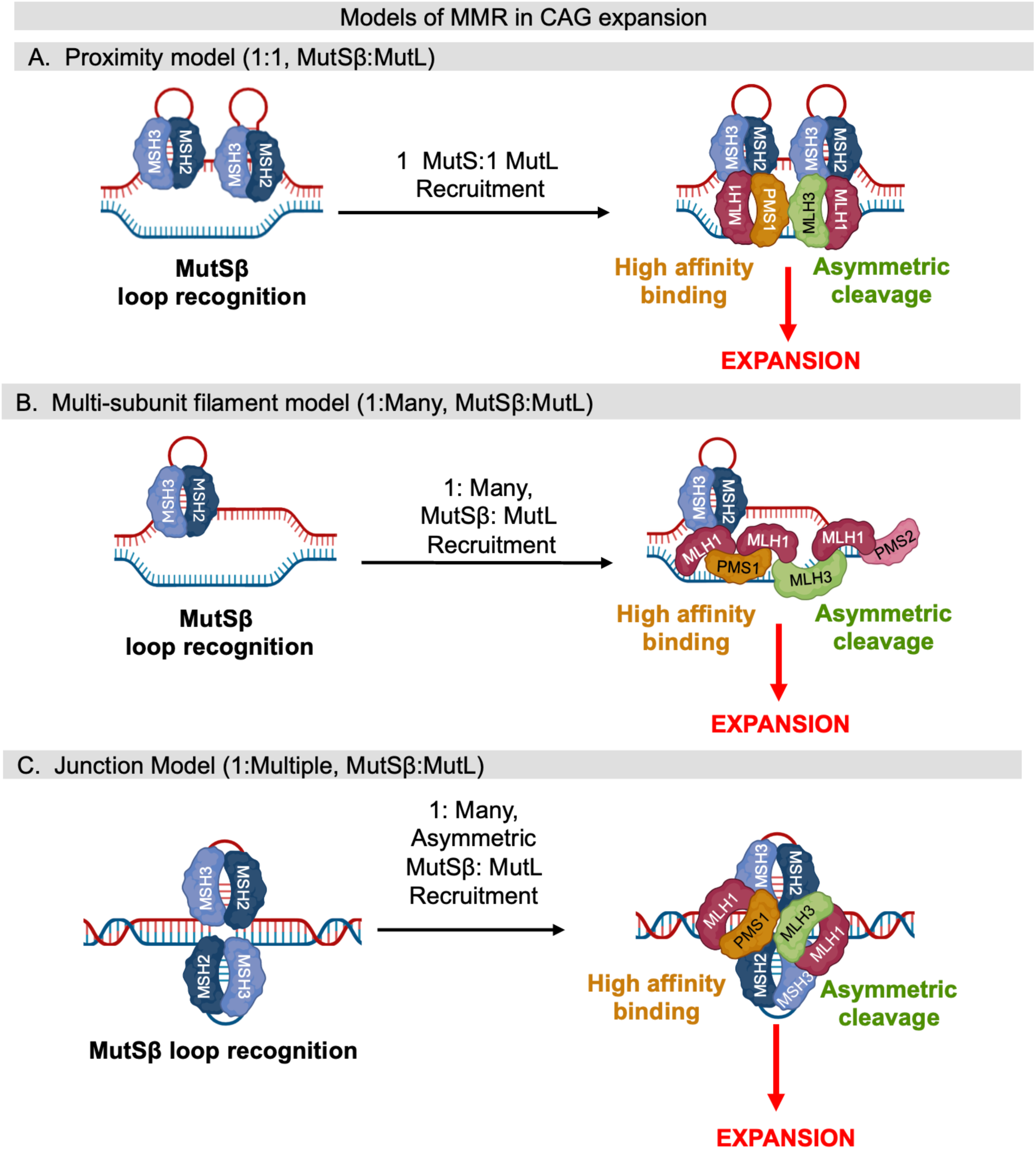
Potential models of MMR involvement in somatic CAG expansion. (A) Proximity model: MutSβ recruits a single MutL complex. When MutLβ and MutLγ are in proximity, PMS1 can serve as a high-affinity anchor to coordinate MLH3 to cleave a single strand of DNA, causing expansion. In PMS1 knockdown conditions, there is no MutLβ to anchor the nuclease-compatible MutL complexes to guide positioning for cleavage. In MLH3 knockdown conditions, no single-stranded DNA cleavage can take place. In PMS2 knockdown conditions, MutLγ and MutLβ form in higher concentrations, as PMS1 and MLH3 outcompete PMS2 for MLH1. (B) Multi-subunit filament model: one MutSβ recruits multiple MutL complexes in a chain. MutLβ and MutLγ are both required to anchor (Pms1-dependent) and cleave (Mlh3-dependent). In PMS2 knockdown conditions, MutLγ and MutLβ form at higher concentrations, promoting expansion. (C) Holliday junction model: MutLγ and MutLβ both bind a slip loop at orthogonal positions, inducing stabilization by MutLβ and cleavage by MutLγ.

In the Proximity Model (Figure 7A), the slipped loop is recognized by MutSβ, which recruits a single MutL complex (MutLα, MutLβ, or MutLγ). For expansion to occur, two MutSβ complexes must be in proximity, either on a single slipped loop or on adjacent loops, each independently recruiting a separate MutL complex. Once bound, MutLβ (PMS1) coordinates the complex while MutLγ (MLH3) cleaves the strand opposite the slipped loop. In the absence of PMS1, MLH3 coordination fails; without MLH3, no cleavage occurs; and without PMS2, MutLβ and MutLγ can complex more frequently, increasing expansion probability.

In the Multi-Subunit Filament Model (Figure 7B), MutSβ recruits multiple MutL complexes that form filaments or a multi-subunit complex. MLH3 performs single-stranded nicks, with PMS1 coordinating MutL positioning and PMS2 regulating expansion by limiting MutLγ or MutLβ formation. This model explains the necessity of both MutLβ and MutLγ due to their distinct interaction interfaces and filament formation. This one-to-many interaction is emerging as a common model to explain human MMR complex interactions in the context of repeat expansion disorders.^83,84^

In the Holliday Junction Model (Figure 7C), MutLβ and MutLγ resolve Holliday junctions, as previously described^85–87^. In yeast, these complexes primarily function in resolving meiotic junctions, which structurally resemble CAG-CTG repeat slipouts. This resemblance may explain the requirement of both MutLβ and MutLγ in somatic expansion. PMS2 does not participate in Holliday junction resolution, aligning with its protective role against expansion.

The Multi-Subunit Filament or Holliday Models align with our observation of a rate-limiting dependence on MutSβ for complex formation and expansion. Further, both models incorporate a MutLβ and MutLγ as a key component. The Holliday junction model is further supported by independent genetic validation of the co-requirement of PMS1 and MLH3. ^85,86,88,89^ The Proximity Model suggests equal dependence on multiple factors, which is inconsistent with our study findings. Future combinatorial studies will be essential for refining these models, particularly in neurons, where cellular context significantly influences the role of PMS2. For example, our finding that PMS2 knockdown profoundly enhance the rate of expansion highlights the protective role of the MutLα complex. Promoting formation of this complex, perhaps by simultaneously targeting MLH3 and PMS1, may enhance protection against expansion or even promote contractions. Co-silencing two genes in the CNS is feasible using a unimolecular dual-targeting divalent siRNA scaffold.^90^ Gaining a deeper understanding of MMR mechanisms will provide critical insights into the molecular processes underlying somatic repeat expansion.

Multiple lines of evidence support blocking somatic expansion as a disease-modifying therapy for HD,^6,8,33,91^ yet much work is needed to translate MMR modulation from preclinical models to the clinic. In HD models, reducing MSH3 protein via genetic knockout, antisense oligonucleotides, or divalent siRNAs reduces mutant HTT inclusions, reverses transcriptional dysregulation, and improves neurodegenerative signatures, with greater effects when combined with HTT-lowering strategies^6,8,33,92^. Both MSH3 and PMS1 emerged as effective targets for blocking somatic expansion. However, PMS1 silencing induced substantially greater proteomic changes than MSH3 silencing and was associated with alterations in a larger number of MMR-related proteins, suggesting a broader involvement in multiple cellular pathways. Additional studies will be required to fully evaluate the clinical safety of PMS1 modulation, particularly given the growing interest in this target for therapeutic development. Importantly, changes in protein abundance alone do not inherently indicate either toxicity or safety and therefore require careful biological interpretation and follow-up investigation. Recent GWAS analyses have not identified deleterious safety signals associated with PMS1 variants (Qingqin Li, personal communication, CHDI Conference 2026, Palm Springs), providing preliminary support for the feasibility of this approach.

A limitation of the current analysis is that only a single PMS1-targeting siRNA was evaluated in the proteomic studies, constraining interpretation of target-specific effects. Furthermore, the extent to which PMS1 reduction influences the abundance or activity of interacting MMR components, including PMS2 and MLH3, which could not be reliably quantified in the present dataset, remains unknown and warrants further investigation.

Targeting MMR via divalent siRNA could potentially be applied to disorders beyond HD, such as Myotonic dystrophy type 1 (DM1), Spinocerebellar ataxia (SCA), Spinal and bulbar muscular atrophy (SBMA), and Dentatorubral–pallidoluysian atrophy (DRPLA) where somatic instability is also driven by MMR-related variants,^93–100^ highlighting a potentially common MMR-driven mechanism. Among siRNA targets tested, only MSH3 shows therapeutic efficacy without known cancer associations in humans or mice.^101–109^

While the available preclinical and human genetic data are generally encouraging, therapeutic modulation of mismatch repair (MMR) pathways should be approached cautiously. MMR proteins play fundamental roles in maintaining genome integrity, and the long-term consequences of partial inhibition may not be fully captured by proteomic analyses, animal studies, or currently available human genetic datasets. As a result, restricting MMR modulation to therapeutically relevant tissues, such as the CNS, may provide an important additional safety margin.

Although intracerebroventricular administration of divalent siRNAs substantially minimizes systemic exposure, low levels of target knockdown have been observed in peripheral tissues, including the liver, as a consequence of CSF clearance and systemic redistribution. This observation highlights the potential value of incorporating liver-blocking strategies, such as GalNAc-conjugated antisense oligonucleotides that selectively inhibit siRNA activity in hepatocytes (ref), thereby reducing peripheral MMR modulation while preserving therapeutic activity within the CNS. Such approaches may become increasingly relevant as multiple CNS-directed siRNA platforms, including divalent, PS-extended, and lipophilic-conjugated scaffolds, advance toward clinical development. Notably, one divalent siRNA program has already entered human clinical evaluation under an open IND, supporting the translational feasibility of long-term MMR modulation in the CNS.^34,110–113^ New delivery strategies using transferrin receptor-targeting antibodies show early promise but may require further safety evaluation due to systemic exposure.^111,114^ Safe CNS administration also depends on calcium and magnesium co-dosing to prevent acute neurotoxicity.^49^ Moving forward, biomarker development, both for MSH3 knockdown and noninvasive measurement of somatic expansion,^115,116^ will be critical for translating these therapies into clinical trials and expanding their relevance across repeat expansion disorders.

## Methods

### Oligonucleotide synthesis

Oligonucleotides were synthesized using a solid-phase synthesis method with phosphoramidites on Dr Oligo 48 (Biolytic, Fremont, CA) or MerMade12 (Biosearch Technologies, Novato, CA) automated synthesizers. Phosphoramidites used were 5’-(E)-Vinyl tetraphosphonate (pivaloyloxymethyl) 2’-O-methyl-uridine 3’-CE for 5’-Vinyl Phosphonate inclusion as well as 2’F and 2’OMe with standard protecting groups. Phosphoramidites were brought into solution at 0.1M with anhydrous acetonitrile (ACN) with the addition of 15% dimethylformamide for 2’-OMe-uridine. All phosphoramidites were purchased from Chemgenes (Wilmington, MA) and Hongene Biotech (Union City, CA). Detritylation was performed on column using 3% trichloroacetic acid in dichloromethane. The activator used was 0.25M 5-(Benzylthio)-1H-tetrazole (BTT), and the coupling times were 4 minutes. Capping reagents include CAP A (20% N-methylimidazole in ACN) and CAP B (20% acetic anhydride and 30% 2,6-lutidine in ACN). Oxidation or sulfurization reagents used were 0.05M iodine in pyridine and water (9:1, v/v) or 0.1M 3-(dimethylaminomethylene)amino-3H-1,2,4-dithiazole-5-thione (DDTT) in pyridine for 3 minutes. All reagents were purchased from Chemgenes. Oligonucleotides used for *in vivo* and *in vitro* experiments were grown on long-chain alkyl amine (LCAA) controlled pore glass (CPG). The oligonucleotides were functionalized with a succinyl linker with 500Ä Unylinker terminus (Chemgenes) for unconjugated strands, and 500Ä di-trityl protected support connected by tetraethylene glycol linker (Hongene) for the divalent and cholesterol strands through a tetraethylene glycol linker (Chemgenes).

### Cleavage, deprotection, and purification of oligonucleotides used for in vitro experimentation

All oligonucleotides were cleaved and deprotected on-column using ammonia gas (Airgas Specialty Gases). Columns were pre-wet with 100µL of water and spun by centrifugation to remove the excess water. After excess water was removed, the columns were placed into a heated pressure chamber (Biolytic) for 90 mins at 65°C. Desalting and counterion exchange were performed using a modified on-column ethanol precipitation method consisting of subsequent rinses with 1 mL of 0.1M sodium acetate in 80% ethanol, followed by 1 mL of 80% ethanol. Excess ethanol was dried from the columns and the oligonucleotides were eluted off the columns with 600 µL of water.

### Cleavage, deprotection, and purification of oligonucleotides used for in vivo experimentation

Oligonucleotides containing 5’-(E)-Vinyl phosphonate were cleaved and deprotected using a solution of 3% diethylamine in ammonium hydroxide for 20 h at 35°C with mild shaking. Divalent oligonucleotides underwent 1:1 40% aqueous monomethylamine with ammonium hydroxide for 2 h at 25°C with mild shaking. After cleavage and deprotection, both oligonucleotide solutions were filtered from the controlled pore glass, rinsed with 5% ACN in water, and dried overnight by centrifugal vacuum concentration. Oligonucleotides were purified using an Agilent 1290 Infinity II HPLC system with Source 15Q anion exchange resin (Cytiva). Mobile phase A was 20 mM sodium acetate with 10% ACN in water, and mobile phase B was mobile phase A with 1M sodium bromide, mixed thoroughly. For all oligonucleotides, a linear gradient of 30 to 70% in 30 mins at 50°C was used, and peaks were monitored at 260nm. Pure fractions were combined and desalted using water in an isocratic size exclusion chromatography method with Sephadex G-25 resin (Cytiva). After salt separation, pure oligonucleotides were dried overnight by centrifugal vacuum concentration and resuspended in water. Fractions and pure oligonucleotide purity and identification were confirmed using IP-RP HPLC on an Agilent 6530 Accurate-mass Q-TOF.

### In vitro screening

HeLa or N2A cells were cultured at 37°C and 5% CO_2_. Duplexed siRNA conjugated with cholesterol for passive cell uptake was diluted into Optimem (ThermoFisher Scientific Cat. 31985070) at a concentration of 3 µM and plated 1:1 with DMEM (Corning, 10-013-CV) with 6% Fetal Bovine Serum (FBS) (Corning, 30-011-CV) (HeLa cells) or EMEM (ATCC, 30-2003) with 6% FBS (Corning, 30-011-CV) for a final siRNA concentration of 1.5 µM. Ten thousand (10,000) cells/well were plated onto the siRNA-containing media in a 96-well plate. Seventy-two (72) hours after plating, media was removed, and cells were flash-frozen for downstream processing. For dose-response experiments, the doses used were 1.5, 0.75, 0.38, 0.19, 0.09, 0.05, 0.02 µM siRNA or untreated. NTC was used as a scramble siRNA control.

### Animal Use

All animals were used in accordance with the Institutional Animal Care and Use Committee at the University of Massachusetts Chan Medical School protocol 202000010 and followed the National Institutes of Health Guide for the Care and Use of Laboratory Animals. Animals were housed and cared for at the pathogen-free facilities at UMass Chan with a 12h light-dark cycle. The facilities were maintained at 23 ± 1°C and 50 ± 20% humidity with free access to food and water. Before arriving at the UMass Chan facilities, the animals were bred and born at Jackson Laboratories (JAX), Farmington, CT. The animals used for all studies were Hdh^Q111^ Het mice, JAX Strain ID 370624, Mc_Q111 KI; C57BL/6J background, heterozygous. The starting CAG tail length was genotyped and provided by JAX for all experiments. An equal number of males and females were used for each study.

### In vivo intracerebroventricular injections

*In vivo* grade di-valent siRNA were duplexed to a concentration of 10 nmol/10 µL in calcium-and magnesium-enriched artificial CSF.^49^ Mice were anesthetized with isoflurane. Animals were monitored throughout the entirety of the procedure. Once the animal was no longer responsive to stimulus (toe pinch), the head was shaved and placed on a stereotaxic frame with a heat pack to maintain temperature throughout the procedure. Using aseptic technique, a small incision was made to the skin just above the skull to expose bregma. Using coordinates from bregma (mediolateral +-1mm, posterior −0.2mm, ventral −2.5mm), the skull was drilled with a burr to make a small opening, and the needle placed for injection. The rate of injection was 750 nl/min. This process was done bilaterally. Once the bilateral side was completed, the needle was removed and the skin is sutured. The mouse was given analgesia of Meloxicam ER and saline. The animal was then placed in a recovery cage with a heating pad under part of the cage to help continue thermoregulation. The animal was monitored until fully sternal. Post-procedure monitoring was for 72 hours and then once a week until the end of the study.

### mRNA measurement

*In vitro* and *in vivo* mRNA was quantified using the branched DNA Quantigene Singleplex assay (Invitrogen ThermoFisher QS0016). For *in vitro* samples, lysate was made using the Quantigene lysis mix (Invitrogen ThermoFisher QS0016). For *in vivo* samples, one 1.5 mm x 1.5 mm x 1 mm punch was lysed into 600 µL homogenizing mix (Invitrogen ThermoFisher QG0517). Branched DNA Quantigene assay performed based on the assay-provided protocol. ThermoFisher Quantigene probes used include *EXO1* (Human, SA-13511), *Exo1* (Mouse, SB-3032575), *FAN1* (Human, SA-3006370), *Fan1* (Mouse, SA-3051642), *HPRT* (Human, SA-10030), *Hprt* (Mouse, SB-15463), *HTT* (Human, SA-50339), *Htt* (Mouse, SB-14150), *MLH1* (Human, SA-3007450), *Mlh1* (Mouse, SA-3030156), *MLH3* (Human, SA-3007450), *Mlh3* (Mouse, SB-3046475), *MSH2* (Human, SA-3002245), *Msh2* (Mouse, SA-3030207), *MSH3* (Human, SA-3002246), *Msh3* (Mouse, SA-3030208), *MSH6* (Human, SA-3001488), *Msh6* (Mouse, SA-3030210), *PMS1* (Human, SA-3002741), *Pms1* (SA-3046971), *PMS2* (Human, SA-3002748), and *Pms2* (Mouse, SA-3030786).

### Protein analysis by Western blot

Frozen tissue punches were homogenized on ice in 10 mM HEPES pH7.2, 250 mM sucrose, 1 mM EDTA + protease inhibitor tablet (Roche) + phosphatase inhibitors (1mM NaF and 1mM Na3VO4), then sonicated for 10 seconds. Protein concentration was determined using the Bradford method (BioRad kit) then equal amounts of protein (10ug) were separated by SDS-PAGE using 3-8% Tris-acetate Criterion gels (BioRad) and transferred to nitrocellulose using a TransblotTurbo apparatus (BioRad). Blots were incubated in blocking buffer (5% blotting-grade buffer (BioRad) in TBST) for 1 hour at room temperature, primary antibody diluted in blocking buffer overnight at 4°C, then peroxidase-labeled secondary antibody diluted in blocking buffer for 1 hour at room temperature (1:2500 for anti-rabbit IgG, 1:5000 for anti-mouse and anti-sheep IgG, Jackson ImmunoResearch). See Table S2 for primary antibodies and dilutions. Blots were imaged with SuperSignal West Pico Plus Chemiluminescent substrate (ThermoFisher) and a CCD camera (AlphaInnotech) or ChemiDoc XRS+ with Image Lab software (BioRad). Pixel intensity quantification was performed using ImageJ software by manually circling each band and multiplying the area by average signal intensity to determine the total signal intensity per each band. The signal intensity for each protein of interest was normalized to GAPDH or B-actin loading control.

### Fragment analysis PCR

CAG lengths and somatic instability were measured with fragment analysis PCR. DNA was isolated from 1.5mm x 1.5mm x 1 mm flash-frozen striatum or medial cortex punch using the Qiagen DNeasy Blood and Tissue Kit (Qiagen Cat. 69506). Once extracted and purified, 150 ng DNA was loaded per PCR reaction. PCR to measure the CAG length was performed using the AmpliTaq 360 DNA Polymerase kit (ThermoFisher Cat. 4398818). Forward primer CAG1: 6FAM-ATG AAG GCC TTC GAG TCC CTC AAG TCC TTC. Reverse primer HU3: GGC GGC TGA GGA AGC TGA GGA. PCR Primers were ordered from and made by Integrated DNA Technology (Coralville, IA). Thermocycling performed at: 94°C, 90s; [94°C, 30s; 63.4°C, 30s; 72°C, 90s] x 35, 72°C, 10m. Fragment analysis performed at the University of Arizona Genetics Core. Somatic instability was analyzed by PeakScanner software by ThermoFisher. Somatic instability was quantified by the index validated by Lee et al. 2010^53^.

### Mass Spectrometry (MS) Sample Preparation for Spectral Library Generation

Unless otherwise stated, 1.5 mL or 0.5 mL Microresico Low Bind Tubes (Amuza, Inc. via Eicom USA, RC9.201.70 or RC9.201.60) were used. Three different samples were used as the basis for experimental spectral library generation:

#### (I) Striatal tissue punches from control Q111 mice

Mouse striatal punch biopsies were lysed in 50 µL of 5% SDS, 50 mM HEPES (pH 8.0), 150 mM NaCl, 5 mM EDTA, 1x HALT Protease Inhibitor Cocktail (Thermo Fisher, 78438), 25 mM tris(2-carboxyethyl)phosphine (TCEP, ThermoFisher 77720), and 50 mM chloroacetamide (CAM, Neta Scientific, SIAL-22790-250G-F). Samples were sonicated 20 times in a cup horn sonicator and subsequently heated at 95°C for 5 minutes, 3 times. Between cycles, samples were centrifuged at 15,000 rpm for 5 minutes on a benchtop centrifuge to pellet material to ensure maximal protein recovery. One aliquot was subjected to In-Solution Digestion (ISD), while another aliquot was subjected to suspension trapping (S-Trap) digestion, as described below.

For ISD, the samples were subjected to methanol/chloroform precipitation by the sequential addition of 200 µL HPLC-MS methanol (Fisher Scientific A456-4), 50 µL HPLC-MS chloroform (Thermo Fisher Scientific AA43685K2), and 150 µL UHPLC-MS water (Thermo Fisher Scientific W81CS), with vortexing after the addition of each solution. Samples were then centrifuged at 15,000 rpm for 5 minutes at 4°C. The protein disk was then carefully rotated and adhered to the side of the sample tube, and both aqueous and organic layers were carefully removed simultaneously. Protein disks were washed an additional 2 times with ice cold 150 µL HPLC-MS methanol, centrifuged at 15,000 rpm for 2 minutes at 4°C. Protein pellet was then resuspended in 50 µL 50 mM HEPES (pH 8.0), and protein concentration was estimated using the BCA assay (Thermo Fisher 23225), after which 50 µg of protein was aliquoted and adjusted to 0.5 mg/mL and digested with 1 µg of sequencing grade trypsin (Thermo Fisher 90059) for 16 hours at 37°C with shaking at 600 rpm in an Eppendorf Thermomixer.

For S-Trap digestion, the samples were lysed as described above, except TCEP and CAM were excluded from the lysis buffer. Protein concentration was estimated with the BCA assay, and 50 µg of protein was aliquoted and reduced and alkylated with 25 mM TCEP and 50 mM CAM, respectively, at 70°C for 20 minutes. The samples were then adjusted to 2.5% phosphoric acid (Sigma Aldrich 79617-250ML), and a 6x sample volume was added of 100 mM TEAB (Thermo Fisher PI90114) dissolved in 90% MS-grade methanol (pH = 7.5, adjusted with phosphoric acid). The samples were then applied to an S-Trap Micro Column (Protifi LLC, C002-MICRO) placed in an empty 1.5 mL Eppendorf receiver tube and centrifuged at 4,000 xg for 30 seconds. The samples were then washed with 150 µL of Binding Buffer and centrifuged at 4,000 xg for 30 seconds 3 additional times. The samples were then washed 3 times with 150 µL of 50% HPLC-MS methanol/50% HPLC chloroform with centrifugation at 4,000 xg for 30 seconds. The samples were then washed an additional 2 times with 150 µL of Binding Buffer with centrifugation at 4,000 xg for 30 seconds. The S-Traps were then transferred into a new, clean Low Bind tube, and 100 µL of 50 mM TEAB (pH 8.5) with 1 µg sequencing grade trypsin, with digestion performed for 16 hours at 37°C without shaking in an Eppendorf Thermomixer. Peptides were eluted from the S-Trap via the sequential addition of 50 µL of 50 mM TEAB (pH 8.5), 0.2% formic acid (FA, Thermo Fisher 28905) in 99.8% UHPLC-MS water, and then 0.2% FA in 80% UHPLC-MS acetonitrile (Thermo Fisher A9561CS) and 19.8% UHPLC-MS water, with centrifugation at 4,000 x g for 1 minute.

Both ISD and S-Trap samples were concentrated by vacuum centrifugation. Concentrated peptides were then separated into 8 fractions with the Pierce High pH Reversed-Phase Peptide Fractionation Kit (Thermo Fisher PI84868), following the manufacturer’s protocol. Fractions 1 and 8 were combined into a single fraction. The peptide fractions were dried down by vacuum centrifugation, and resuspended in 50 µL 0.015% n-dodecyl-β-D-maltoside (DDM, Neta Scientific Inc CMX-21950-5G), 0.1% FA, and 99.885% UHPLC-MS water (MS Resuspension Solution). Peptide concentrations were estimated via NanoDrop using the Scopes method (Scopes, 1974), and samples were adjusted to achieve a peptide concentration of 150 ng/µL in 20 µL MS Resuspension Solution in MS Autosampler Vials (Fisher Scientific 6PMCK36LVW) for MS acquisition.

#### (II) Cytoplasmic and Nuclear fractions from striatal mouse tissues

Mouse striatal tissues from Q20 and Q140 homozygous knock-in mouse models were previously processed^54^ and used for this study as described below.

#### (III) Purified human recombinant mismatch repair (MMR) proteins

Purified recombinant human MSH3, MSH2, MSH6, MLH1, MLH3, PMS1, PMS2, and FAN1 were provided by the CHDI Foundation and processed as described in Greco et al., in preparation.^117^ Briefly, aliquots of each protein (100 ng) were mixed and digested by the S-Trap protocol as described above. Recovered peptides were dried down by vacuum centrifugation and solubilized in MS Resuspension Solution to achieve 500 pg/µL per protein based on digested input. Peptides (0.5 µL) were analyzed by nanoLC-MS/MS. Peptides were separated by LC over a 45 min reverse phase gradient and analyzed by tandem mass spectrometry using DDA-PASEF, with instrument parameters described below in “*MS Acquisition for Spectral Library Generation”*.

### MS Sample Preparation for Quantitative Analysis of striatal tissues from siRNA-treated Q111 mice

Mouse striatal punch biopsies for quantitative analysis were prepared as described above for the samples prepared for Spectral Library Generation, except samples were prepared using the ISD method and were not subjected to peptide fractionation. After digestion, samples were adjusted to 1% trifluoroacetic acid (TFA, Thermo Fisher PI28904), and peptides were desalted via Stage Tip desalting with styrenedivinylbenzene reverse phase sulfonate (SDB-RPS, CDS Analytical 98-0604-0226-4EA). Five disks were assembled by punching out with a Hamilton 14-gauge needle (Hamilton 90514) and carefully packed into a p200 pipette tip (USA Scientific 1112-1720). Stage Tips were conditioned with 50 µL 100% HPLC-MS methanol and centrifuged at 600 xg for 1 minute. Stage Tips were sequentially conditioned with 50 µL of 0.5% FA in 80% UHPLC-MS acetonitrile and 19.5% UHPLC-MS water (Wash Buffer 2) and then 50 µL of 0.5% FA in 99.5% UHPLC-MS water (Wash Buffer 1), centrifuged at 600 x g for 1 minute after the addition of each Wash Buffer. The full volume (∼51 µL) of each sample was then applied to the conditioned Stage Tips and centrifuged at 600 x g for 5 minutes. Samples were then sequentially washed with 50 µL each of Wash Buffer 1 and then Wash Buffer 2, with centrifugation at 600 x g for 5 minutes after the addition of each Wash Buffer. Peptides were then eluted with 50 µL 5% ammonium hydroxide in 80% UHPLC-MS acetonitrile and 15% UHPLC-MS water. Eluted peptides were dried down in a speed-vac, and then subsequently resuspended in 50 µL MS Resuspension Solution. Peptide concentrations were estimated as described above, and samples were adjusted to achieve a peptide concentration of 300 ng/µL in 20 µL MS Resuspension Solution in MS Autosampler Vials for MS acquisition.

### MS Acquisition for Spectral Library Generation

MS analysis was performed on a Bruker timsTOF Ultra equipped with a nanoElute 2 autosampler. Samples from (I) peptide fractions of Q111 striatal tissues (ISD and S-Trap) were injected (1 µL) and resolved by a PepSep Ultra column (Bruker Scientific 1893484) equipped with a 10 µm emitter (Bruker Scientific 1811112). An equal amount of fractionated samples from ISD digests was injected and resolved by an Ion Opticks Aurora Ultimate column (Ion Opticks, AUR3-25075C18-CSI). Peptide separation was achieved with a 120 minute linear gradient of 0.1% FA in UHPLC-MS water (Buffer A) and 0.1% FA in UHPLC-MS acetonitrile (Buffer B). The flow rate was set to 200 nL/minute, with a linear gradient of 2% Buffer B to 35% Buffer B from 0 to 120 minutes, then Buffer B increased to 95% at 120.5 minutes, and was maintained until 127 min.

For MS acquisition, the standard Bruker DDA-PASEF method was used, with a polyhedron for DDA acquisition that covered the 100-1,700 *m/z* region and 0.45-1.55 1/k_0_ region. The TIMS ramp and accumulation times were matched and set to 45ms for recombinant MMR samples, otherwise 100 ms. TIMS ramp cycles were set at 5 for MMR samples, and 10 for all others. Active exclusion was turned on. Precursors were released from exclusion after 0.4 minutes and were considered for re-acquisition if the precursor intensity was 4 times the previous precursor intensity. High sensitivity detection was turned on to increase the number of peptides identified for inclusion in the spectral library.

Prior to MS acquisition of quantitative samples, unfractionated control striatal ISD and S-Trap samples (prepared as described above) were injected using the same LC gradient and MS methods as for quantitative analysis described below, except DDA-PASEF was performed instead of DIA-PASEF. The DDA-PASEF analyses were processed by FragPipe (v23) and py_diAID^118^ to generate optimized DIA-PASEF windows (see Data S1) for quantitative analysis of striatal tissues from siRNA-treated Q111 mice.

### DIA-PASEF MS Analysis of Striatal Tissues from siRNA-treated Q111 mice

MS analysis was performed on a Bruker timsTOF Ultra equipped with a nanoElute 2 autosampler. Unfractionated ISD samples were injected (0.5 µL of 300 ng/uL) on either an Ion Opticks Aurora Ultimate column or a PepSep Ultra column equipped with a 10 µm emitter. For the Ion Opticks Aurora Ultimate column, peptide separation was achieved with a 45-minute linear gradient of Buffer A and Buffer B. The flow rate was set to 200 nL/minute, with a linear gradient of 4% Buffer B to 35% Buffer B from 0 to 45 minutes, then Buffer B increased to 95% at 45.5 minutes, and was maintained until 52.27 minutes. For the PepSep Ultra column, peptide separation was achieved with a 55-minute linear gradient of Buffer A and Buffer B. The flow rate was set to 300 nL/minute and 4% at the beginning of the run, but was reduced to 200 nL/minute and 4% B by 5 minutes, with a linear gradient of 4% Buffer B to 35% Buffer B from 5 to 55 minutes, then the flow rate increased to 300 nL/minute and 95% B 55.5 minutes, and was maintained until 62.27 minutes. For MS acquisition, DIA-PASEF acquisition was performed using a 16 window scheme optimized by py_diAID, as described above. The MS1, MS2, and TIMS scan regions were 100 to 1,700 *m/z,* 250 to 1600 *m/z*, and 0.6-1.50 1/K_0_. The TIMS ramp and accumulation times were matched at 45 ms, and high sensitivity detection was turned off.

### Assembly of Experimental Spectral Library and Computational Analysis of DIA-PASEF Datasets

A hybrid DDA and DIA spectral library was generated and used for quantification of DIA-PASEF data using the FragPipe Informatics pipeline (v24.0). Briefly, the DIA_SpecLib_Quant_diaPASEF workflow was used for processing with default parameters, except where noted. Three DDA-PASEF datasets were loaded: (I) 3 sets of 7 bRP fractions from Q111 striatum, (II) 16 bRP fractions from nuclear and cytoplasmic fractions of KI HD mice (MassIVE MSV000094509), and (III) 3 technical replicates of tryptic peptides from human MMR proteins. DDA-PASEF datasets were processed with the DDA+ data type to allow for identification of up to 5 precursors per MS/MS scan (Yu *et al*, 2025). DIA-PASEF data were processed using diaTracer,^119^ which creates pseudo-MS/MS spectra from timsTOF DIA-PASEF data that can be searched by MSFragger ^120^ and scored by Percolator (analogous to DDA-PASEF MSFragger search results). The hybrid search results were filtered to 1% FDR at the protein, peptide, and spectra level, then assembled into a experimental library, which was used by DIA-NN (ver. 2.3.2) to perform quantification of DIA-PASEF experimental files (n = 103 files). The default DIA-NN settings in FragPipe were used except quantification mode was set to “QuantUMS (Precision)” and the following commands were specified “--window 9 --mass-acc 15 --mass-acc-ms1 15”. The resulting “pg.matrix.tsv” protein group quantitative report was used for downstream analysis in Excel and R.

The pg.matrix file was saved in xlsx format and worksheets were added for metadata annotation and proteins of interest lists (see Data S1). The resulting excel file was processed by an in-house R-based script written for this study, which integrated preprocessing, quality control, differential analysis, visualization, and pathway enrichment (Data S2). Briefly, the excel file containing the quantitative matrix and metadata were imported together along with user-specified gene lists representing siRNA targets or DNA repair and handling (DNA-RH) proteins.^117^ Protein rows were filtered using sequence-counts (N > 1, except N > 0 for DNA-RH subset) and valid values in 50% of samples. The siRNA target FAN1 did not meet this empirical threshold, but after manual inspection of its assigned MS/MS spectrum, it was retained. When gene lists were provided, processed data were subseted by targets and downstream analysis proceeded. For siRNA targets, knockdown efficiency was evaluated after after imputation of missing values using a left-censoring strategy based on low-abundance truncated Gaussian sampling. For DNA-RH and full dataset analyses imputation was not performed. A batch group covariate was specified in the metadata. To evaluate the effect of batch correction, PCA of pre and post-batch corrected quantitative matrices using RemoveBatchEffect function in limma (v3.66.0) (see Figure S12B). Replicate-level QC diagnostics were additionally computed for groups, including missing-value fraction, median intensity shifts, correlation to the group median profile, and PCA-based outlier metrics before and after batch correction. Based on this analysis, one of the twelve NTC controls was removed from downstream analysis. For statistical testing, log2-transformed protein abundances were normalized relative to the control condition and analyzed using linear modeling with limma followed by DEqMS peptide count variance modeling.^56^ Statistical modeling was performed by DEqMS,^56^ using the uncorrected matrix, with covariates were specified in the design model. Results were summarized as per-protein log2 fold-changes, p-values, adjusted p-values, derived percent-of-control values, as well as annotation columns for significance (Data S3).

Downstream visualization included per-contrast volcano plots, differential summary plots, target-protein heatmaps (for DNA-RH analysis), and heatmaps of significantly regulated proteins (for proteome analysis). Heatmaps were generated by ComplexHeatmap (v2.26.1) and ggplot2 (v4.0.2) using calculated log2 values from limma analysis, specifically siRNA group / NTC and (NTC / aCSF). For DNA-RH analysis, the target protein heatmap was clustered by protein (rows). For proteome analysis, the differential target heatmapwas clustered by siRNA group (column) and protein (row)-clustered. Cluster-wise summaries were exported together with spaghetti-profile plots (Data S4). Finally, functional enrichment analyses were conducted by clusterProfiler (v4.18.4) on differential protein clusters using a redundancy-reduced (clusterProfiler::simplify) mouse GO Biological Process annotation (org.Hm.eg.db, v3.22.0).

### Statistics

Statistics were performed using GraphPad Prism Version 10.2.3. When comparing two groups, a Student’s t-test was used. When comparing more than two groups with one variable, a one-way ANOVA with multiple comparisons was used. When comparing two variables, a two-way ANOVA with multiple comparisons was used. *In vitro* experiments were performed in N=3 biological replicates. *In vivo* experiments were performed with N=6-10 mice per condition. Where graphically reported, statistics are denoted as * p < 0.05, ** p < 0.01, *** p < 0.001, **** p < 0.0001.

## Acknowledgements

The authors thank the CHDI Foundation for supporting this work (CHDI Foundation A-17281, A-5038, A-18331). IMC acknowledges contributions from NIH NIAID AI174515, NIH NIGMS GM11414, and Allen Family Philanthropies towards mass spectrometry instrumentation. JNB was supported by NINDS F31 NS132424.

## Data availability

All data generated are provided within the main or supplemental materials. The proteome datasets produced from this study are available at MassIVE (ID: MSV000102182) ProteomeXchange (ID: PXD079836).

## Author contributions

AK, JB, DO, TFV, MF, BP, MD, IC, NA conceptualized the study. Study design was by AK, NA, IC, MD, BP, MF, TFV, JB, DO, TG. Chemistry and siRNA synthesis was performed by DO, NY, HF, DE, NM, BB, DC, SA. In vitro work was performed by JB, DO, EL, EF, CF, RF, NY, SA, NG, AM, KG, DC. In vivo animal work was completed by AS, JB, EL, RF, SA, DO. Tissue processing and analysis was performed by JB, ES, EL, RF, SH. Proteomics work was designed and completed by TG, JH, BP, IC. Methodology was designed by AK, IC, AS, TG, JH, DE, DO, ES, HF. Resources were provided by TFV, MF, BP, MD, IC, NA, AK, DO, TG. Writing (original draft) was performed by JB, TG, AK, NA, MD, with review and editing by JB, TG, ES, TFV, MF, BP, MD, IC, NA, AK. Data visualization was by JB, TG, IC, TFV, MF, BP, MD, NA, AK. Supervision for this work was provided by AK, NA, IC, MD, BP, MF, TFV.

## References

1. Tabrizi, S.J., Flower, M.D., Ross, C.A., and Wild, E.J. (2020). Huntington disease: new insights into molecular pathogenesis and therapeutic opportunities. Nat Rev Neurol 16, 529–546. 10.1038/s41582-020-0389-4.

2. MacDonald, M.E., Ambrose, C.M., Duyao, M.P., Myers, R.H., Lin, C., Srinidhi, L., Barnes, G., Taylor, S.A., James, M., Groot, N., et al. (1993). A novel gene containing a trinucleotide repeat that is expanded and unstable on Huntington’s disease chromosomes. Cell 72, 971–983. 10.1016/0092-8674(93)90585-E.

3. Aronin, N., Chase, K., Young, C., Sapp, E., Schwarz, C., Matta, N., Kornreich, R., Landwehrmeyer, B., Bird, E., Beal, M.F., and, et al. (1995). CAG expansion affects the expression of mutant Huntingtin in the Huntington’s disease brain. Neuron 15, 1193–1201. 10.1016/0896-6273(95)90106-x.

4. Gusella, J.F., Wexler, N.S., Conneally, P.M., Naylor, S.L., Anderson, M.A., Tanzi, R.E., Watkins, P.C., Ottina, K., Wallace, M.R., Sakaguchi, A.Y., and, et al. (1983). A polymorphic DNA marker genetically linked to Huntington’s disease. Nature 306, 234–238. 10.1038/306234a0.

5. Aldous, S.G., Smith, E.J., Landles, C., Osborne, G.F., Cañibano-Pico, M., Nita, I.M., Phillips, J., Zhang, Y., Jin, B., Hirst, M.B., et al. (2024). A CAG repeat threshold for therapeutics targeting somatic instability in Huntington’s disease. Brain 147, 1784–1798. 10.1093/brain/awae063.

6. Belgrad, J., Summers, A., Landles, C., Greene, J.R., Hildebrand, S., Knox, E., Sapp, E., Yamada, N., Furgal, R., Miller, R., et al. (2025). Blocking somatic repeat expansion and lowering huntingtin via RNA interference synergize to prevent Huntington’s disease pathogenesis in mice. bioRxiv, 2025.2006.2024.661398. 10.1101/2025.06.24.661398.

7. Handsaker, R.E., Kashin, S., Reed, N.M., Tan, S., Lee, W.-S., McDonald, T.M., Morris, K., Kamitaki, N., Mullally, C.D., Morakabati, N., et al. (2025). Long somatic DNA-repeat expansion drives neurodegeneration in Huntington disease. Cell. 10.1016/j.cell.2024.11.038.

8. Wang, N., Zhang, S., Langfelder, P., Ramanathan, L., Plascencia, M., Gao, F., Vaca, R., Gu, X., Deng, L., Dionisio, L.E., et al. (2025). Distinct mismatch-repair complex genes set neuronal CAG-repeat expansion rate to drive selective pathogenesis in HD mice. Cell. 10.1016/j.cell.2025.01.031.

9. Lee, J.-M., Correia, K., Loupe, J., Kim, K.-H., Barker, D., Hong, E.P., Chao, M.J., Long, J.D., Lucente, D., Vonsattel, J.P.G., et al. (2019). CAG Repeat Not Polyglutamine Length Determines Timing of Huntington’s Disease Onset. Cell 178, 887–900.e814. 10.1016/j.cell.2019.06.036.

10. Lee, J.M., Huang, Y., Orth, M., Gillis, T., Siciliano, J., Hong, E., Mysore, J.S., Lucente, D., Wheeler, V.C., Seong, I.S., et al. (2022). Genetic modifiers of Huntington disease differentially influence motor and cognitive domains. Am J Hum Genet 109, 885–899. 10.1016/j.ajhg.2022.03.004.

11. Goold, R., Flower, M., Moss, D.H., Medway, C., Wood-Kaczmar, A., Andre, R., Farshim, P., Bates, G.P., Holmans, P., Jones, L., and Tabrizi, S.J. (2019). FAN1 modifies Huntington’s disease progression by stabilizing the expanded HTT CAG repeat. Hum Mol Genet 28, 650–661. 10.1093/hmg/ddy375.

12. McAllister, B., Donaldson, J., Binda, C.S., Powell, S., Chughtai, U., Edwards, G., Stone, J., Lobanov, S., Elliston, L., Schuhmacher, L.N., et al. (2022). Exome sequencing of individuals with Huntington’s disease implicates FAN1 nuclease activity in slowing CAG expansion and disease onset. Nat Neurosci 25, 446–457. 10.1038/s41593-022-01033-5.

13. Deshmukh, A.L., Porro, A., Mohiuddin, M., Lanni, S., Panigrahi, G.B., Caron, M.C., Masson, J.Y., Sartori, A.A., and Pearson, C.E. (2021). FAN1, a DNA Repair Nuclease, as a Modifier of Repeat Expansion Disorders. J Huntingtons Dis 10, 95–122. 10.3233/jhd-200448.

14. Dragileva, E., Hendricks, A., Teed, A., Gillis, T., Lopez, E.T., Friedberg, E.C., Kucherlapati, R., Edelmann, W., Lunetta, K.L., MacDonald, M.E., and Wheeler, V.C. (2009). Intergenerational and striatal CAG repeat instability in Huntington’s disease knock-in mice involve different DNA repair genes. Neurobiol Dis 33, 37–47. 10.1016/j.nbd.2008.09.014.

15. Ferguson, R., Goold, R., Coupland, L., Flower, M., and Tabrizi, S.J. (2024). Therapeutic validation of MMR-associated genetic modifiers in a human ex vivo model of Huntington disease. Am J Hum Genet 111, 1165–1183. 10.1016/j.ajhg.2024.04.015.

16. Gomes-Pereira, M., Fortune, M.T., Ingram, L., McAbney, J.P., and Monckton, D.G. (2004). Pms2 is a genetic enhancer of trinucleotide CAG·CTG repeat somatic mosaicism: implications for the mechanism of triplet repeat expansion. Human Molecular Genetics 13, 1815–1825. 10.1093/hmg/ddh186.

17. Kadyrova, L.Y., Gujar, V., Burdett, V., Modrich, P.L., and Kadyrov, F.A. (2020). Human MutLγ, the MLH1–MLH3 heterodimer, is an endonuclease that promotes DNA expansion. Proceedings of the National Academy of Sciences 117, 3535–3542. 10.1073/pnas.1914718117.

18. Kovalenko, M., Dragileva, E., St Claire, J., Gillis, T., Guide, J.R., New, J., Dong, H., Kucherlapati, R., Kucherlapati, M.H., Ehrlich, M.E., et al. (2012). Msh2 acts in medium-spiny striatal neurons as an enhancer of CAG instability and mutant huntingtin phenotypes in Huntington’s disease knock-in mice. PLoS One 7, e44273. 10.1371/journal.pone.0044273.

19. McLean, Z.L., Gao, D., Correia, K., Roy, J.C.L., Shibata, S., Farnum, I.N., Valdepenas-Mellor, Z., Kovalenko, M., Rapuru, M., Morini, E., et al. (2024). Splice modulators target PMS1 to reduce somatic expansion of the Huntington’s disease-associated CAG repeat. Nature Communications 15, 3182. 10.1038/s41467-024-47485-0.

20. Mouro Pinto, R., Arning, L., Giordano, J.V., Razghandi, P., Andrew, M.A., Gillis, T., Correia, K., Mysore, J.S., Grote Urtubey, D.M., Parwez, C.R., et al. (2020). Patterns of CAG repeat instability in the central nervous system and periphery in Huntington’s disease and in spinocerebellar ataxia type 1. Hum Mol Genet 29, 2551–2567. 10.1093/hmg/ddaa139.

21. Mouro Pinto, R., Murtha, R., Azevedo, A., Douglas, C., Kovalenko, M., Ulloa, J., Crescenti, S., Burch, Z., Oliver, E., Vitalo, A., et al. (2025). In vivo CRISPR–Cas9 genome editing in mice identifies genetic modifiers of somatic CAG repeat instability in Huntington’s disease. Nature Genetics. 10.1038/s41588-024-02054-5.

22. O’Reilly, D., Belgrad, J., Ferguson, C., Summers, A., Sapp, E., McHugh, C., Mathews, E., Boudi, A., Buchwald, J., Ly, S., et al. (2023). Di-valent siRNA-mediated silencing of MSH3 blocks somatic repeat expansion in mouse models of Huntington’s disease. Mol Ther 31, 1661–1674. 10.1016/j.ymthe.2023.05.006.

23. Roy, J.C.L., Vitalo, A., Andrew, M.A., Mota-Silva, E., Kovalenko, M., Burch, Z., Nhu, A.M., Cohen, P.E., Grabczyk, E., Wheeler, V.C., and Mouro Pinto, R. (2021). Somatic CAG expansion in Huntington’s disease is dependent on the MLH3 endonuclease domain, which can be excluded via splice redirection. Nucleic Acids Res 49, 3907–3918. 10.1093/nar/gkab152.

24. Slean, M.M., Panigrahi, G.B., Castel, A.L., Pearson, A.B., Tomkinson, A.E., and Pearson, C.E. (2016). Absence of MutSβ leads to the formation of slipped-DNA for CTG/CAG contractions at primate replication forks. DNA Repair (Amst) 42, 107–118. 10.1016/j.dnarep.2016.04.002.

25. Wheeler, V.C., and Dion, V. (2021). Modifiers of CAG/CTG Repeat Instability: Insights from Mammalian Models. J Huntingtons Dis 10, 123–148. 10.3233/jhd-200426.

26. Wheeler, V.C., Lebel, L.A., Vrbanac, V., Teed, A., te Riele, H., and MacDonald, M.E. (2003). Mismatch repair gene Msh2 modifies the timing of early disease in Hdh(Q111) striatum. Hum Mol Genet 12, 273–281. 10.1093/hmg/ddg056.

27. Guo, J., Gu, L., Leffak, M., and Li, G.M. (2016). MutSβ promotes trinucleotide repeat expansion by recruiting DNA polymerase β to nascent (CAG)n or (CTG)n hairpins for error-prone DNA synthesis. Cell Res 26, 775–786. 10.1038/cr.2016.66.

28. Lin, Y., Dent, S.Y., Wilson, J.H., Wells, R.D., and Napierala, M. (2010). R loops stimulate genetic instability of CTG.CAG repeats. Proc Natl Acad Sci U S A 107, 692–697. 10.1073/pnas.0909740107.

29. Iyer, R.R., and Pluciennik, A. (2021). DNA Mismatch Repair and its Role in Huntington’s Disease. J Huntingtons Dis 10, 75–94. 10.3233/jhd-200438.

30. Putnam, C.D. (2016). Evolution of the methyl directed mismatch repair system in Escherichia coli. DNA Repair (Amst) 38, 32–41. 10.1016/j.dnarep.2015.11.016.

31. Piscitelli, J.M., and Manhart, C.M. (2025). The GHKL ATPase Family as a Paradigm for MutL Homolog Function in DNA Mismatch Repair. Int J Mol Sci 26. 10.3390/ijms262412157.

32. Nakamori, M., Pearson, C.E., and Thornton, C.A. (2011). Bidirectional transcription stimulates expansion and contraction of expanded (CTG)*(CAG) repeats. Hum Mol Genet 20, 580–588. 10.1093/hmg/ddq501.

33. Bunting, E.L., Donaldson, J., Cumming, S.A., Olive, J., Broom, E., Miclăuș, M., Hamilton, J., Tegtmeyer, M., Zhao, H.T., Brenton, J., et al. (2025). Antisense oligonucleotide-mediated MSH3 suppression reduces somatic CAG repeat expansion in Huntington’s disease iPSC-derived striatal neurons. Sci Transl Med 17, eadn4600. 10.1126/scitranslmed.adn4600.

34. Alterman, J.F., Godinho, B.M.D.C., Hassler, M.R., Ferguson, C.M., Echeverria, D., Sapp, E., Haraszti, R.A., Coles, A.H., Conroy, F., Miller, R., et al. (2019). A divalent siRNA chemical scaffold for potent and sustained modulation of gene expression throughout the central nervous system. Nature Biotechnology 37, 884–894. 10.1038/s41587-019-0205-0.

35. Godinho, B.M.D.C., Knox, E.G., Hildebrand, S., Gilbert, J.W., Echeverria, D., Kennedy, Z., Haraszti, R.A., Ferguson, C.M., Coles, A.H., Biscans, A., et al. (2022). PK-modifying anchors significantly alter clearance kinetics, tissue distribution, and efficacy of therapeutics siRNAs. Molecular Therapy Nucleic Acids 29, 116–132. 10.1016/j.omtn.2022.06.005.

36. Goold, R., Hamilton, J., Menneteau, T., Flower, M., Bunting, E.L., Aldous, S.G., Porro, A., Vicente, J.R., Allen, N.D., Wilkinson, H., et al. (2021). FAN1 controls mismatch repair complex assembly via MLH1 retention to stabilize CAG repeat expansion in Huntington’s disease. Cell Reports 36, 109649. 10.1016/j.celrep.2021.109649.

37. Iyer, R.R., Pluciennik, A., Burdett, V., and Modrich, P.L. (2006). DNA Mismatch Repair: Functions and Mechanisms. Chemical Reviews 106, 302–323. 10.1021/cr0404794.

38. Iyer, R.R., Pluciennik, A., Genschel, J., Tsai, M.-S., Beese, L.S., and Modrich, P. (2010). MutL&#x3b1; and Proliferating Cell Nuclear Antigen Share Binding Sites on MutS&#x3b2; *. Journal of Biological Chemistry 285, 11730–11739. 10.1074/jbc.M110.104125.

39. Lee, J.-M., Correia, K., Loupe, J., Kim, K.-H., Barker, D., Hong, E.P., Chao, M.J., Long, J.D., Lucente, D., Vonsattel, J.P.G., et al. (2019). CAG Repeat Not Polyglutamine Length Determines Timing of Huntington&#x2019;s Disease Onset. Cell 178, 887–900.e814. 10.1016/j.cell.2019.06.036.

40. Loupe, J.M., Pinto, R.M., Kim, K.H., Gillis, T., Mysore, J.S., Andrew, M.A., Kovalenko, M., Murtha, R., Seong, I., Gusella, J.F., et al. (2020). Promotion of somatic CAG repeat expansion by Fan1 knock-out in Huntington’s disease knock-in mice is blocked by Mlh1 knock-out. Hum Mol Genet 29, 3044–3053. 10.1093/hmg/ddaa196.

41. Mätlik, K., Baffuto, M., Kus, L., Deshmukh, A.L., Davis, D.A., Paul, M.R., Carroll, T.S., Caron, M.-C., Masson, J.-Y., Pearson, C.E., and Heintz, N. (2024). Cell-type-specific CAG repeat expansions and toxicity of mutant Huntingtin in human striatum and cerebellum. Nature Genetics 56, 383–394. 10.1038/s41588-024-01653-6.

42. Tomé, S., Manley, K., Simard, J.P., Clark, G.W., Slean, M.M., Swami, M., Shelbourne, P.F., Tillier, E.R., Monckton, D.G., Messer, A., and Pearson, C.E. (2013). MSH3 polymorphisms and protein levels affect CAG repeat instability in Huntington’s disease mice. PLoS Genet 9, e1003280. 10.1371/journal.pgen.1003280.

43. Monopoli, K.R., Korkin, D., and Khvorova, A. (2023). Asymmetric trichotomous partitioning overcomes dataset limitations in building machine learning models for predicting siRNA efficacy. Molecular Therapy Nucleic Acids 33, 93–109. 10.1016/j.omtn.2023.06.010.

44. Khvorova, A., and Watts, J.K. (2017). The chemical evolution of oligonucleotide therapies of clinical utility. Nat Biotechnol 35, 238–248. 10.1038/nbt.3765.

45. Egli, M., and Manoharan, M. (2023). Chemistry, structure and function of approved oligonucleotide therapeutics. Nucleic Acids Res 51, 2529–2573. 10.1093/nar/gkad067.

46. Hassler, M.R., Turanov, A.A., Alterman, J.F., Haraszti, R.A., Coles, A.H., Osborn, M.F., Echeverria, D., Nikan, M., Salomon, W.E., Roux, L., et al. (2018). Comparison of partially and fully chemically-modified siRNA in conjugate-mediated delivery in vivo. Nucleic Acids Research 46, 2185–2196. 10.1093/nar/gky037.

47. Wheeler, V.C., Auerbach, W., White, J.K., Srinidhi, J., Auerbach, A., Ryan, A., Duyao, M.P., Vrbanac, V., Weaver, M., Gusella, J.F., et al. (1999). Length-dependent gametic CAG repeat instability in the Huntington’s disease knock-in mouse. Hum Mol Genet 8, 115–122. 10.1093/hmg/8.1.115.

48. Elkayam, E., Parmar, R., Brown, C.R., Willoughby, J.L., Theile, C.S., Manoharan, M., and Joshua-Tor, L. (2017). siRNA carrying an (E)-vinylphosphonate moiety at the 5΄ end of the guide strand augments gene silencing by enhanced binding to human Argonaute-2. Nucleic Acids Res 45, 3528–3536. 10.1093/nar/gkw1171.

49. Miller, R., Paquette, J., Barker, A., Sapp, E., McHugh, N., Bramato, B., Yamada, N., Alterman, J., Echeveria, D., Yamada, K., et al. (2024). Preventing acute neurotoxicity of CNS therapeutic oligonucleotides with the addition of Ca(2+) and Mg(2+) in the formulation. Mol Ther Nucleic Acids 35, 102359. 10.1016/j.omtn.2024.102359.

50. Allen, S., O’Reilly, D., Miller, R., Sapp, E., Summers, A., Paquette, J., Echeverria Moreno, D., Bramato, B., McHugh, N., Yamada, K., et al. (2024). mRNA Nuclear Clustering Leads to a Difference in Mutant Huntingtin mRNA and Protein Silencing by siRNAs In Vivo. Nucleic Acid Ther 34, 164–172. 10.1089/nat.2024.0027.

51. Goold, R., Hamilton, J., Menneteau, T., Flower, M., Bunting, E.L., Aldous, S.G., Porro, A., Vicente, J.R., Allen, N.D., Wilkinson, H., et al. (2021). FAN1 controls mismatch repair complex assembly via MLH1 retention to stabilize CAG repeat expansion in Huntington’s disease. Cell Rep 36, 109649. 10.1016/j.celrep.2021.109649.

52. Bhatia, M., Phadte, A.S., Lakhina, A., Monte Carlo III, A.R., Barndt, S., and Pluciennik, A. (2025). DNA extrusion size determines pathway choice during CAG repeat expansion. Nucleic Acids Research 53, gkaf1393. 10.1093/nar/gkaf1393.

53. Lee, J.M., Zhang, J., Su, A.I., Walker, J.R., Wiltshire, T., Kang, K., Dragileva, E., Gillis, T., Lopez, E.T., Boily, M.J., et al. (2010). A novel approach to investigate tissue-specific trinucleotide repeat instability. BMC Syst Biol 4, 29. 10.1186/1752-0509-4-29.

54. Justice, J.L., Greco, T.M., Hutton, J.E., Reed, T.J., Mair, M.L., Botas, J., and Cristea, I.M. (2025). Multi-epitope immunocapture of huntingtin reveals striatum-selective molecular signatures. Mol Syst Biol 21, 492–522. 10.1038/s44320-025-00096-3.

55. Wiśniewski, J.R., Hein, M.Y., Cox, J., and Mann, M. (2014). A “proteomic ruler” for protein copy number and concentration estimation without spike-in standards. Mol Cell Proteomics 13, 3497–3506. 10.1074/mcp.M113.037309.

56. Zhu, Y., Orre, L.M., Zhou Tran, Y., Mermelekas, G., Johansson, H.J., Malyutina, A., Anders, S., and Lehtiö, J. (2020). DEqMS: A Method for Accurate Variance Estimation in Differential Protein Expression Analysis. Mol Cell Proteomics 19, 1047–1057. 10.1074/mcp.TIR119.001646.

57. Edelbrock, M.A., Kaliyaperumal, S., and Williams, K.J. (2013). Structural, molecular and cellular functions of MSH2 and MSH6 during DNA mismatch repair, damage signaling and other noncanonical activities. Mutat Res 743-744, 53–66. 10.1016/j.mrfmmm.2012.12.008.

58. Guerrette, S., Wilson, T., Gradia, S., and Fishel, R. (1998). Interactions of human hMSH2 with hMSH3 and hMSH2 with hMSH6: examination of mutations found in hereditary nonpolyposis colorectal cancer. Mol Cell Biol 18, 6616–6623. 10.1128/mcb.18.11.6616.

59. Kennedy, M.A., Greco, T.M., Song, B., and Cristea, I.M. (2022). HTT-OMNI: A Web-based Platform for Huntingtin Interaction Exploration and Multi-omics Data Integration. Mol Cell Proteomics 21, 100275. 10.1016/j.mcpro.2022.100275.

60. Landles, C., Osborne, G.F., Phillips, J., Canibano-Pico, M., Nita, I.M., Ali, N., Bobkov, K., Greene, J.R., Sathasivam, K., and Bates, G.P. (2024). Mutant huntingtin protein decreases with CAG repeat expansion: implications for therapeutics and bioassays. Brain Commun 6, fcae410. 10.1093/braincomms/fcae410.

61. Abildgaard, A.B., Stein, A., Nielsen, S.V., Schultz-Knudsen, K., Papaleo, E., Shrikhande, A., Hoffmann, E.R., Bernstein, I., Gerdes, A.-M., Takahashi, M., et al. (2019). Computational and cellular studies reveal structural destabilization and degradation of MLH1 variants in Lynch syndrome. eLife 8, e49138. 10.7554/eLife.49138.

62. Gu, X., Richman, J., Langfelder, P., Wang, N., Zhang, S., Bañez-Coronel, M., Wang, H.B., Yang, L., Ramanathan, L., Deng, L., et al. (2022). Uninterrupted CAG repeat drives striatum-selective transcriptionopathy and nuclear pathogenesis in human Huntingtin BAC mice. Neuron 110, 1173–1192.e1177. 10.1016/j.neuron.2022.01.006.

63. Ciosi, M., Maxwell, A., Cumming, S.A., Hensman Moss, D.J., Alshammari, A.M., Flower, M.D., Durr, A., Leavitt, B.R., Roos, R.A.C., Holmans, P., et al. (2019). A genetic association study of glutamine-encoding DNA sequence structures, somatic CAG expansion, and DNA repair gene variants, with Huntington disease clinical outcomes. EBioMedicine 48, 568–580. 10.1016/j.ebiom.2019.09.020.

64. Jordheim, L.P., Sève, P., Trédan, O., and Dumontet, C. (2011). The ribonucleotide reductase large subunit (RRM1) as a predictive factor in patients with cancer. The Lancet Oncology 12, 693–702. 10.1016/S1470-2045(10)70244-8.

65. Chen, Z., Morris, H.R., Polke, J., Wood, N.W., Gandhi, S., Ryten, M., Houlden, H., and Tucci, A. (2025). Repeat expansion disorders. Practical Neurology 25, 204. 10.1136/pn-2023-003938.

66. Paulson, H. (2018). Repeat expansion diseases. Handb Clin Neurol 147, 105–123. 10.1016/b978-0-444-63233-3.00009-9.

67. Malik, I., Kelley, C.P., Wang, E.T., and Todd, P.K. (2021). Molecular mechanisms underlying nucleotide repeat expansion disorders. Nature Reviews Molecular Cell Biology 22, 589–607. 10.1038/s41580-021-00382-6.

68. Depienne, C., and Mandel, J.-L. (2021). 30 years of repeat expansion disorders: What have we learned and what are the remaining challenges? The American Journal of Human Genetics 108, 764–785. 10.1016/j.ajhg.2021.03.011.

69. Lan, Z.-q., Ge, Z.-y., Lv, S.-k., Zhao, B., and Li, C.-x. (2023). The regulatory role of lipophagy in central nervous system diseases. Cell Death Discovery 9, 229. 10.1038/s41420-023-01504-z.

70. Cannavo, E., Gerrits, B., Marra, G., Schlapbach, R., and Jiricny, J. (2007). Characterization of the Interactome of the Human MutL Homologues MLH1, PMS1, and PMS2 *. Journal of Biological Chemistry 282, 2976–2986. 10.1074/jbc.M609989200.

71. Bossy-Wetzel, E., Petrilli, A., and Knott, A.B. (2008). Mutant huntingtin and mitochondrial dysfunction. Trends Neurosci 31, 609–616. 10.1016/j.tins.2008.09.004.

72. Joshi, D.C., Chavan, M.B., Gurow, K., Gupta, M., Dhaliwal, J.S., and Ming, L.C. (2025). The role of mitochondrial dysfunction in Huntington’s disease: Implications for therapeutic targeting. Biomedicine & Pharmacotherapy 183, 117827. 10.1016/j.biopha.2025.117827.

73. Yablonska, S., Ganesan, V., Ferrando, L.M., Kim, J., Pyzel, A., Baranova, O.V., Khattar, N.K., Larkin, T.M., Baranov, S.V., Chen, N., et al. (2019). Mutant huntingtin disrupts mitochondrial proteostasis by interacting with TIM23. Proceedings of the National Academy of Sciences 116, 16593–16602. 10.1073/pnas.1904101116.

74. Park, J.-C., Kim, Y.-J., Han, J.H., Kim, D., Park, M.J., Kim, J., Jang, H.-K., Bae, S., and Cha, H.-J. (2023). MutSalpha and MutSbeta as size-dependent cellular determinants for prime editing in human embryonic stem cells. Molecular Therapy Nucleic Acids 32, 914–922. 10.1016/j.omtn.2023.05.015.

75. Miller, C.J., and Usdin, K. (2022). Mismatch repair is a double-edged sword in the battle against microsatellite instability. Expert Rev Mol Med 24, e32. 10.1017/erm.2022.16.

76. Groothuizen, F.S., Winkler, I., Cristóvão, M., Fish, A., Winterwerp, H.H., Reumer, A., Marx, A.D., Hermans, N., Nicholls, R.A., Murshudov, G.N., et al. (2015). MutS/MutL crystal structure reveals that the MutS sliding clamp loads MutL onto DNA. Elife 4, e06744. 10.7554/eLife.06744.

77. Pluciennik, A., Burdett, V., Baitinger, C., Iyer, R.R., Shi, K., and Modrich, P. (2013). Extrahelical (CAG)/(CTG) triplet repeat elements support proliferating cell nuclear antigen loading and MutLα endonuclease activation. Proceedings of the National Academy of Sciences 110, 12277–12282. doi:10.1073/pnas.1311325110.

78. Yang, X.-W., Han, X.-P., Han, C., London, J., Fishel, R., and Liu, J. (2022). MutS functions as a clamp loader by positioning MutL on the DNA during mismatch repair. Nature Communications 13, 5808. 10.1038/s41467-022-33479-3.

79. Campbell, C.S., Hombauer, H., Srivatsan, A., Bowen, N., Gries, K., Desai, A., Putnam, C.D., and Kolodner, R.D. (2014). Mlh2 is an accessory factor for DNA mismatch repair in Saccharomyces cerevisiae. PLoS Genet 10, e1004327. 10.1371/journal.pgen.1004327.

80. Ranjha, L., Anand, R., and Cejka, P. (2014). The Saccharomyces cerevisiae Mlh1-Mlh3 heterodimer is an endonuclease that preferentially binds to Holliday junctions. J Biol Chem 289, 5674–5686. 10.1074/jbc.M113.533810.

81. Claeys Bouuaert, C., and Keeney, S. (2017). Distinct DNA-binding surfaces in the ATPase and linker domains of MutLγ determine its substrate specificities and exert separable functions in meiotic recombination and mismatch repair. PLoS Genet 13, e1006722. 10.1371/journal.pgen.1006722.

82. Piscitelli, J.M., Witte, S.J., Sakinejad, Y.S., and Manhart, C.M. (2024). The Mlh1-Pms1 endonuclease uses ATP to preserve DNA discontinuities as strand discrimination signals to facilitate mismatch repair. bioRxiv. 10.1101/2024.06.13.598860.

83. Witte, Scott J., Rosa, Isabella M., Collingwood, Bryce W., Piscitelli, Jonathan M., and Manhart, Carol M. (2023). The mismatch repair endonuclease MutLα tethers duplex regions of DNA together and relieves DNA torsional tension. Nucleic Acids Research 51, 2725–2739. 10.1093/nar/gkad096.

84. Manhart, C.M., Ni, X., White, M.A., Ortega, J., Surtees, J.A., and Alani, E. (2017). The mismatch repair and meiotic recombination endonuclease Mlh1-Mlh3 is activated by polymer formation and can cleave DNA substrates in trans. PLOS Biology 15, e2001164. 10.1371/journal.pbio.2001164.

85. Rogacheva, M.V., Manhart, C.M., Chen, C., Guarne, A., Surtees, J., and Alani, E. (2014). Mlh1-Mlh3, a meiotic crossover and DNA mismatch repair factor, is a Msh2-Msh3-stimulated endonuclease. J Biol Chem 289, 5664–5673. 10.1074/jbc.M113.534644.

86. Matos, J., and West, S.C. (2014). Holliday junction resolution: regulation in space and time. DNA Repair (Amst) 19, 176–181. 10.1016/j.dnarep.2014.03.013.

87. Rahman, M.M., Mohiuddin, M., Shamima Keka, I., Yamada, K., Tsuda, M., Sasanuma, H., Andreani, J., Guerois, R., Borde, V., Charbonnier, J.B., and Takeda, S. (2020). Genetic evidence for the involvement of mismatch repair proteins, PMS2 and MLH3, in a late step of homologous recombination. J Biol Chem 295, 17460–17475. 10.1074/jbc.RA120.013521.

88. Dai, J., Sanchez, A., Adam, C., Ranjha, L., Reginato, G., Chervy, P., Tellier-Lebegue, C., Andreani, J., Guérois, R., Ropars, V., et al. (2021). Molecular basis of the dual role of the Mlh1-Mlh3 endonuclease in MMR and in meiotic crossover formation. Proceedings of the National Academy of Sciences 118, e2022704118. doi:10.1073/pnas.2022704118.

89. Pannafino, G., and Alani, E. (2021). Coordinated and Independent Roles for MLH Subunits in DNA Repair. Cells 10. 10.3390/cells10040948.

90. Belgrad, J., Tang, Q., Hildebrand, S., Summers, A., Sapp, E., Echeverria, D., O’Reilly, D., Luu, E., Bramato, B., Allen, S., et al. (2024). A programmable dual-targeting siRNA scaffold supports potent two-gene modulation in the central nervous system. Nucleic Acids Research 52, 6099–6113. 10.1093/nar/gkae368.

91. Oliver, E., Kovalenko, M., Louçã, M., Jiang, A., Westerdahl, J., Correia, K., Jones, B., Saif, F., Romano, N., Sidhu, A., et al. (2026). Somatic CRISPR editing of Msh3 mitigates Huntington’s disease pathology in mice. bioRxiv, 2026.2006.2008.730940. 10.64898/2026.06.08.730940.

92. Papadopoulou, A.S., Alterman, J., Landles, C., Smith, E.J., Conroy, F., Phillips, J., Canibano-Pico, M., Nita, I.M., Osborne, G.F., Iqbal, A., et al. (2025). Lowering the HTT1a transcript as an effective therapy for Huntington’s disease. bioRxiv, 2025.2006.2010.658804. 10.1101/2025.06.10.658804.

93. Kraus-Perrotta, C., and Lagalwar, S. (2016). Expansion, mosaicism and interruption: mechanisms of the CAG repeat mutation in spinocerebellar ataxia type 1. Cerebellum & Ataxias 3, 20. 10.1186/s40673-016-0058-y.

94. Koide, R., Kobayashi, S., Shimohata, T., Ikeuchi, T., Maruyama, M., Saito, M., Yamada, M., Takahashi, H., and Tsuji, S. (1999). A Neurological Disease Caused By an Expanded CAG Trinucleotide Repeat in The TATA-Binding Protein Gene: A New Polyglutamine Disease? Human Molecular Genetics 8, 2047–2053. 10.1093/hmg/8.11.2047.

95. Fratta, P., Collins, T., Pemble, S., Nethisinghe, S., Devoy, A., Giunti, P., Sweeney, M.G., Hanna, M.G., and Fisher, E.M. (2014). Sequencing analysis of the spinal bulbar muscular atrophy CAG expansion reveals absence of repeat interruptions. Neurobiol Aging 35, 443.e441–443. 10.1016/j.neurobiolaging.2013.07.015.

96. Langbehn, D.R., Hayden, M.R., and Paulsen, J.S. (2010). CAG-repeat length and the age of onset in Huntington disease (HD): a review and validation study of statistical approaches. Am J Med Genet B Neuropsychiatr Genet 153b, 397–408. 10.1002/ajmg.b.30992.

97. Foiry, L., Dong, L., Savouret, C., Hubert, L., te Riele, H., Junien, C., and Gourdon, G. (2006). Msh3 is a limiting factor in the formation of intergenerational CTG expansions in DM1 transgenic mice. Hum Genet 119, 520–526. 10.1007/s00439-006-0164-7.

98. Morales, F., Vásquez, M., Santamaría, C., Cuenca, P., Corrales, E., and Monckton, D.G. (2016). A polymorphism in the MSH3 mismatch repair gene is associated with the levels of somatic instability of the expanded CTG repeat in the blood DNA of myotonic dystrophy type 1 patients. DNA Repair (Amst) 40, 57–66. 10.1016/j.dnarep.2016.01.001.

99. Sobczak, K., and Krzyzosiak, W.J. (2004). Imperfect CAG Repeats Form Diverse Structures in SCA1 Transcripts*. Journal of Biological Chemistry 279, 41563–41572. 10.1074/jbc.M405130200.

100. Kacher, R., Lejeune, F.X., David, I., Boluda, S., Coarelli, G., Leclere-Turbant, S., Heinzmann, A., Marelli, C., Charles, P., Goizet, C., et al. (2024). CAG repeat mosaicism is gene specific in spinocerebellar ataxias. Am J Hum Genet 111, 913–926. 10.1016/j.ajhg.2024.03.015.

101. Das, A., Ercan, A.B., and Tabori, U. (2024). An update on central nervous system tumors in germline replication-repair deficiency syndromes. Neuro-Oncology Advances 6. 10.1093/noajnl/vdae102.

102. Edelmann, W., Yang, K., Umar, A., Heyer, J., Lau, K., Fan, K., Liedtke, W., Cohen, P.E., Kane, M.F., Lipford, J.R., et al. (1997). Mutation in the mismatch repair gene Msh6 causes cancer susceptibility. Cell 91, 467–477. 10.1016/s0092-8674(00)80433-x.

103. Guerrini-Rousseau, L., Merlevede, J., Denizeau, P., Andreiuolo, F., Varlet, P., Puget, S., Beccaria, K., Blauwblomme, T., Cabaret, O., Hamzaoui, N., et al. (2024). Glioma oncogenesis in the Constitutional mismatch repair deficiency (CMMRD) syndrome. Neuro-Oncology Advances 6. 10.1093/noajnl/vdae120.

104. Guerrini-Rousseau, L., Varlet, P., Colas, C., Andreiuolo, F., Bourdeaut, F., Dahan, K., Devalck, C., Faure-Conter, C., Genuardi, M., Goldberg, Y., et al. (2019). Constitutional mismatch repair deficiency-associated brain tumors: report from the European C4CMMRD consortium. Neurooncol Adv 1, vdz033. 10.1093/noajnl/vdz033.

105. Kim, H., Lim, K.Y., Park, J.W., Kang, J., Won, J.K., Lee, K., Shim, Y., Park, C.-K., Kim, S.-K., Choi, S.-H., et al. (2022). Sporadic and Lynch syndrome-associated mismatch repair-deficient brain tumors. Laboratory Investigation 102, 160–171. 10.1038/s41374-021-00694-3.

106. Lee, K., Tosti, E., and Edelmann, W. (2016). Mouse models of DNA mismatch repair in cancer research. DNA Repair (Amst) 38, 140–146. 10.1016/j.dnarep.2015.11.015.

107. Martín-López, J.V., and Fishel, R. (2013). The mechanism of mismatch repair and the functional analysis of mismatch repair defects in Lynch syndrome. Fam Cancer 12, 159–168. 10.1007/s10689-013-9635-x.

108. Reyes, G.X., Zhao, B., Schmidt, T.T., Gries, K., Kloor, M., and Hombauer, H. (2020). Identification of MLH2/hPMS1 dominant mutations that prevent DNA mismatch repair function. Communications Biology 3, 751. 10.1038/s42003-020-01481-4.

109. Vageli, D.P., Giannopoulos, S., Doukas, S.G., Kalaitzis, C., Giannakopoulos, S., Giatromanolaki, A., Koukoulis, G.K., and Touloupidis, S. (2013). Mismatch repair hMSH2, hMLH1, hMSH6 and hPMS2 mRNA expression profiles in precancerous and cancerous urothelium. Oncol Lett 5, 283–294. 10.3892/ol.2012.979.

110. Ferguson, C.M., Godinho, B., Echeverria, D., Hassler, M., Vangjeli, L., Sousa, J., McHugh, N., Alterman, J., Hariharan, V., Krishnamurthy, P.M., et al. (2024). A combinatorial approach for achieving CNS-selective RNAi. Nucleic Acids Res 52, 5273–5284. 10.1093/nar/gkae100.

111. Belgrad, J., Fakih, H.H., and Khvorova, A. (2024). Nucleic Acid Therapeutics: Successes, Milestones, and Upcoming Innovation. Nucleic Acid Ther 34, 52–72. 10.1089/nat.2023.0068.

112. Brown, K.M., Nair, J.K., Janas, M.M., Anglero-Rodriguez, Y.I., Dang, L.T.H., Peng, H., Theile, C.S., Castellanos-Rizaldos, E., Brown, C., Foster, D., et al. (2022). Expanding RNAi therapeutics to extrahepatic tissues with lipophilic conjugates. Nature Biotechnology 40, 1500–1508. 10.1038/s41587-022-01334-x.

113. Gentile, J.E., Corridon, T.L., Echeverria, D., Serack, F.E., Kennedy, Z.E., Gallant-Behm, C.L., Hassler, M.R., Kinberger, G., Kamath, N.G., Gross, K.Y., et al. (2024). Divalent siRNA for prion disease. bioRxiv, 2024.2012.2005.627039. 10.1101/2024.12.05.627039.

114. Barker, S.J., Thayer, M.B., Kim, C., Tatarakis, D., Simon, M.J., Dial, R., Nilewski, L., Wells, R.C., Zhou, Y., Afetian, M., et al. (2024). Targeting the transferrin receptor to transport antisense oligonucleotides across the mammalian blood-brain barrier. Sci Transl Med 16, eadi2245. 10.1126/scitranslmed.adi2245.

115. Scahill, R.I., Farag, M., Murphy, M.J., Hobbs, N.Z., Leocadi, M., Langley, C., Knights, H., Ciosi, M., Fayer, K., Nakajima, M., et al. (2025). Somatic CAG repeat expansion in blood associates with biomarkers of neurodegeneration in Huntington’s disease decades before clinical motor diagnosis. Nature Medicine. 10.1038/s41591-024-03424-6.

116. Tabrizi, S.J., Schobel, S., Gantman, E.C., Mansbach, A., Borowsky, B., Konstantinova, P., Mestre, T.A., Panagoulias, J., Ross, C.A., Zauderer, M., et al. (2022). A biological classification of Huntington’s disease: the Integrated Staging System. Lancet Neurol 21, 632–644. 10.1016/s1474-4422(22)00120-x.

117. Greco, T.M. (2026). Development and application of targeted mass spectrometry-based assays for the quantification of DNA repair and handling proteins in human and mouse HD models systems. In Preparation.

118. Skowronek, P., Thielert, M., Voytik, E., Tanzer, M.C., Hansen, F.M., Willems, S., Karayel, O., Brunner, A.D., Meier, F., and Mann, M. (2022). Rapid and In-Depth Coverage of the (Phospho-)Proteome With Deep Libraries and Optimal Window Design for dia-PASEF. Mol Cell Proteomics 21, 100279. 10.1016/j.mcpro.2022.100279.

119. Li, K., Teo, G.C., Yang, K.L., Yu, F., and Nesvizhskii, A.I. (2024). diaTracer enables spectrum-centric analysis of diaPASEF proteomics data. bioRxiv. 10.1101/2024.05.25.595875.

120. Li, Mujia J., Meyer, Larissa C., Meier, N., Witte, J., Maldacker, M., Seredynska, A., Schueler, J., Schilling, O., and Föll, Melanie C. (2025). Spatial Proteomics by Parallel Accumulation-Serial Fragmentation Supported MALDI MS/MS Imaging: A First Glance Into Multiplexed and Spatial Peptide Identification. Rapid Communications in Mass Spectrometry 39, e10006. 10.1002/rcm.10006.

