## Supplemental Data for "Mismatch repair dissection by in vivo RNAi reveals dose-dependent modulators of somatic instability and proteome remodeling in Huntington’s disease"

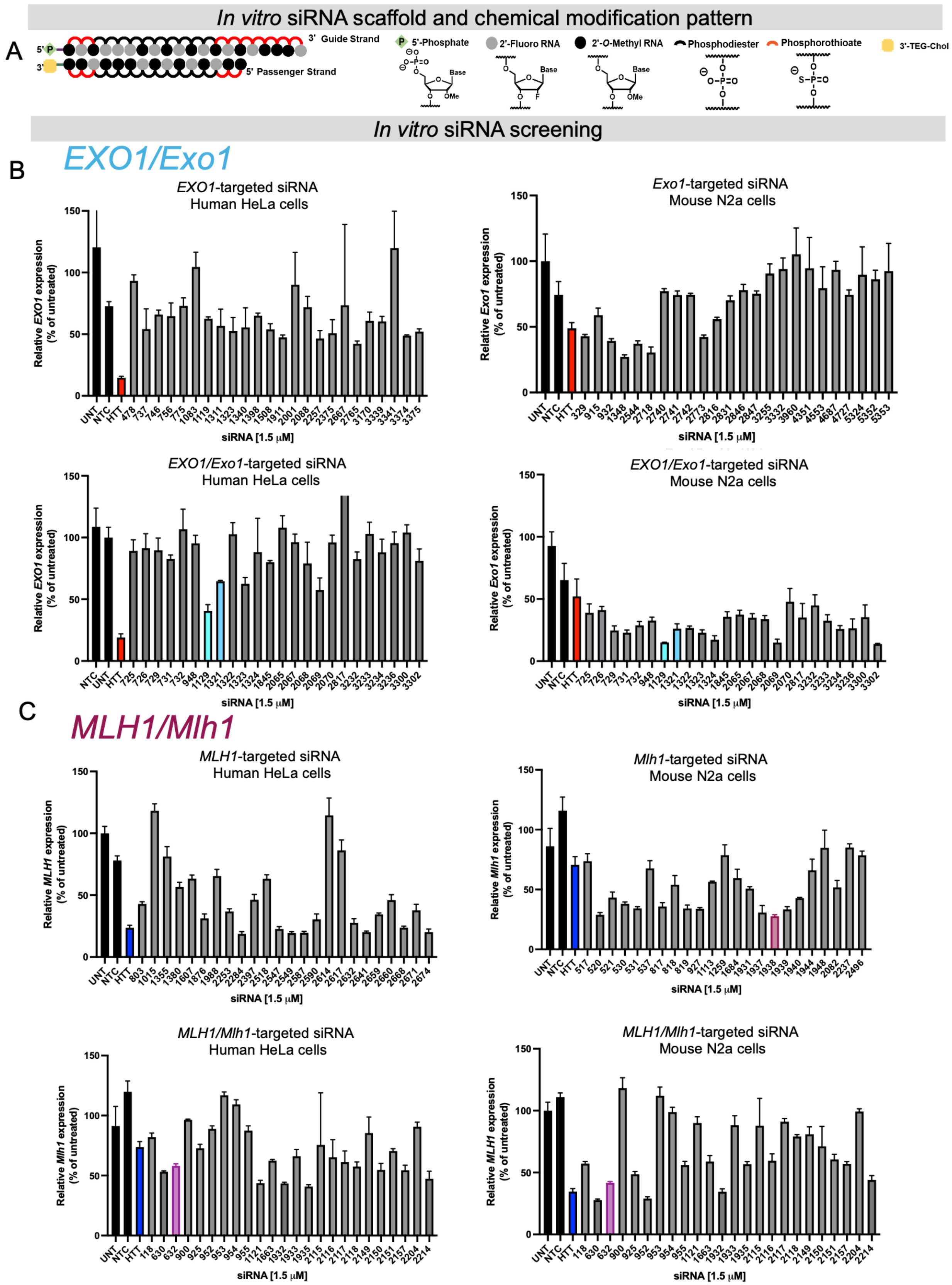
In vitro siRNA scaffold and chemical modification pattern
A
5' P
3'
3' Guide Strand
5' Passenger Strand
5'-Phosphate
2'-Fluoro RNA
2'-O-Methyl RNA
Phosphodiester
Phosphorothioate
3'-TEG-Chol
Base
OMe
F
In vitro siRNA screening
B
EXO1/Exo1
EXO1-targeted siRNA
Human HeLa cells
Relative EXO1 expression (% of untreated)
UNT NTC HTT 478 737 746 756 775 1083 1119 1311 1323 1340 1398 1508 1911 2001 2088 2257 2375 2667 2765 3170 3339 3341 3374 3375
siRNA [1.5 μM]
Exo1-targeted siRNA
Mouse N2a cells
Relative Exo1 expression (% of untreated)
UNT NTC HTT 329 915 932 1348 2544 2718 2740 2741 2742 2773 2816 2831 2846 2847 3255 3332 3960 4351 4553 4687 4727 5324 5352 5353
siRNA [1.5 μM]
EXO1/Exo1-targeted siRNA
Human HeLa cells
Relative EXO1 expression (% of untreated)
NTC UNT HTT 725 726 729 731 732 948 1129 1321 1322 1323 1324 1845 2065 2067 2068 2069 2070 2817 3232 3233 3234 3236 3300 3302
siRNA [1.5 μM]
EXO1/Exo1-targeted siRNA
Mouse N2a cells
Relative Exo1 expression (% of untreated)
UNT NTC HTT 725 726 729 731 732 948 1129 1321 1322 1323 1324 1845 2065 2067 2068 2069 2070 2817 3232 3233 3234 3236 3300 3302
siRNA [1.5 μM]
C
MLH1/Mlh1
MLH1-targeted siRNA
Human HeLa cells
Relative MLH1 expression (% of untreated)
UNT NTC HTT 803 1015 1355 1380 1607 1876 1988 2253 2284 2397 2518 2547 2549 2587 2590 2614 2617 2632 2641 2659 2660 2668 2671 2674
siRNA [1.5 μM]
Mlh1-targeted siRNA
Mouse N2a cells
Relative Mlh1 expression (% of untreated)
UNT NTC HTT 517 520 521 530 531 537 817 818 819 927 1113 1259 1684 1931 1937 1938 1939 1940 1944 1948 2082 2237 2496
siRNA [1.5 μM]
MLH1/Mlh1-targeted siRNA
Human HeLa cells
Relative Mlh1 expression (% of untreated)
UNT NTC HTT 118 630 632 900 925 952 953 954 955 1121 1663 1932 1933 1935 2115 2116 2117 2118 2149 2150 2151 2157 2204 2214
siRNA [1.5 μM]
MLH1/Mlh1-targeted siRNA
Mouse N2a cells
Relative MLH1 expression (% of untreated)
UNT NTC HTT 118 630 632 900 925 952 953 954 955 1121 1663 1932 1933 1935 2115 2116 2117 2118 2149 2150 2151 2157 2204 2214
siRNA [1.5 μM]

### *In vitro* siRNA screening

#### D *MLH3/Mlh3*

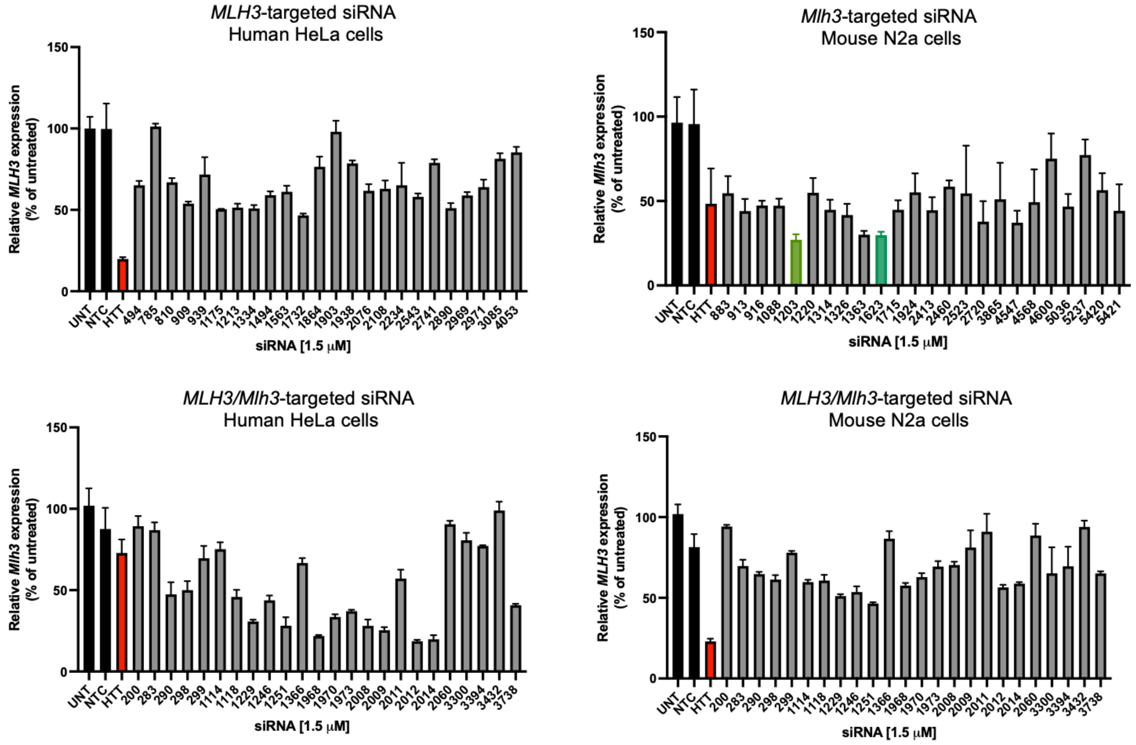

#### E *MSH2/Msh2*

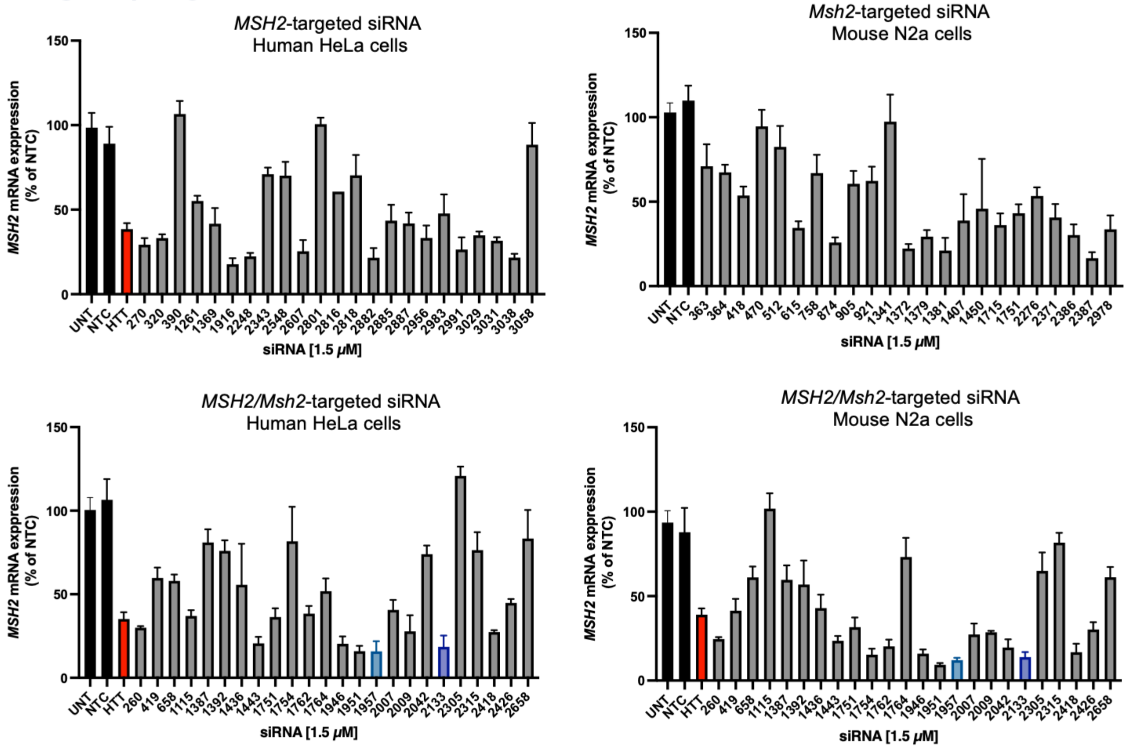

### *In vitro* siRNA screening

F *MSH6/Msh6*

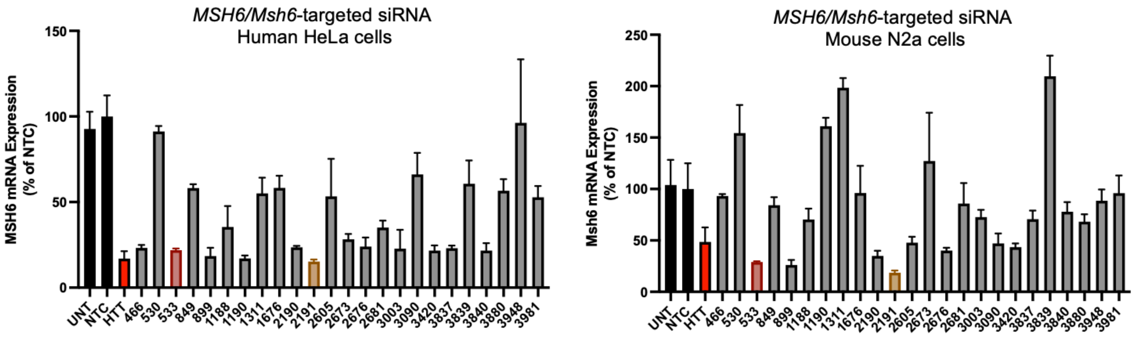

G *FAN1/Fan1*

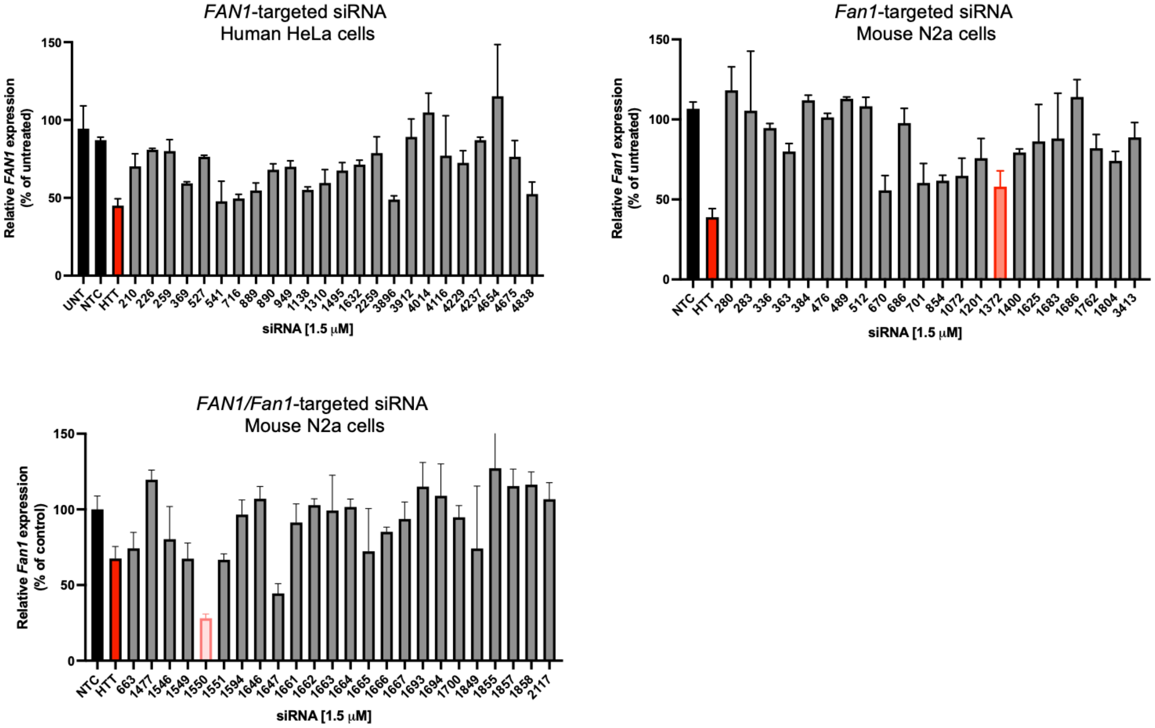

### *In vitro* siRNA screening

H

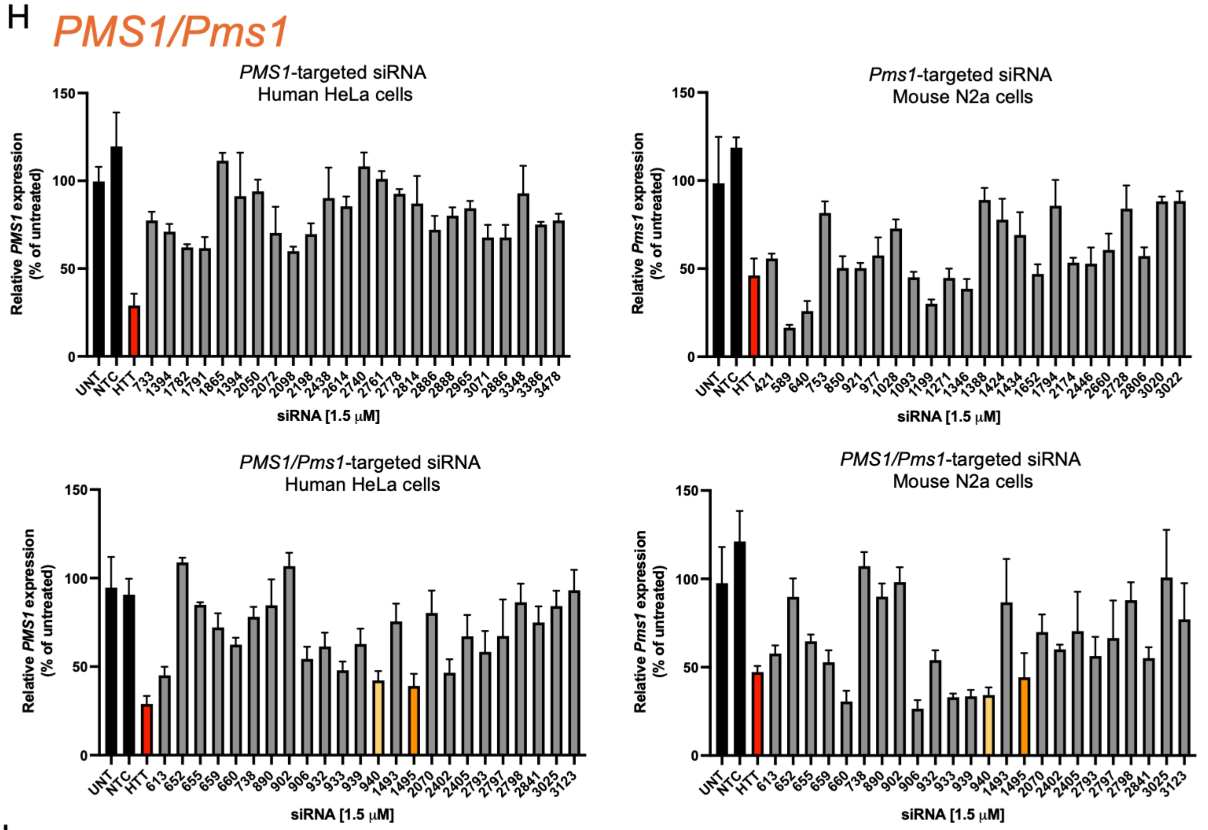

I

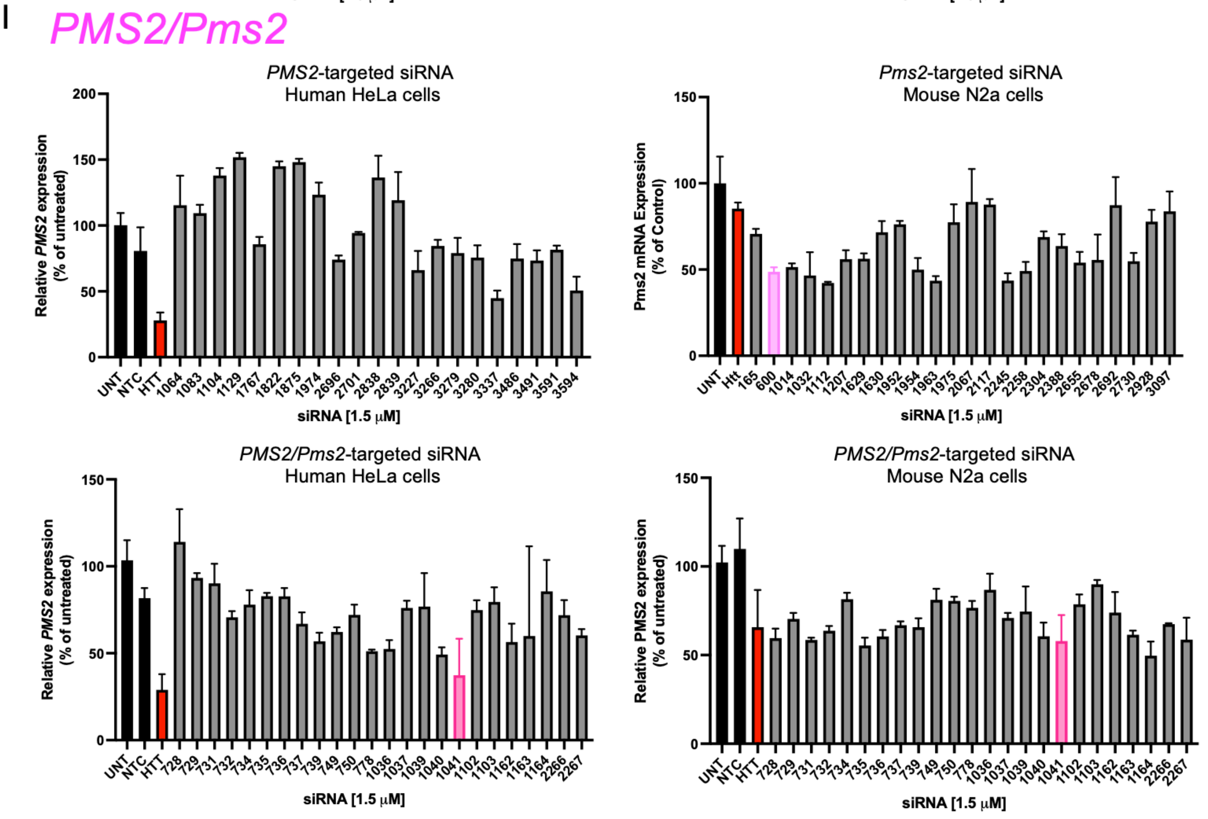

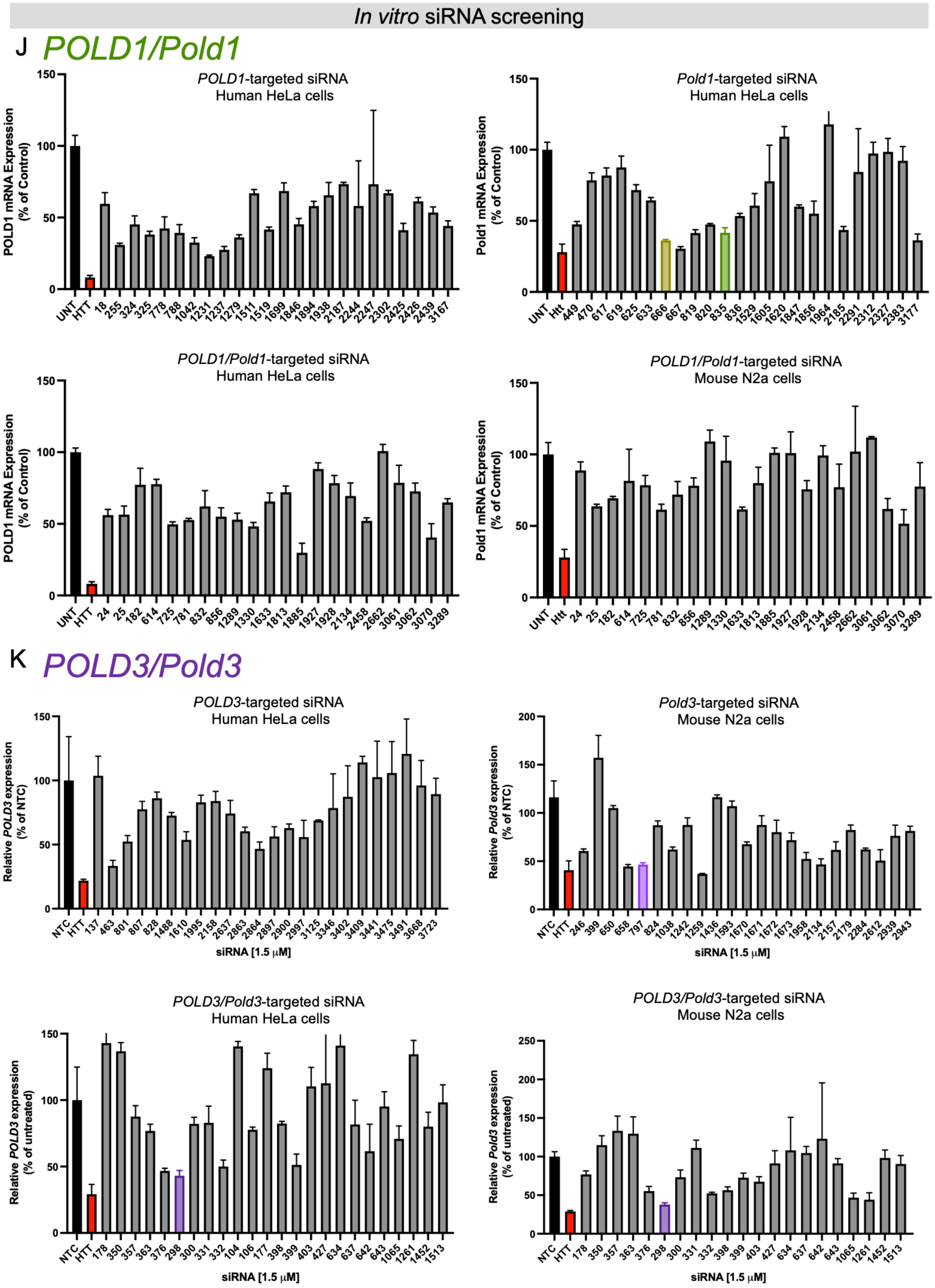

In vitro siRNA screening
J POLD1/Pold1
POLD1-targeted siRNA
Human HeLa cells
POLD1 mRNA Expression (% of Control)
UNT HTT 18 255 324 325 778 788 1042 1231 1237 1279 1511 1519 1699 1846 1894 1938 2187 2244 2247 2302 2425 2426 2439 3167
Pold1-targeted siRNA
Human HeLa cells
Pold1 mRNA Expression (% of Control)
UNT Htt 449 470 617 619 625 633 666 667 819 820 835 836 1529 1605 1620 1847 1856 1964 2185 2291 2312 2327 2383 3177
POLD1/Pold1-targeted siRNA
Human HeLa cells
POLD1 mRNA Expression (% of Control)
UNT HTT 24 25 182 614 725 781 832 856 1289 1330 1633 1813 1885 1927 1928 2134 2458 2662 3061 3062 3070 3289
POLD1/Pold1-targeted siRNA
Mouse N2a cells
Pold1 mRNA Expression (% of Control)
UNT Htt 24 25 182 614 725 781 832 856 1289 1330 1633 1813 1885 1927 1928 2134 2458 2662 3061 3062 3070 3289
K POLD3/Pold3
POLD3-targeted siRNA
Human HeLa cells
Relative POLD3 expression (% of NTC)
NTC HTT 137 463 801 807 828 1488 1610 1995 2158 2637 2863 2864 2897 2900 2997 3125 3346 3402 3409 3441 3475 3491 3668 3723
siRNA [1.5 µM]
Pold3-targeted siRNA
Mouse N2a cells
Relative Pold3 expression (% of NTC)
NTC HTT 246 399 650 658 797 824 1038 1242 1259 1436 1593 1670 1671 1672 1673 1958 2134 2157 2179 2284 2612 2939 2943
siRNA [1.5 µM]
POLD3/Pold3-targeted siRNA
Human HeLa cells
Relative POLD3 expression (% of untreated)
NTC HTT 178 350 357 363 376 298 300 331 332 104 106 177 398 399 403 427 634 637 642 643 1065 1261 1452 1513
siRNA [1.5 µM]
POLD3/Pold3-targeted siRNA
Mouse N2a cells
Relative Pold3 expression (% of untreated)
NTC HTT 178 350 357 363 376 298 300 331 332 398 399 403 427 634 637 642 643 1065 1261 1452 1513
siRNA [1.5 µM]

**Figure S1. Extensive *in vitro* screening identifies potent siRNA targeting mouse and human mismatch repair targets.** (A) chemical scaffold and chemical modification pattern used *in vitro*. siRNA screening for (B) *Exo1/EXO1* (C) *Mlh1/MLH1*, (D) *Mlh3/MLH3*, (E) *Msh2/MSH2*, (F) *Msh6/MSH6* (G) *Fan1/FAN1*, (H) *Pms1/PMS1* (I) *Pms2/PMS2,* (J) *Pold1/POLD1,* (K) *Pold3/POLD3.* Target mRNA quantified using branched DNA assay at 72 hours following passive uptake in HeLa (human) or N2a (mouse) cell lines. 24-48 siRNA were screened per target with unique homology against the Human and Mouse target sequence, or dual homology targeting conserved regions between the two species. UNT- untreated control, NTC, non-targeting control, HTT- huntingtin-targeting siRNA positive control for siRNA and assay function. Each number is a unique code given to the sequences screened. Non-gray bars indicate the siRNA selected for further *in vivo* validation.

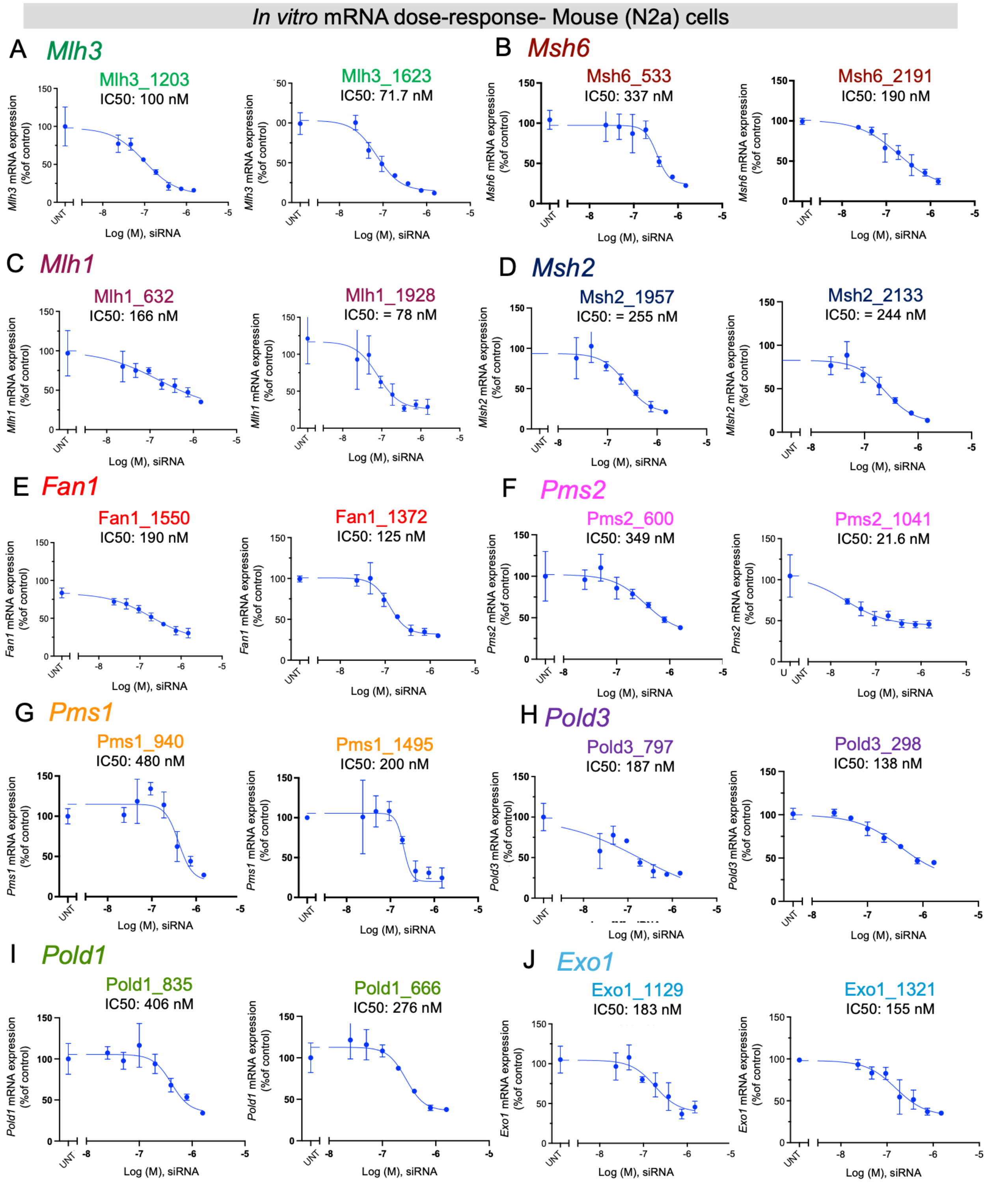

**Figure S2. Top-performing siRNAs silence mouse mismatch repair targets *in vitro*.** 7-point dose response with IC50 for (A) *Mlh3* targeting siRNA, MLH3_1203 and MLH3_1623. (B) *Msh6* targeting siRNA, MSH6_533 and MSH6_2191. (C) *Mlh1* targeting siRNA, MLH1_632 and MLH1_1938. (D) *Msh2*

targeting siRNA, MSH2_1957 and MSH2_2133. (E) *Fan1* targeting siRNA, FAN1_1550 and FAN1_1372. (F) *Pms2* targeting siRNA, PMS2_600 and PMS2_1041. (G) *Pms1* targeting siRNA, PMS1_940 and PMS1_1495. (H) *Pold3* targeting siRNA, POLD3_797 and POLD3_298. (I) *Pold1* targeting siRNA, POLD1_835 and POLD1_666. (J) *Exo1* targeting siRNA, EXO1_1129 and EXO1_1321.

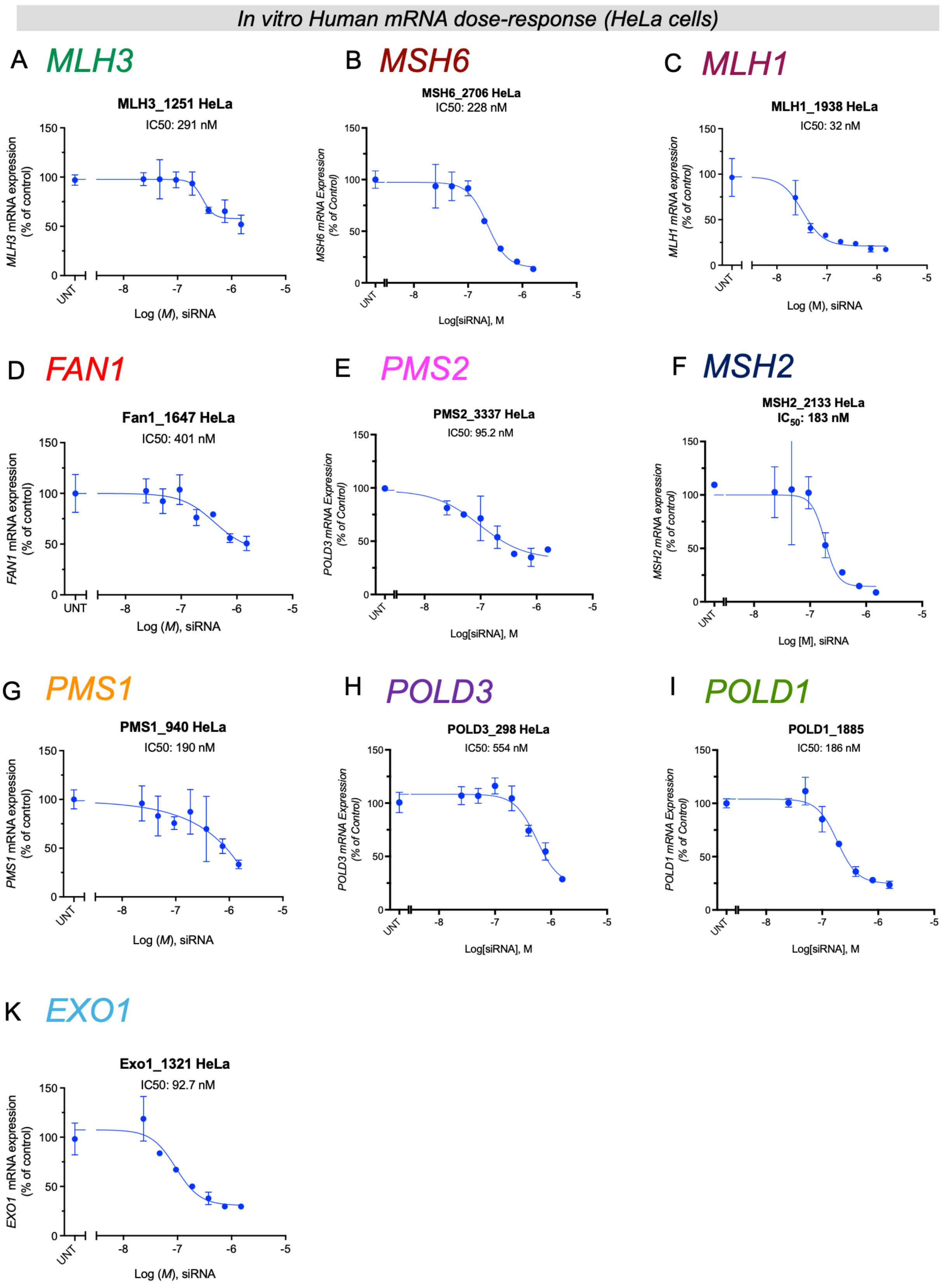

In vitro Human mRNA dose-response (HeLa cells)
A MLH3
MLH3_1251 HeLa
IC50: 291 nM
MLH3 mRNA expression (% of control)
Log (M), siRNA
B MSH6
MSH6_2706 HeLa
IC50: 228 nM
MSH6 mRNA Expression (% of Control)
Log[siRNA], M
C MLH1
MLH1_1938 HeLa
IC50: 32 nM
MLH1 mRNA expression (% of control)
Log (M), siRNA
D FAN1
Fan1_1647 HeLa
IC50: 401 nM
FAN1 mRNA expression (% of control)
Log (M), siRNA
E PMS2
PMS2_3337 HeLa
IC50: 95.2 nM
POLD3 mRNA Expression (% of Control)
Log[siRNA], M
F MSH2
MSH2_2133 HeLa
IC50: 183 nM
MSH2 mRNA expression (% of control)
Log [M], siRNA
G PMS1
PMS1_940 HeLa
IC50: 190 nM
PMS1 mRNA expression (% of control)
Log (M), siRNA
H POLD3
POLD3_298 HeLa
IC50: 554 nM
POLD3 mRNA Expression (% of Control)
Log[siRNA], M
I POLD1
POLD1_1885
IC50: 186 nM
POLD1 mRNA Expression (% of Control)
Log[siRNA], M
K EXO1
Exo1_1321 HeLa
IC50: 92.7 nM
EXO1 mRNA expression (% of control)
Log (M), siRNA
UNT

**Figure S3. Top-performing siRNAs silence human mismatch repair targets *in vitro*.** 7-point dose response with IC50 in HeLa cells for (A) *MLH3*-targeting siRNA, MLH3_1251 (B) *MSH6* targeting siRNA, MSH6_2706. (C) *MLH1* targeting siRNA, MLH1_1938. (D) *FAN1* targeting siRNA, FAN1_1647 (E) *PMS2* targeting siRNA, PMS2_3337, (F) *MSH2* targeting siRNA, MSH2_2133. (G) *PMS1* targeting siRNA, PMS1_940 (I) *POLD3* targeting siRNA, POLD3_298. (J) *POLD1* targeting siRNA, POLD1_1885. (K) *EXO1* targeting siRNA, EXO1_1321.

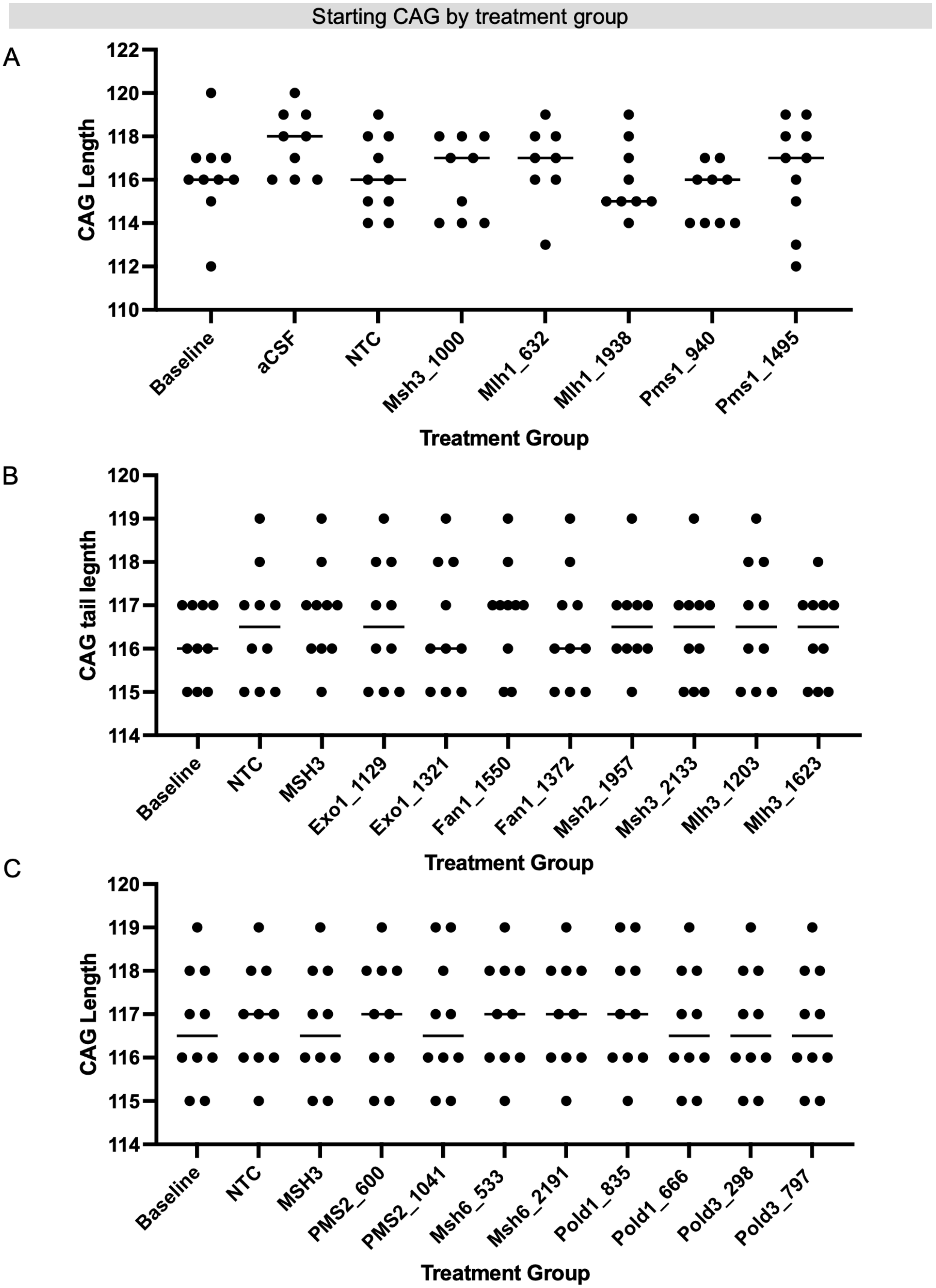
Starting CAG by treatment group
A
CAG Length
122
120
118
116
114
112
110
Baseline
aCSF
NTC
Msh3_1000
Mlh1_632
Mlh1_1938
Pms1_940
Pms1_1495
Treatment Group
B
CAG tail legnth
120
119
118
117
116
115
114
Baseline
NTC
MSH3
Exo1_1129
Exo1_1321
Fan1_1550
Fan1_1372
Msh2_1957
Msh3_2133
Mlh3_1203
Mlh3_1623
Treatment Group
C
CAG Length
120
119
118
117
116
115
114
Baseline
NTC
MSH3
PMS2_600
PMS2_1041
Msh6_533
Msh6_2191
Pold1_835
Pold1_666
Pold3_298
Pold3_797
Treatment Group

**Figure S4. Distribution of inherited CAG lengths in Q111 mice across treatment and experimental cohorts.** Inherited CAG lengths in (A) study 1, (B) study 2, (C) study 3.

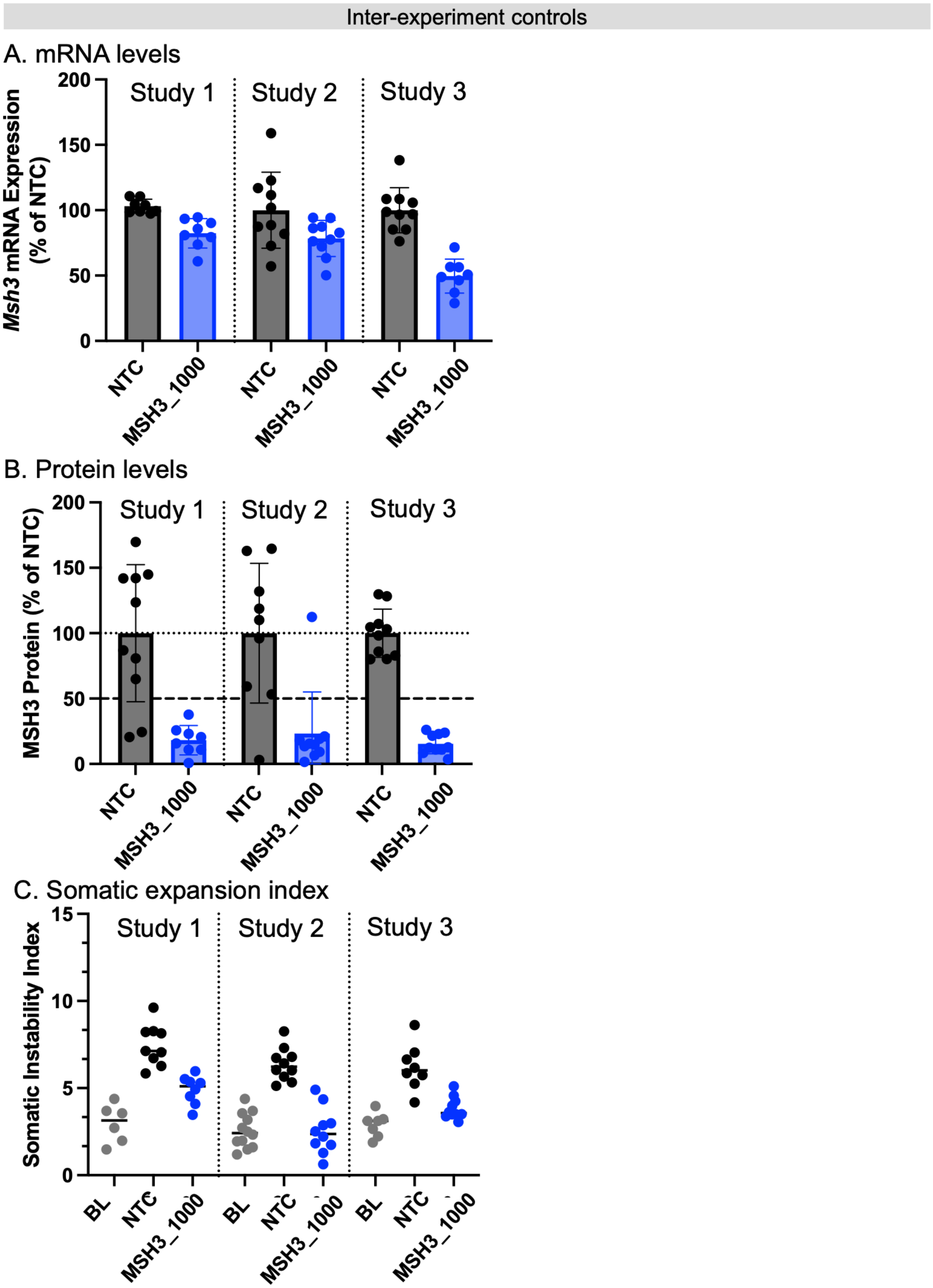

Inter-experiment controls
A. mRNA levels
Msh3 mRNA Expression (% of NTC)
Study 1
Study 2
Study 3
200
150
100
50
0
NTC
MSH3_1000
B. Protein levels
MSH3 Protein (% of NTC)
C. Somatic expansion index
Somatic Instability Index
15
10
5
BL

**Figure S5. MSH3 lowering across experimental studies**. Inter-experiment quality control of *Msh3*/MSH3 (A) mRNA (B) protein or (C) somatic instability index across experimental studies 1, 2, or 3. Baseline ("BL") refers to striatal instability in a 3-month-old Q111 mice at the time of injection. NTC is the non-targeting control group.

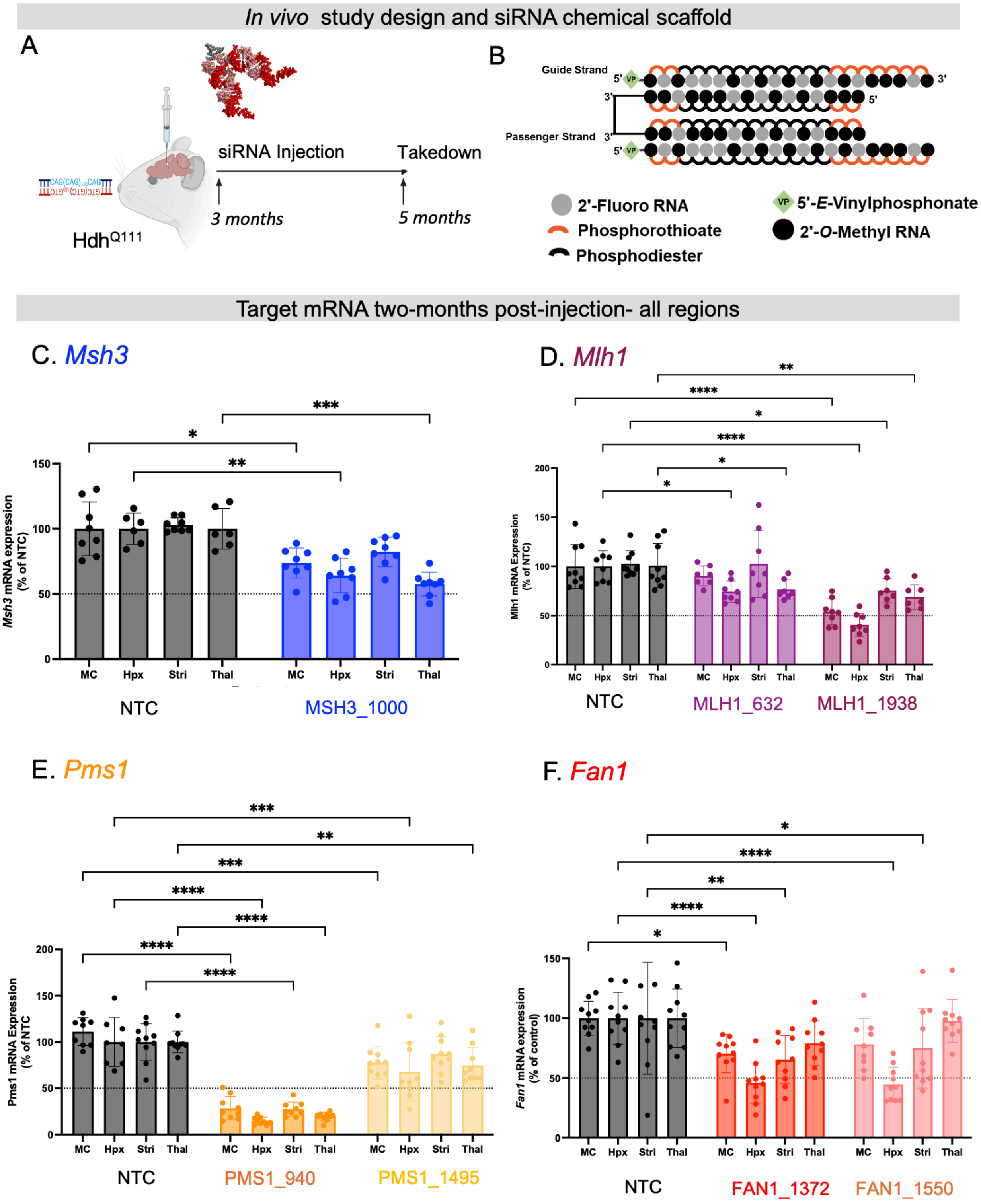

In vivo study design and siRNA chemical scaffold
A
siRNA Injection
Takedown
3 months
5 months
HdhQ111
B
Guide Strand
Passenger Strand
2'-Fluoro RNA
Phosphorothioate
Phosphodiester
5'-E-Vinylphosphonate
2'-O-Methyl RNA
Target mRNA two-months post-injection- all regions
C. Msh3
Msh3 mRNA expression (% of NTC)
MC
Hpx
Stri
Thal
NTC
MSH3_1000
D. Mlh1
Mlh1 mRNA Expression (% of NTC)
NTC
MLH1_632
MLH1_1938
E. Pms1
Pms1 mRNA Expression (% of NTC
NTC
PMS1_940
PMS1_1495
F. Fan1
Fan1 mRNA expression (% of control)
NTC
FAN1_1372
FAN1_1550

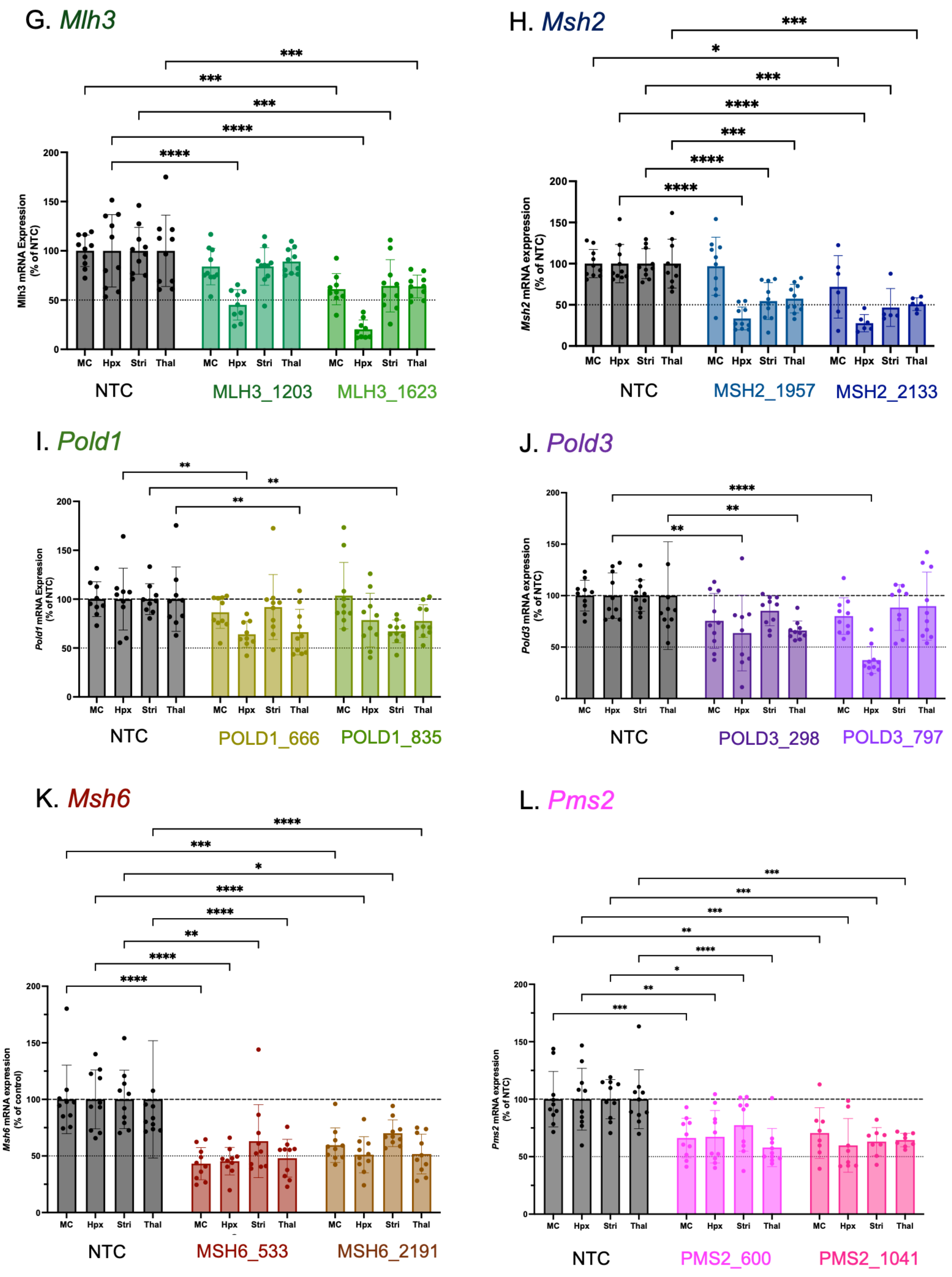

**Figure S6. mRNA levels in Q111 mice two months post-injection with mismatch repair pathway-**

**targeting divalent siRNA.** (A) *in vivo* experimental design (B) divalent siRNA scaffold and chemical modification pattern. mRNA levels of (C) *Msh3* (D) *Mlh1* (E) *Pms1* (F) *Fan1* (G) *Mlh3* (H) *Msh2* (I) *Pold1* (J) *Pold3* (K) *Msh6* or (L) *Pms2* following treatment with respective siRNA. mRNA levels measured by branched DNA assay normalized to non-targeting control (NTC) two months post-injection with 10 nmol/10uL (240 ug) per mouse. Each dot is the mRNA levels from a single mouse. N= 6-10/group. Statistics are two-way ANOVA with multiple comparisons for each brain region. MC- medial cortex, Hpx- hippocampus, Stri- striatum, Thal- thalamus. * $p < 0.05$, ** $p < 0.01$, *** $p < 0.001$, **** $p < 0.0001$.

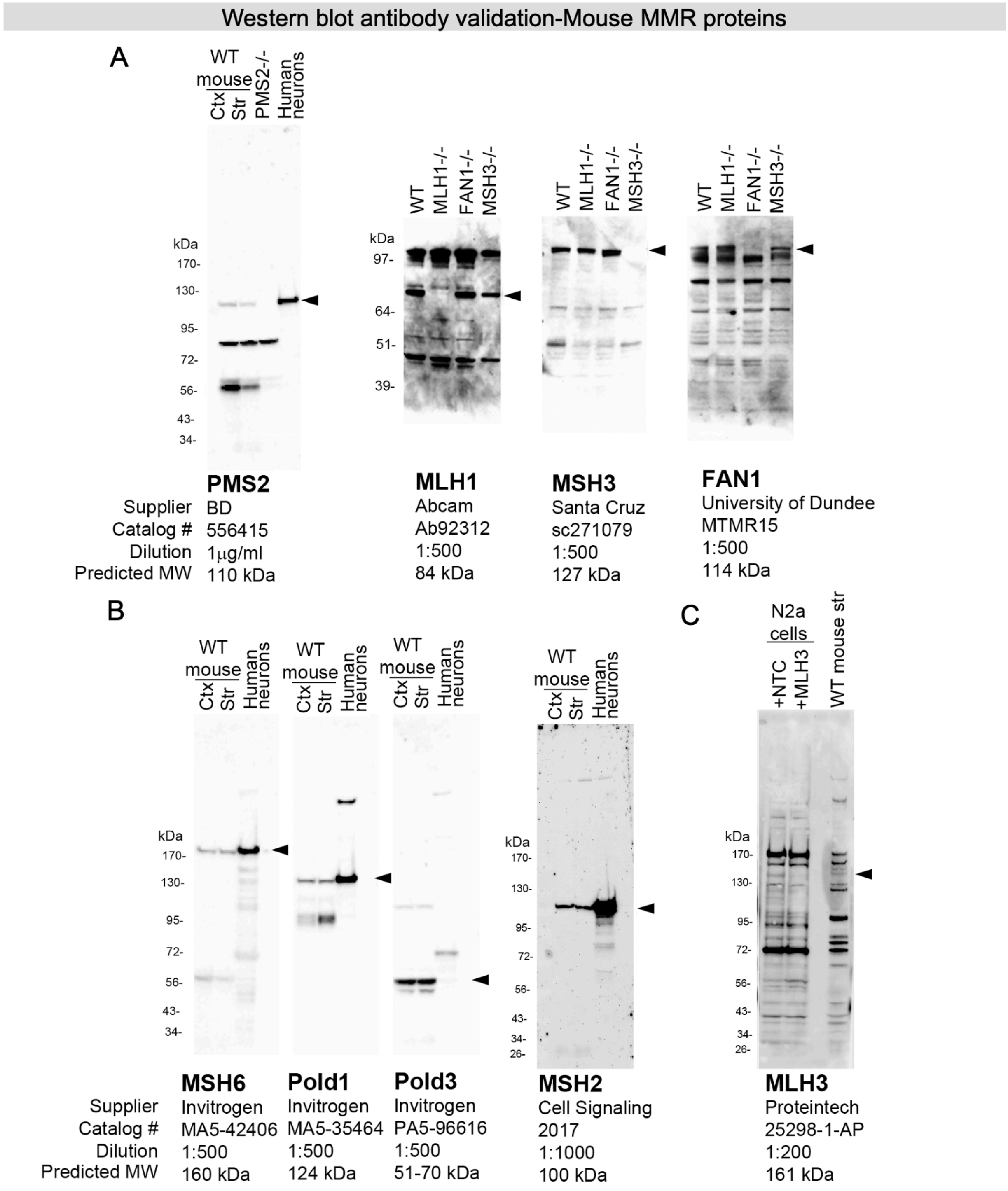

**Figure S7. Mouse MMR antibody validation.** (A) A band was detected at the appropriate size with antibodies to PMS2, MLH1, MSH3, and FAN1 in brain tissue from WT mice but not knockout mice. PMS-/- brain was a generous gift from Dr. William Yang, and MLH1-/-, MSH3-/-, and FAN1-/- brains were a generous gift from Dr. Vanessa Wheeler. (B) MSH6, Pold1, Pold3, and MSH2 antibodies gave a clean signal in WT mouse cortex and striatum, and no knockout tissue was available. (C) The MLH3 antibody detected many bands in the WT mouse striatum, and no knockout tissue was available, so the band of interest was identified using N2a cells treated with MLH3 siRNA. The band at the arrowhead was detected in mouse brain and is absent in the N2a cells treated with MLH3 siRNA. Human control neurons were also run because many of these antibodies were targeted to the human sequence. 10-20μg protein per lane were run on 3-8% Tris-acetate (for PMS2, MSH6, Pold1, Pold3, MSH2, MLH3) or 4-12% Bis-tris (for MLH1, MSH3 and FAN1) gels. Arrowhead to the right of each blot indicates the band of interest.

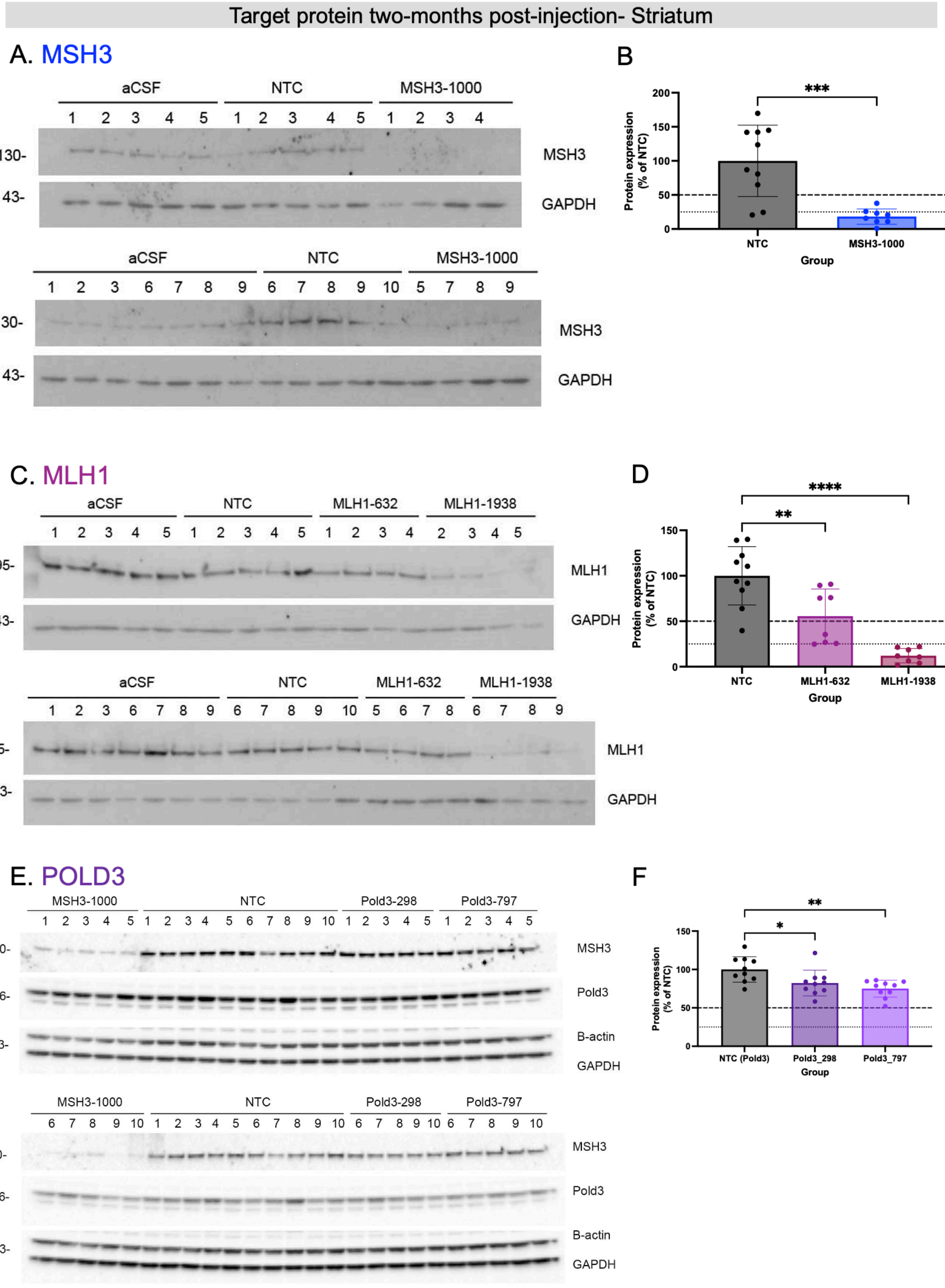
Target protein two-months post-injection- Striatum
A. MSH3
aCSF
NTC
MSH3-1000
130-
43-
MSH3
GAPDH
B
Protein expression (% of NTC)
***
Group
NTC
MSH3-1000
C. MLH1
aCSF
NTC
MLH1-632
MLH1-1938
95-
43-
MLH1
GAPDH
D
****
**
NTC
MLH1-632
MLH1-1938
Group
E. POLD3
MSH3-1000
NTC
Pold3-298
Pold3-797
130-
56-
43-
30-
MSH3
Pold3
B-actin
GAPDH
F
**
*
NTC (Pold3)
Pold3_298
Pold3_797
Group

Target protein two-months post-injection- Striatum

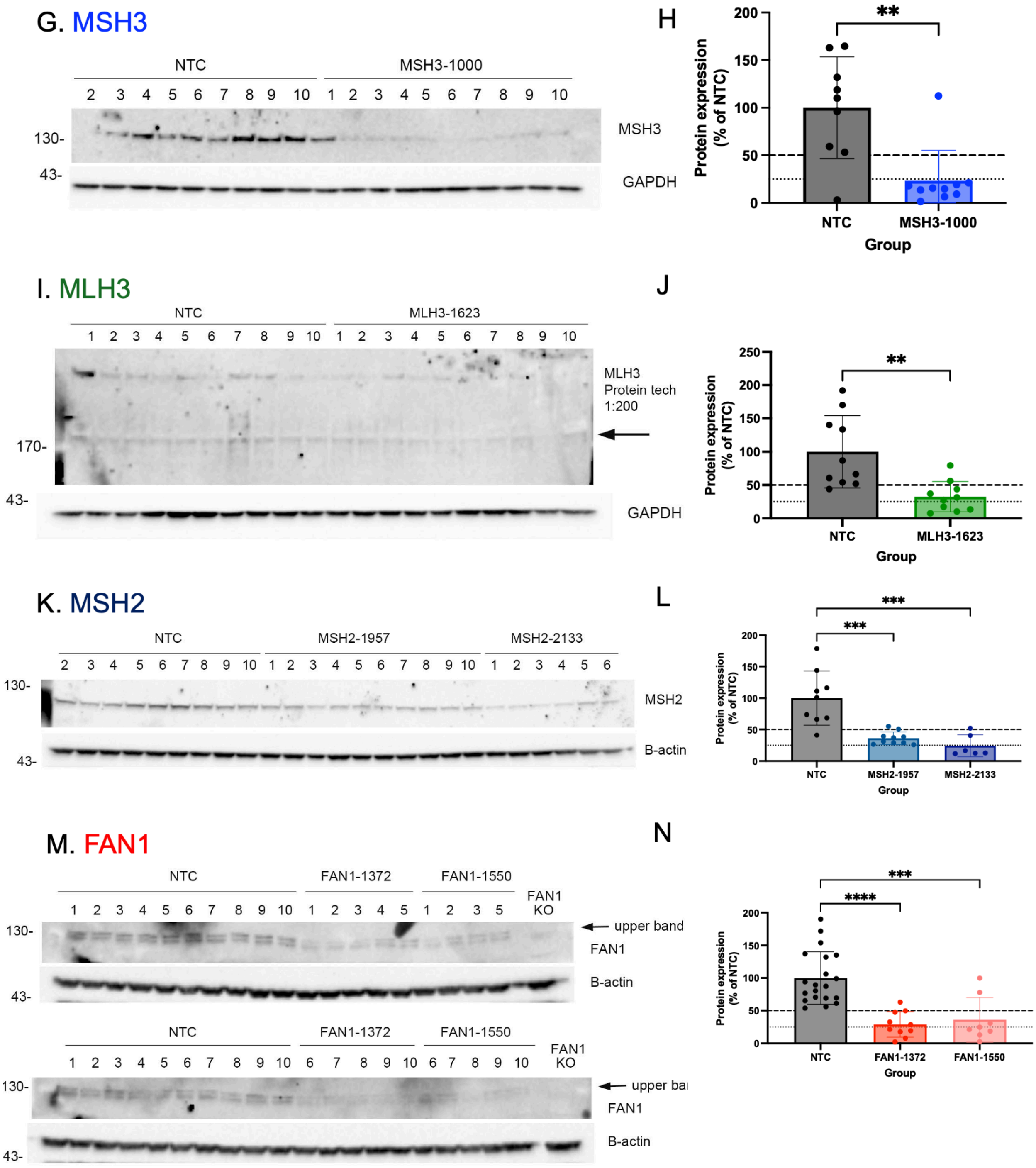

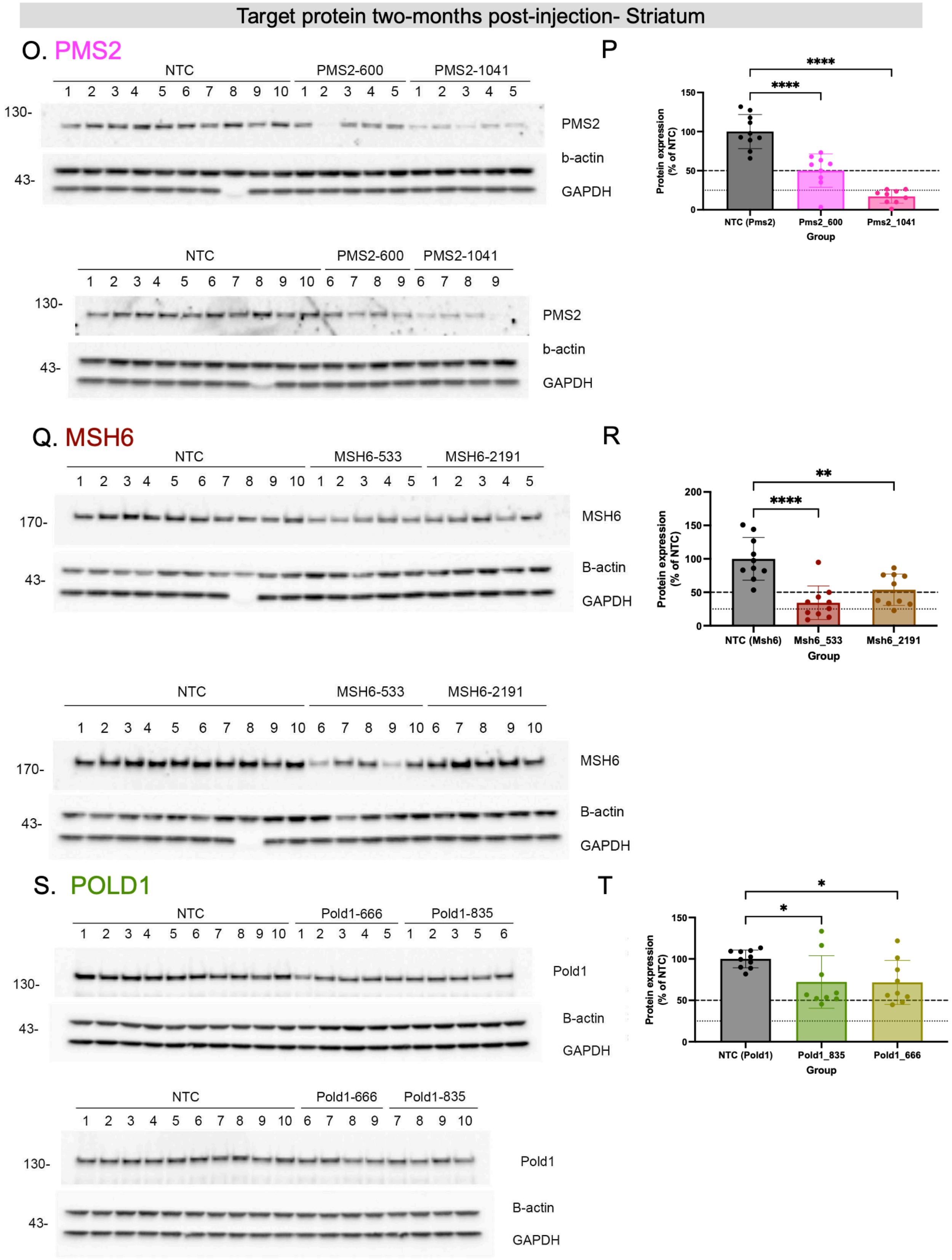

**Figure S8. Protein levels in Q111 mouse striatum two months post-injection with mismatch repair**

**pathway-targeting divalent siRNA.** (A-T) Raw and quantified protein levels in striatum two months post-siRNA treatment for each MMR target. (A) Western blot probing against MSH3 (B) quantification of MSH3 protein levels normalized to loading control and NTC. (C) Western blot probing against MLH1 (D) quantification of MLH1 protein levels normalized to loading control and NTC. (E) Western blot probing against POLD3 (F) quantification of POLD3 protein levels normalized to loading control and NTC. (G) Western blot probing against MSH3 (Study 2) (H) quantification of MSH3 protein levels normalized to loading control and NTC (study 2) (I) Western blot probing against MLH3 (J) quantification of MLH3 protein levels normalized to loading control and NTC. (K) Western blot probing against MSH2 (L) quantification of MSH2 protein levels normalized to loading control and NTC. (M) Western blot probing against FAN1 (N) quantification of FAN1 protein levels normalized to loading control and NTC. (O) Western blot probing against PMS2 (P) quantification of PMS2 protein levels normalized to loading control and NTC. (Q) Western blot probing against MSH6 (R) quantification of MSH6 protein levels normalized to loading control and NTC. (S) Western blot probing against POLD1 (T) quantification of POLD1 protein levels normalized to loading control and NTC. Each dot is the protein levels from a single mouse. N= 6-10/group. GAPDH and/or Beta-actin used as loading control. Statistics are one-way ANOVA versus NTC. * $p < 0.05$, ** $p < 0.01$, *** $p < 0.001$, **** $p < 0.0001$.

Target protein two-months post-injection- Medial Cortex

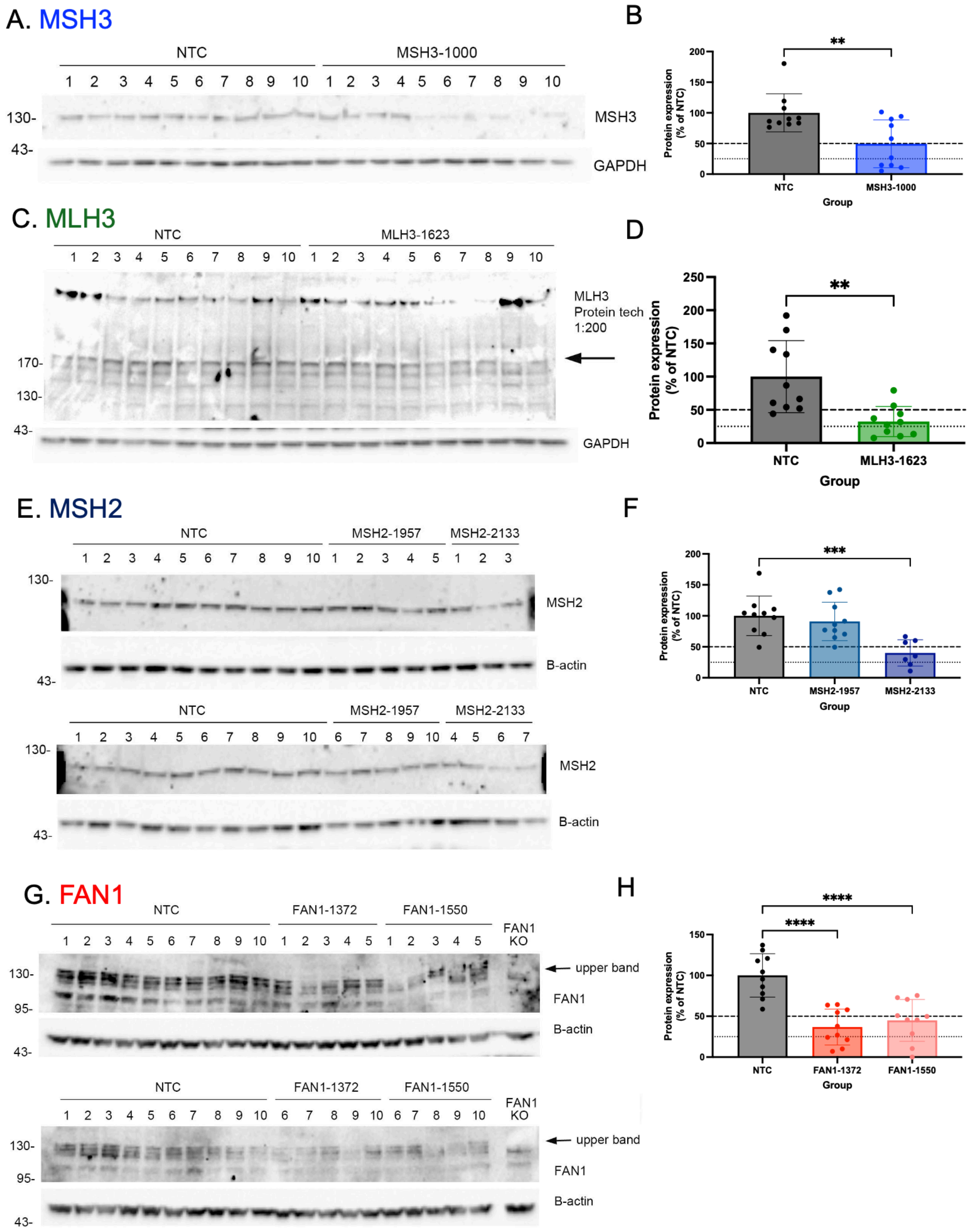

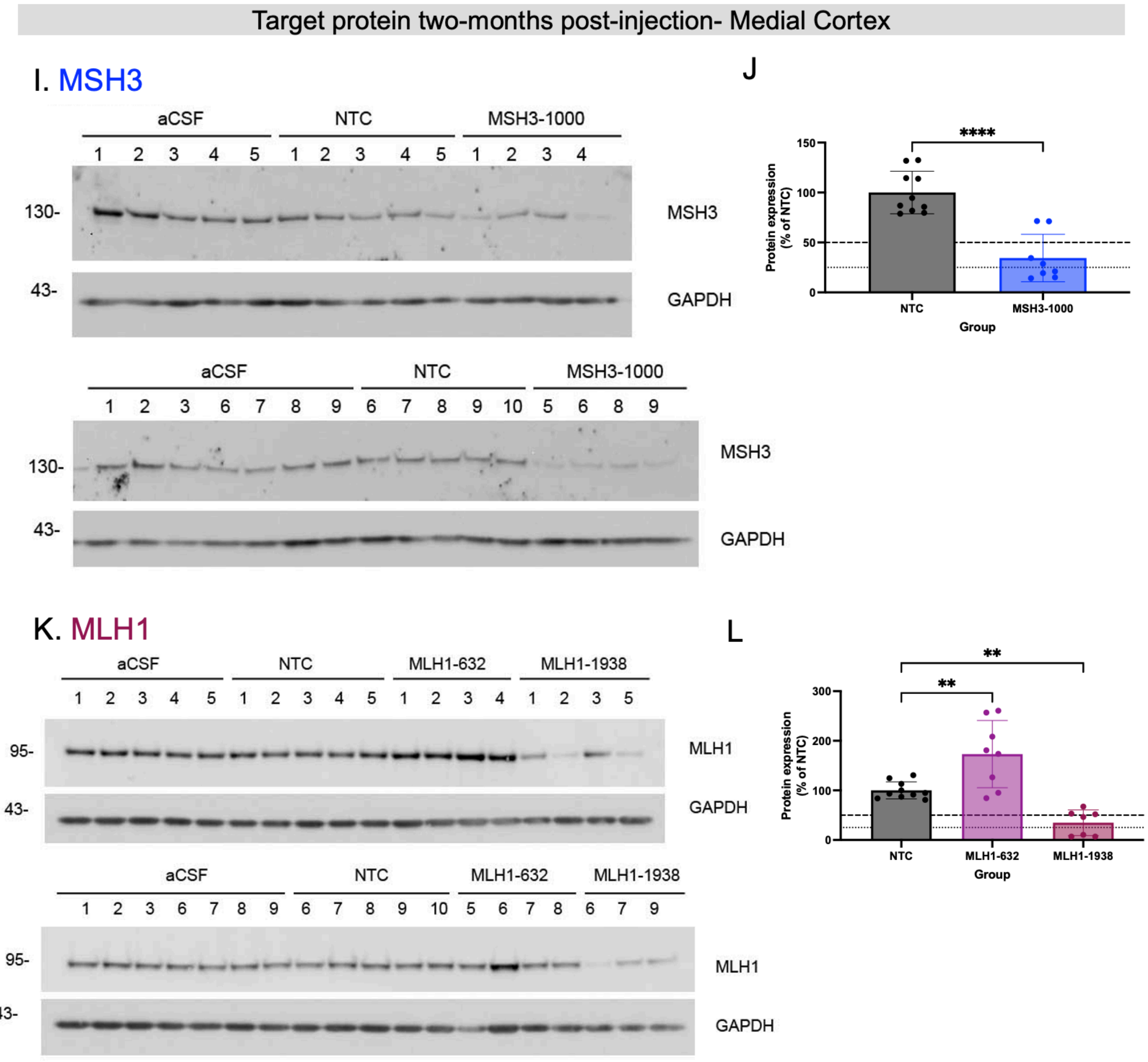
Target protein two-months post-injection- Medial Cortex
I. MSH3
aCSF
NTC
MSH3-1000
1 2 3 4 5 1 2 3 4 5 1 2 3 4
130-
MSH3
43-
GAPDH
aCSF
NTC
MSH3-1000
1 2 3 6 7 8 9 6 7 8 9 10 5 6 8 9
130-
MSH3
43-
GAPDH
J
****
Protein expression (% of NTC)
150
100
50
0
NTC
MSH3-1000
Group
K. MLH1
aCSF
NTC
MLH1-632
MLH1-1938
1 2 3 4 5 1 2 3 4 5 1 2 3 4 1 2 3 5
95-
MLH1
43-
GAPDH
aCSF
NTC
MLH1-632
MLH1-1938
1 2 3 6 7 8 9 6 7 8 9 10 5 6 7 8 6 7 9
95-
MLH1
43-
GAPDH
L
**
**
Protein expression (% of NTC)
300
200
100
0
NTC
MLH1-632
MLH1-1938
Group

Target protein two-months post-injection- Medial Cortex

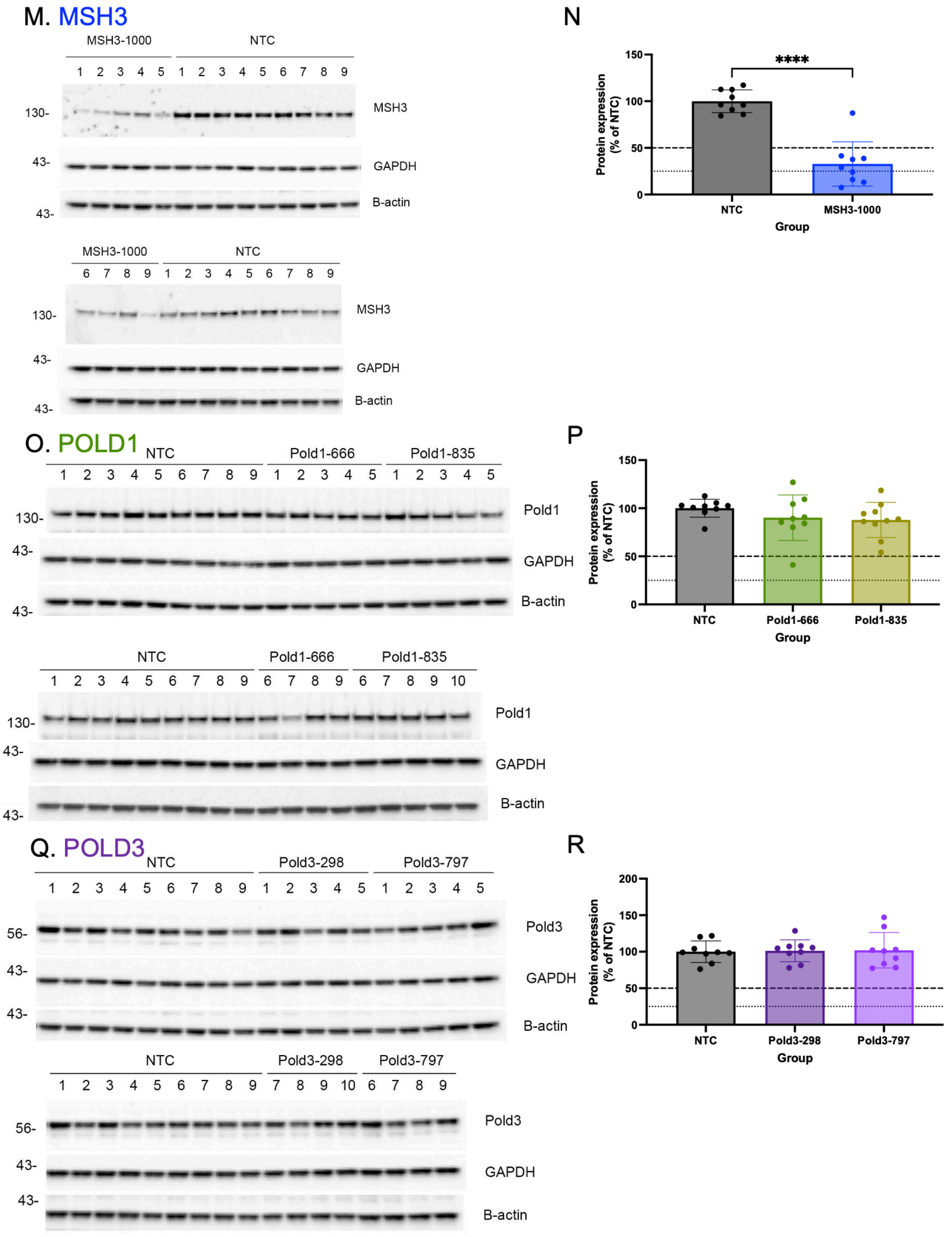

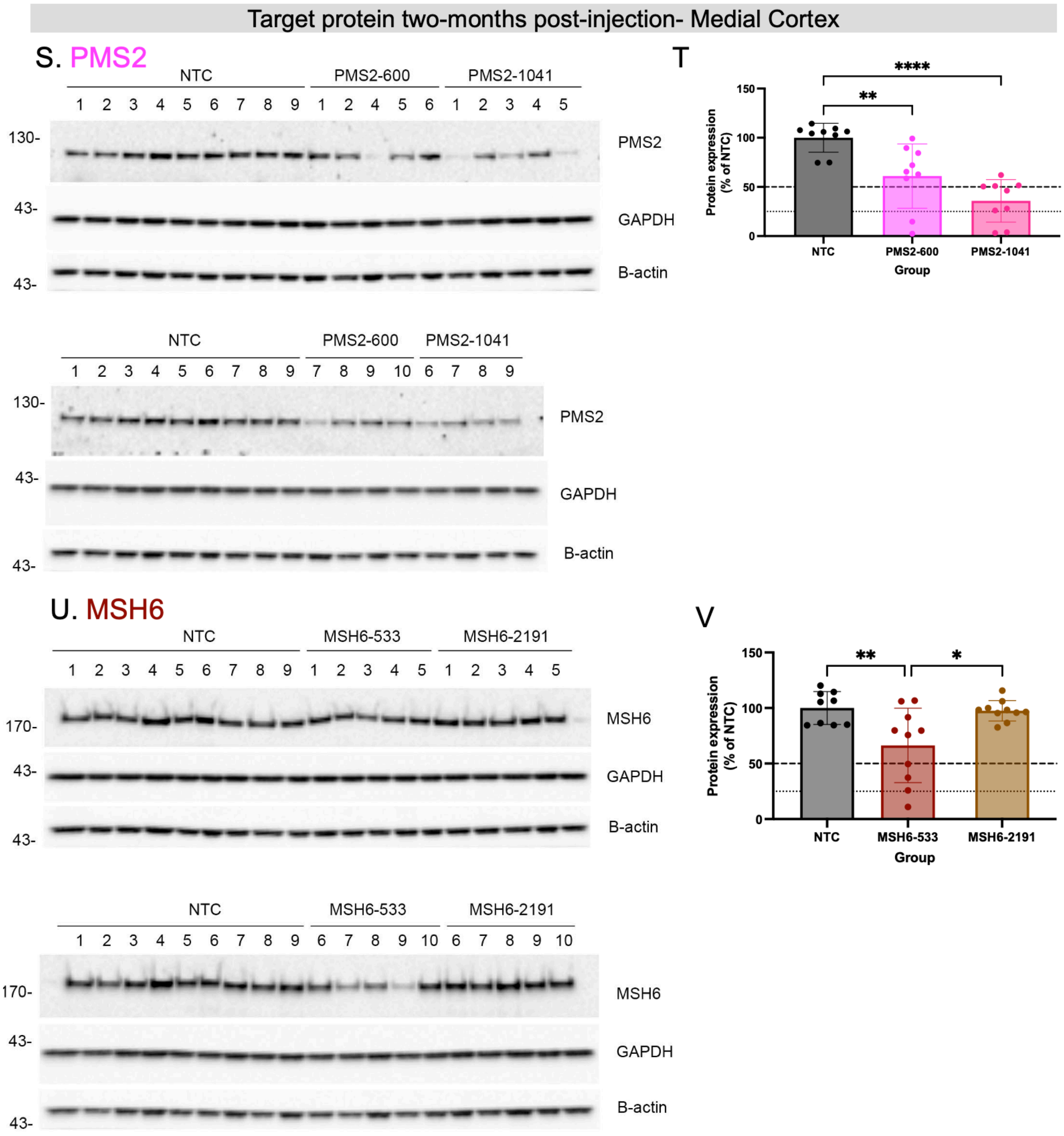

**Figure S9. Protein levels in Q111 mouse medial cortex two months post-injection with mismatch repair pathway-targeting divalent siRNA.** (A-V) Raw and quantified protein levels in medial cortex two months post-siRNA treatment for each MMR target. (A) Western blot probing against MSH3 (B) quantification of MSH3 protein levels normalized to loading control and NTC. **(**C) Western blot probing against MLH3 (D) quantification of MLH3 protein levels normalized to loading control and NTC. (E)

Western blot probing against MSH2 (F) quantification of MSH2 protein levels normalized to loading control and NTC. (G) Western blot probing against Fan1 (H) quantification of FAN1 protein levels normalized to loading control and NTC (I) Western blot probing against MSH3 (study 2) (J) quantification of MSH3 (study 2) protein levels normalized to loading control and NTC. (K) Western blot probing against MLH1 (L) quantification of MLH1 protein levels normalized to loading control and NTC. (M) Western blot probing against MSH3 (study 3) (N) quantification of MSH3 protein levels normalized to loading control and NTC. (O) Western blot probing against POLD1 (P) quantification of POLD1 protein levels normalized to loading control and NTC. (Q) Western blot probing against POLD3 (R) quantification of MSH6 protein levels normalized to loading control and NTC. (S) Western blot probing against PMS2 (T) quantification of PMS2 protein levels normalized to loading control and NTC. (U) Western blot probing against MSH6 (V) quantification of MSH6 protein levels normalized to loading control and NTC. Each dot is the protein level from a single mouse. N = 6-10/group. GAPDH and/or Beta-actin used as loading control. Statistics are one-way ANOVA versus NTC. * $p < 0.05$, ** $p < 0.01$, *** $p < 0.001$, **** $p < 0.0001$.

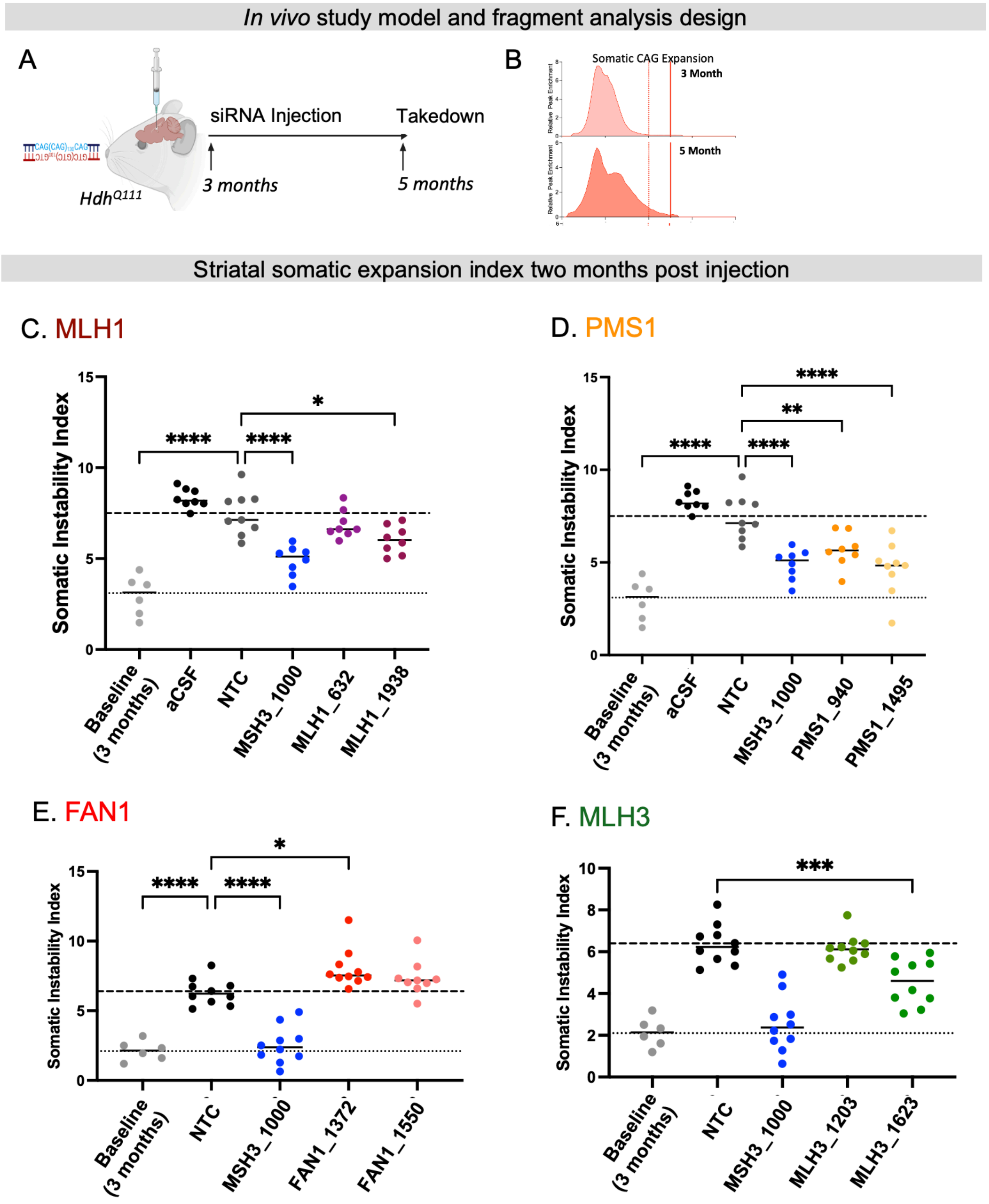
In vivo study model and fragment analysis design
A
CAG(CAG)CAG
GTC(GTC)GTC
HdhQ111
siRNA Injection
Takedown
3 months
5 months
B
Somatic CAG Expansion
3 Month
5 Month
Relative Peak Enrichment
Striatal somatic expansion index two months post injection
C. MLH1
Somatic Instability Index
Baseline (3 months)
aCSF
NTC
MSH3_1000
MLH1_632
MLH1_1938
D. PMS1
Somatic Instability Index
Baseline (3 months)
aCSF
NTC
MSH3_1000
PMS1_940
PMS1_1495
E. FAN1
Somatic Instability Index
Baseline (3 months)
NTC
MSH3_1000
FAN1_1372
FAN1_1550
F. MLH3
Somatic Instability Index
Baseline (3 months)
NTC
MSH3_1000
MLH3_1203
MLH3_1623

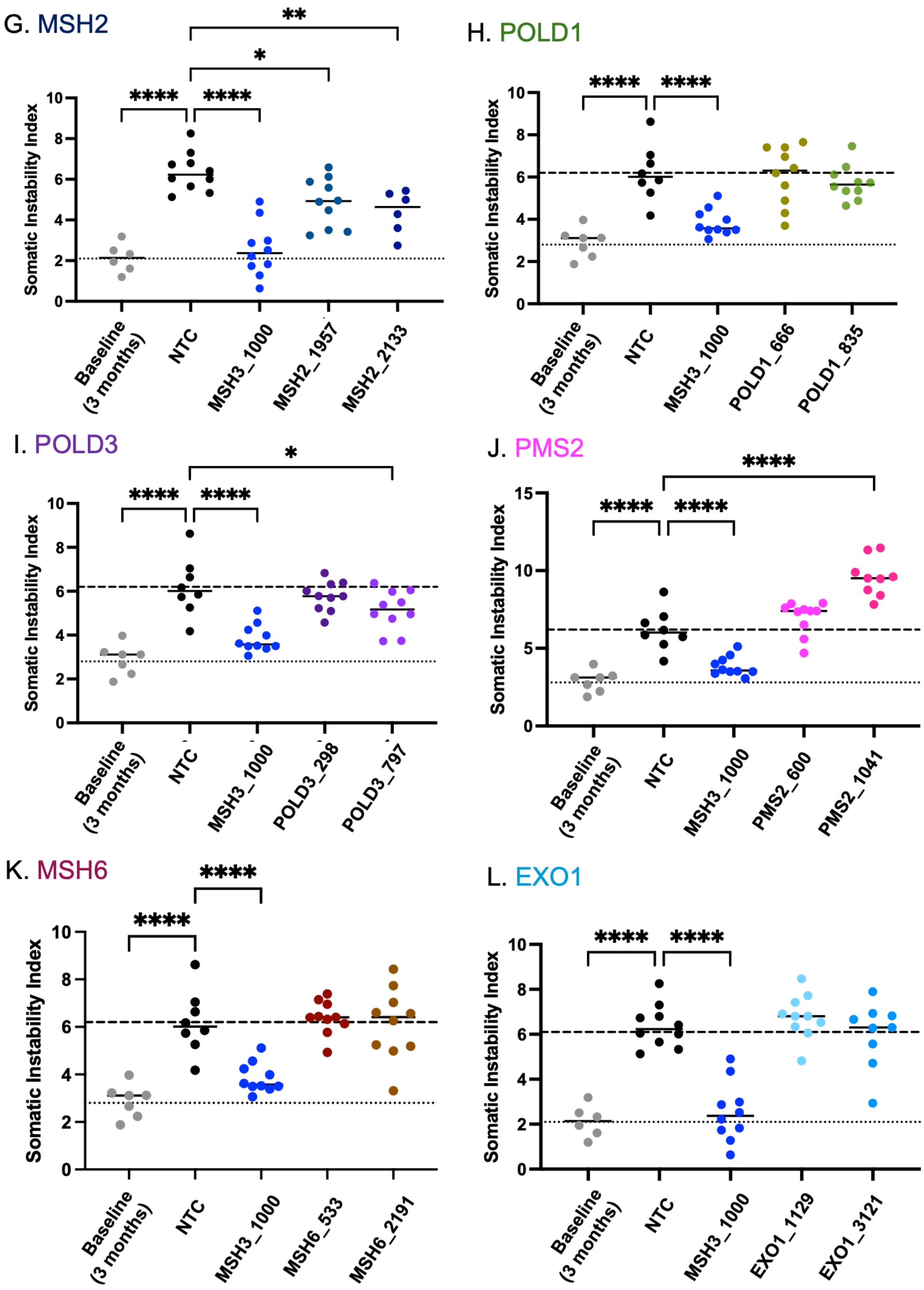
G. MSH2
Somatic Instability Index
Baseline (3 months)
NTC
MSH3_1000
MSH2_1957
MSH2_2133
****
*
**
H. POLD1
Somatic Instability Index
Baseline (3 months)
NTC
MSH3_1000
POLD1_666
POLD1_835
I. POLD3
Somatic Instability Index
Baseline (3 months)
NTC
MSH3_1000
POLD3_298
POLD3_797
J. PMS2
Somatic Instability Index
Baseline (3 months)
NTC
MSH3_1000
PMS2_600
PMS2_1041
K. MSH6
Somatic Instability Index
Baseline (3 months)
NTC
MSH3_1000
MSH6_533
MSH6_2191
L. EXO1
Somatic Instability Index
Baseline (3 months)
NTC
MSH3_1000
EXO1_1129
EXO1_3121

**Figure S10. Somatic instability index in the striatum of Q111 at age 5 months, two months post-injection with MMR-targeted divalent siRNA.** (A) Experimental paradigm and (B) representative fragment analysis used to quantify somatic instability index. (C-L) Baseline is striatal somatic instability of an untreated Q111 cohort at the time of injection (age 3 months). Non-targeting control (NTC), MSH3 controls and target siRNA are striatal somatic instability index of Q111 mice two-months post-injection (age 5 months). Somatic instability after (C) MLH1 modulation with MLH1_632, MLH1_1938 (D) PMS1 modulation PMS1_940, PMS1_1495 (E) FAN1 modulation with FAN1_1372, FAN1_1550 (F) MLH3 modulation with MLH3_1203, MLH3_1623 (G) MSH2 modulation with MSH2_1957, MSH2_2133 (H) POLD1 modulation with POLD1_666, POLD1_835 (I) POLD3 modulation with POLD3_298, POLD3_797 (J) PMS2 modulation with PMS2_600, PMS2_1041 (K) MSH6 modulation with MSH6_533, MSH6_2191 (L) EXO1 modulation, EXO1_1129, EXO1_1321. Fragment analysis quantified using somatic instability index. Each dot is the somatic instability index quantified from one mouse. Light dash line represents average baseline ("BL") somatic instability at the time of injection (age 3 months). Dark dash line represents average NTC somatic instability at the end of the experiment (age 5 months). aCSF is artificial cerebrospinal fluid vehicle buffer. N = 6-10/group. Statistics are one-way ANOVA vs NTC. * $p < 0.05$, ** $p < 0.01$, *** $p < 0.001$, **** $p < 0.0001$.

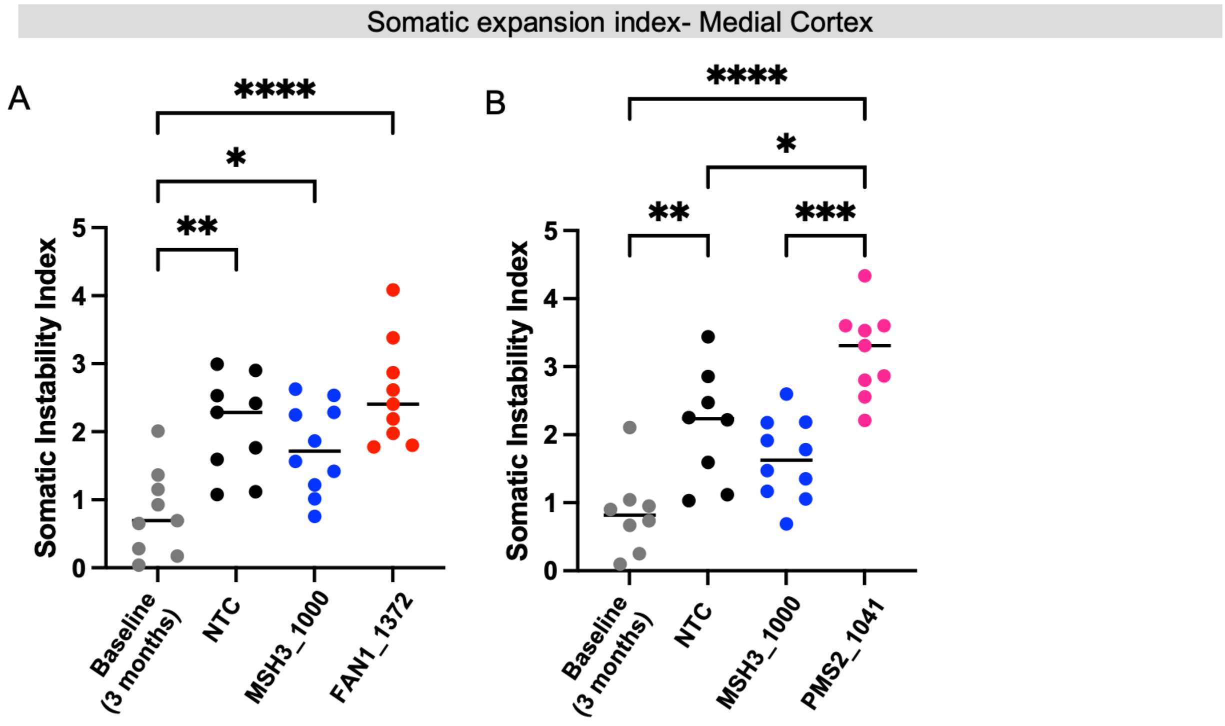

**Figure S11. Somatic instability in the medial cortex with select expansion-enhancing agents. (A-B)** Baseline is motor/medial cortex somatic instability of an untreated Q111 cohort at the time of injection (age 3 months). Non-targeting control (NTC), MSH3 controls, and target siRNA are striatal somatic instability index of Q111 mice two months post-injection (age 5 months). (A) Somatic instability in the cortex of Q111 mice treated with NTC, MSH3 or FAN1_1372 (B) Somatic instability in the cortex of Q111 mice treated with NTC, MSH3 or PMS2_1041. Statistics are one-way ANOVA with multiple comparisons.

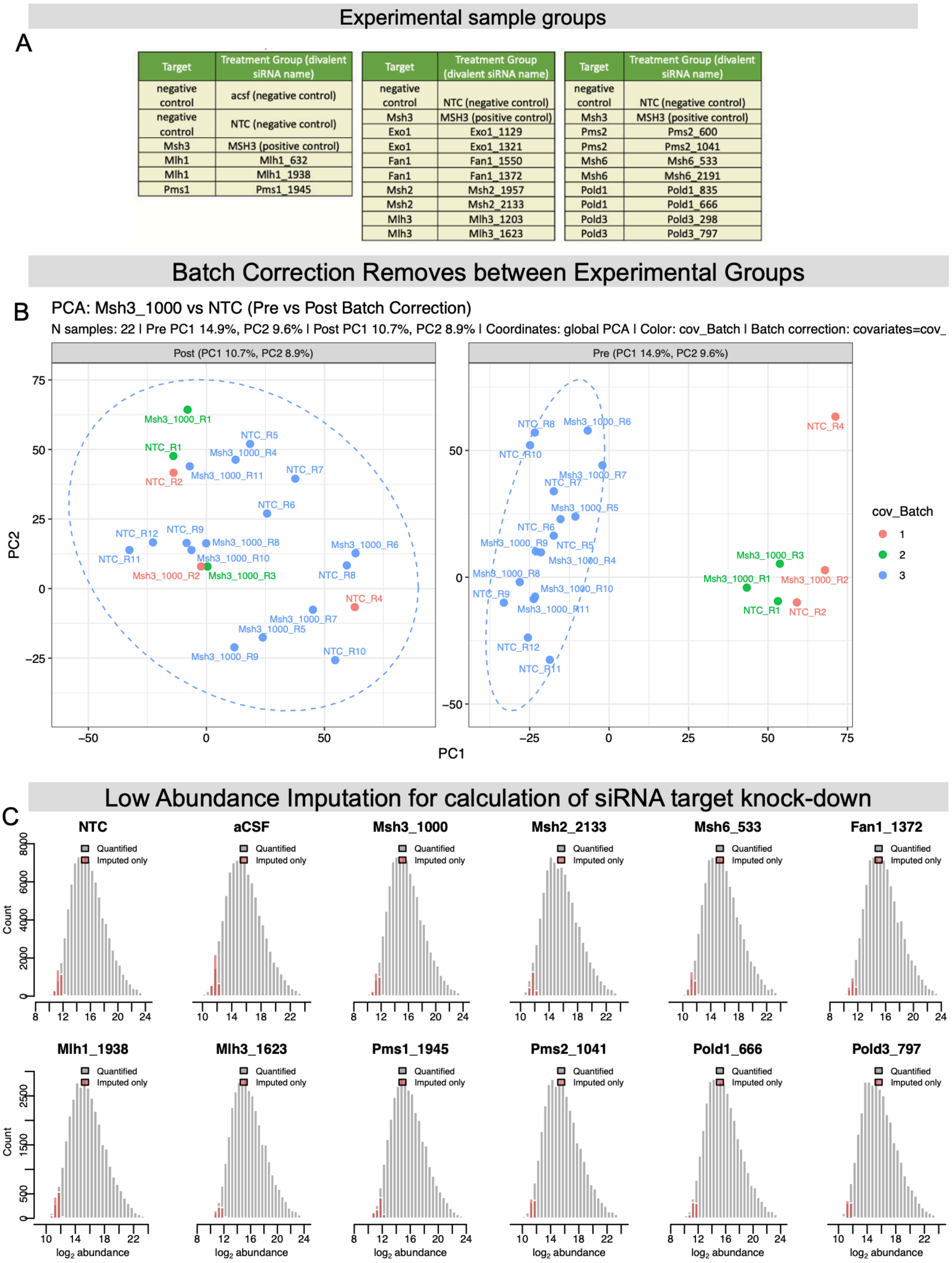

| Target | Treatment Group (divalent siRNA name) |
|---|---|
| negative control | acsf (negative control) |
| negative control | NTC (negative control) |
| Msh3 | MSH3 (positive control) |
| Mlh1 | Mlh1_632 |
| Mlh1 | Mlh1_1938 |
| Pms1 | Pms1_1945 |

| Target | Treatment Group (divalent siRNA name) |
|---|---|
| negative control | NTC (negative control) |
| Msh3 | MSH3 (positive control) |
| Exo1 | Exo1_1129 |
| Exo1 | Exo1_1321 |
| Fan1 | Fan1_1550 |
| Fan1 | Fan1_1372 |
| Msh2 | Msh2_1957 |
| Msh2 | Msh2_2133 |
| Mlh3 | Mlh3_1203 |
| Mlh3 | Mlh3_1623 |

| Target | Treatment Group (divalent siRNA name) |
|---|---|
| negative control | NTC (negative control) |
| Msh3 | MSH3 (positive control) |
| Pms2 | Pms2_600 |
| Pms2 | Pms2_1041 |
| Msh6 | Msh6_533 |
| Msh6 | Msh6_2191 |
| Pold1 | Pold1_835 |
| Pold1 | Pold1_666 |
| Pold3 | Pold3_298 |
| Pold3 | Pold3_797 |

**Figure S12. Proteomic analysis experimental grouping and quality control analysis** (A) Sample processing analysis groups (B) Representative Principal Component Analyses of NTC and MSH3 samples, N = 11 per group, showing batch effects (right) in PC1, which were mititgated significantly after batch correction (left) (C) Distribution of average log2 protein abundance values (gray bars) overlaid with low abundance imputed values (red bars), which were used for calculation of siRNA target knockdown.

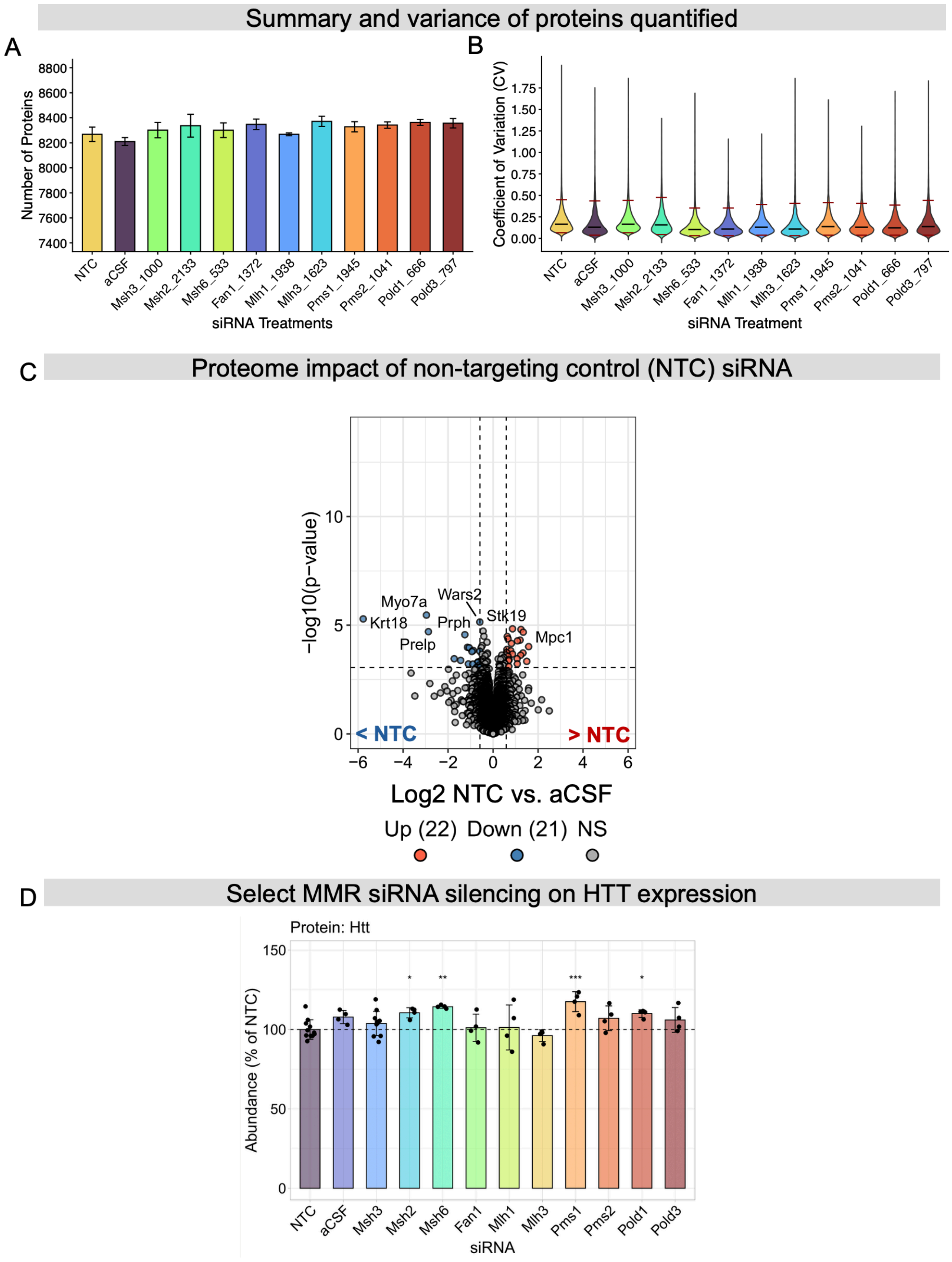

**Figure S13. Mass spectrometry and experimental quality control metrics.** (A) Proteins quantified across siRNA treatments. (B) Variance distribution of protein abundance across samples. (C) Volcano plots showing proteome abundance differences between non-targeting control vehicle and artificial CSF groups. Horizontal and vertical dashed lines indicate differential significance thresholds for p-value (10% FDR) and log2 fold-change, respectively. (D) Proteome abundances of HTT across siRNA treatment groups. N = 4/group, except for NTC and Msh3, which were N = 11. Statistics were calculated vs NTC using limma-based modeling to compute FDR-corrected p-values (* $p < 0.1$, ** $p < 0.01$, ** $p < 0.001$), as described in the Methods.

A PMS1 knockdown (Q111 mice, siRNA) vs. PMS1 homozygous KO (Q140 mice)

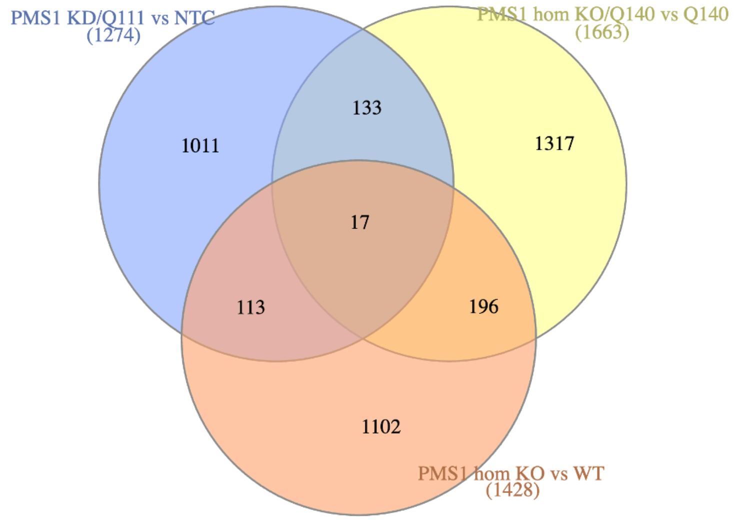

B PMS1-dependent transcriptome and proteome targets – STRING assembly

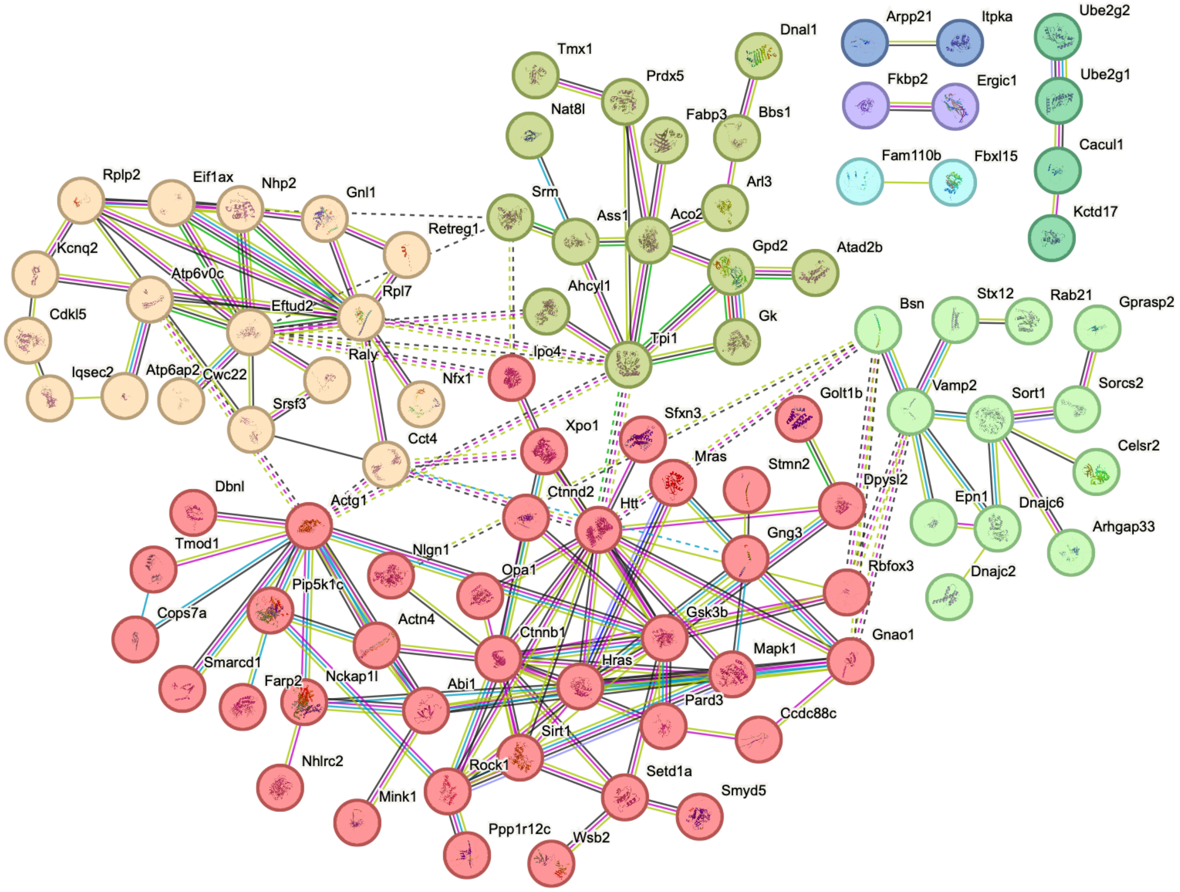

**Figure S14: Comparison of PMS1 knockdown and PMS1 knockout.** (A) Overlap of DE proteins (FDR < 0.1) in striatum of PMS1 KD Q111 mice (5m) compared with DE genes (FDR < 0.1) in striatum of PMS1 homozygous KO in WT and Q140 mice (6m) (Wang et al. Cell, 2025). No fold-change thresholds were applied to filter DE genes/proteins (B) STRING network of 133 shared DE targets between PMS1 KD and PMS1 KO in HD background. Node were color-coded in STRING by k-means clustering.

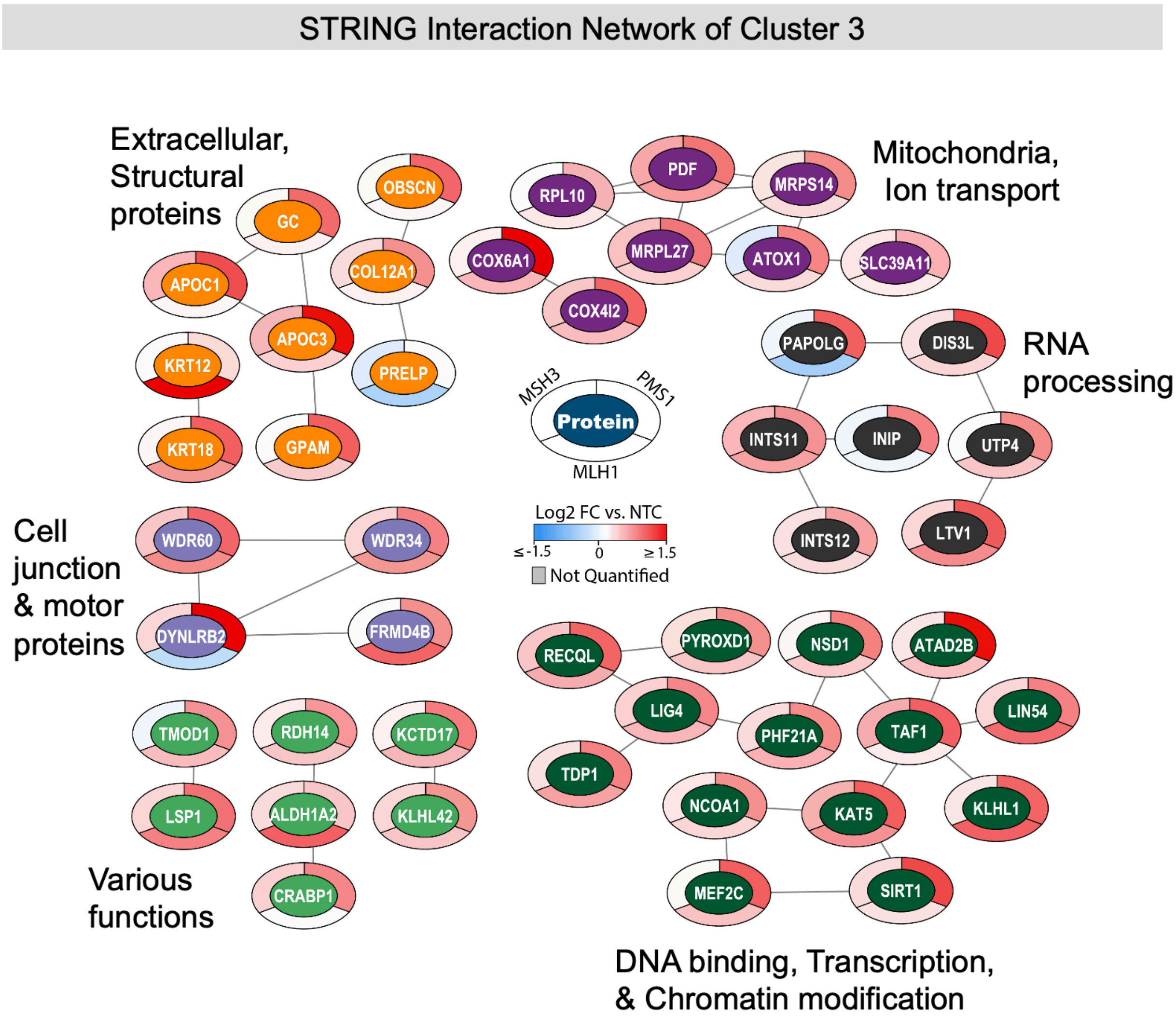

**Figure S15. STRING functional network of Cluster 3 proteins.** STRING network analysis was performed on the differential proteins assigned to cluster 3, the majority of which represented preferential up-regulation in PMS1 silencing (see Figure 6A & B). Mouse proteins were mapped to human identifiers and assembled with the STRING database default settings. Shown here are the 49 out of 121 proteins that had at least one scored interaction in STRING (score > 0.4). Donut graphs compare the effect of MSH3, PMS1, and MLH1 silencing versus NTC, represented by log2 fold-change (N = 4/group., except MSH3 and NTC which were N = 11). Statistics were calculated vs NTC using limma-based modeling to compute FDR-corrected p-values (* $p < 0.1$, ** $p < 0.01$, ** $p < 0.001$), as described in the Methods. Node fill colors were manually assigned by interconnectivity and/or shared cellular functions.

| **Target** | **siRNA name** | **Homology** | **siRNA antisense sequence and chemical modificaiton pattern** | **siRNA sense sequence and chemical modificaiton pattern** |
|---|---|---|---|---|
| *MSH3* | MSH3_1000 | Mouse, Human | P(mU)#(fG)#(mC)(fA)(fG)(fU)(mU)(fU)(mC)(fA)(mG)(fU)(mU)(fU)#(mG)#(fC)#(mU)#(mU)#(mC)#(fA)#(mU) | (mG)#(mC)#(mA)(fA)(mA)(fC)(mU)(fG)(mA)(fA)(mA)(mC)(mU)(fG)#(mC)#(mA)-TegChol |
| *MLH1* | Mlh1_1938 | Mouse, Human | P(mU)#(fC)#(mA)(fU)(fA)(fA)(mA)(fU)(mG)(fA)(mG)(fU)(mA)(fU)#(mC)#(fU)#(mG)#(mG)#(mU)#(fA)#(mU) | (mA)#(mG)#(mA)(fU)(mA)(fC)(mU)(fC)(mA)(fU)(mU)(mU)(mA)(fU)#(mG)#(mA)-TegChol |
| *Mlh3* | Mlh3_1203 | Mouse | P(mU)#(fU)#(mA)(fA)(fA)(fU)(mU)(fC)(mC)(fU)(mU)(fA)(mA)(fU)#(mA)#(fU)#(mC)#(mU)#(mU)#(fC)#(mU) | (mA)#(mU)#(mA)(fU)(mU)(fA)(mA)(fG)(mG)(fA)(mA)(mU)(mU)(fU)#(mA)#(mA)-TegChol |
| *MLH3* | MLH3_1251 | Human | P(mU)#(fA)#(mA)(fA)(fC)(fU)(mA)(fA)(mA)(fA)(mC)(fC)(mA)(fU)#(mU)#(fA)#(mU)#(mC)#(mU)#(fU)#(mU) | (mU)#(mA)#(mA)(fU)(mG)(fG)(mU)(fU)(mU)(fU)(mA)(mG)(mU)(fU)#(mU)#(mA)-TegChol |
| *PMS1* | PMS1_940 | Mouse, Human | P(mU)#(fU)#(mA)(fC)(fA)(fG)(mU)(fU)(mG)(fU)(mA)(fC)(mC)(fU)#(mU)#(fG)#(mA)#(mC)#(mC)#(fA)#(mU) | (mC)#(mA)#(mA)(fG)(mG)(fU)(mA)(fC)(mA)(fA)(mC)(mU)(mG)(fU)#(mA)#(mA)-TegChol |
| *Fan1* | Fan1_1372 | Mouse | P(mU)#(fA)#(mC)(fC)(fA)(fU)(mU)(fU)(mG)(fA)(mU)(fU)(mC)(fU)#(mG)#(fC)#(mC)#(mU)#(mC)#(fC)#(mU) | (mG)#(mC)#(mA)(fG)(mA)(fA)(mU)(fC)(mA)(fA)(mA)(mU)(mG)(fG)#(mU)#(mA)-TegChol |
| *FAN1* | FAN1_1647 | Human | P(mU)#(fU)#(mG)(fU)(fC)(fU)(mG)(fU)(mA)(fG)(mA)(fA)(mA)(fG)#(mC)#(fC)#(mU)#(mG)#(mC)#(fA)#(mU) | (mG)#(mG)#(mC)(fU)(mU)(fU)(mC)(fU)(mA)(fC)(mA)(mG)(mA)(fC)#(mA)#(mA)-TegChol |

| | | | | |
|---|---|---|---|---|
| *Pms2* | Pms2_1041 | Mouse, Human | P(mU)#(fA)#(mA)(fU)(fU)(fU)(mG)(fC)(mC)(fU)(mU)(fU)(mU)(fA)#(mU)#(fC)#(mU)#(mG)#(mG)#(fA)#(mU) | (mG)#(mA)#(mU)(fA)(mA)(fA)(mA)(fG)(mG)(fC)(mA)(mA)(mA)(fU)#(mU)#(mA)-TegChol |
| *PMS2* | PMS2_3337 | Human | P(mU)#(fA)#(mG)(fC)(fC)(fC)(mU)(fG)(mA)(fU)(mU)(fC)(mA)(fC)#(mA)#(fU)#(mU)#(mA)#(mA)#(fA)#(mU) | (mA)#(mU)#(mG)(fU)(mG)(fA)(mA)(fU)(mC)(fA)(mG)(mG)(mG)(fC)#(mU)#(mA)-TegChol |
| *EXO1* | EXO1_1321 | Mouse, Human | P(mU)#(fA)#(mU)(fU)(fU)(fU)(mC)(fU)(mU)(fG)(mA)(fU)(mA)(fA)#(mC)#(fC)#(mU)#(mU)#(mU)#(fA)#(mU) | (mG)#(mG)#(mU)(fU)(mA)(fU)(mC)(fA)(mA)(fG)(mA)(mA)(mA)(fA)#(mU)#(mA)-TegChol |
| *MSH2* | MSH2_2133 | Mouse, Human | P(mU)#(fG)#(mC)(fA)(fG)(fA)(mA)(fG)(mU)(fG)(mU)(fC)(mC)(fA)#(mU)#(fU)#(mG)#(mU)#(mG)#(fG)#(mU) | (mA)#(mA)#(mU)(fG)(mG)(fA)(mC)(fA)(mC)(fU)(mU)(mC)(mU)(fG)#(mC)#(mA)-TegChol |
| *Msh6* | Msh6_533 | Mouse, Human | P(mU)#(fU)#(mG)(fA)(fA)(fC)(mC)(fU)(mG)(fU)(mA)(fU)(mA)(fU)#(mG)#(fG)#(mC)#(mU)#(mU)#(fU)#(mU) | (mC)#(mC)#(mA)(fU)(mA)(fU)(mA)(fC)(mA)(fG)(mG)(mU)(mU)(fC)#(mA)#(mA)-TegChol |
| *MSH6* | MSH6_2706 | Human | P(mU)#(fC)#(mA)(fA)(fC)(fU)(mU)(fC)(mU)(fU)(mC)(fC)(mA)(fU)#(mG)#(fA)#(mU)#(mC)#(mC)#(fC)#(mU) | (mU)#(mC)#(mA)(fU)(mG)(fG)(mA)(fA)(mG)(fA)(mA)(mG)(mU)(fU)#(mG)#(mA)-TegChol |
| *Pold1* | Pold1_666 | Mouse | P(mU)#(fA)#(mG)(fA)(fA)(fA)(mU)(fG)(mG)(fA)(mG)(fA)(mA)(fG)#(mG)#(fG)#(mC)#(mC)#(mA)#(fU)#(mU) | (mC)#(mC)#(mC)(fU)(mU)(fC)(mU)(fC)(mC)(fA)(mU)(mU)(mU)(fC)#(mU)#(mA)-TegChol |
| *POLD1* | POLD1_1885 | Human | P(mU)#(fU)#(mG)(fG)(fU)(fG)(mU)(fA)(mA)(fC)(mA)(fC)(mA)(fG)#(mG)#(fU)#(mU)#(mG)#(mU)#(fG)#(mU) | (mA)#(mC)#(mC)(fU)(mG)(fU)(mG)(fU)(mU)(fA)(mC)(mA)(mC)(fC)#(mA)#(mA)-TegChol |
| *POLD3* | POLD3_298 | Mouse, Human | P(mU)#(fA)#(mC)(fU)(fA)(fC)(mU)(fG)(mC)(fA)(mA)(fC)(mC)(fU)#(mU)#(fG)#(mU)#(mG)#(mG)#(fC)#(mU) | (mC)#(mA)#(mA)(fG)(mG)(fU)(mU)(fG)(mC)(fA)(mG)(mU)(mA)(fG)#(mU)#(mA)-TegChol |

| |
|---|
| *Modifications key: P- phosphate, m- 2'OMe, f- 2'F (fluoro), #- phophorotioate, TegChol- cholesterol* |

**Table S1:** *In vitro* validated siRNA for mouse and human modulation of the mismatch repair pathway

| siRNA sense sequence and chemical modificaiton pattern | | | | |
|---|---|---|---|---|
| **Target** | **siRNA name** | **Homology** | **Antisense** | **Sense** |
| Non targeting control | NTC | Human, Mouse | V(mU)#(fA)#(mA)(fU)(fC)(fG)(mU)(fA)(mU)(fU)(mU)(fG)(mU)(fC)#(mA)#(fA)#(mU)#(mC)#(mA)#(fU)#(mU) | (mU)#(mU)#(mG)(fA)(mC)(fA)(mA)(fA)(mU)(fA)(mC)(mG)(mA)(fU)#(mU)#(mA)-DIO |
| Msh3 | MSH3_1000 | Human, Mouse | V(mU)#(fG)#(mC)(fA)(fG)(fU)(mU)(fU)(mC)(fA)(mG)(fU)(mU)(fU)#(mG)#(fC)#(mU)#(mU)#(mC)#(fA)#(mU) | (mG)#(mC)#(mA)(fA)(mA)(fC)(mU)(fG)(mA)(fA)(mA)(mC)(mU)(fG)#(mC)#(mA)-DIO |
| Mlh1 | MLH1_1938 | Human, Mouse | V(mU)#(fC)#(mA)(fU)(fA)(fA)(mA)(fU)(mG)(fA)(mG)(fU)(mA)(fU)#(mC)#(fU)#(mG)#(mG)#(mU)#(fA)#(mU) | (mA)#(mG)#(mA)(fU)(mA)(fC)(mU)(fC)(mA)(fU)(mU)(mU)(mA)(fU)#(mG)#(mA)-DIO |
| Mlh1 | MLH1__632 | Mouse | V(mU)#(fU)#(mG)(fA)(fG)(fU)(mA)(fA)(mC)(fU)(mU)(fG)(mC)(fU)#(mC)#(fU)#(mG)#(mU)#(mA)#(fU)#(mU) | (mA)#(mG)#(mA)(fG)(mC)(fA)(mA)(fG)(mU)(fU)(mA)(mC)(mU)(fC)#(mA)#(mA)-DIO |
| Pms1 | PMS1_940 | Human, Mouse | V(mU)#(fU)#(mA)(fC)(fA)(fG)(mU)(fU)(mG)(fU)(mA)(fC)(mC)(fU)#(mU)#(fG)#(mA)#(mC)#(mC)#(fA)#(mU) | (mC)#(mA)#(mA)(fG)(mG)(fU)(mA)(fC)(mA)(fA)(mC)(mU)(mG)(fU)#(mA)#(mA)-DIO |
| Pms1 | PMS1_1495 | Human, Mouse | V(mU)#(fA)#(mG)(fC)(fA)(fA)(mU)(fU)(mA)(fA)(mA)(fA)(mC)(fA)#(mG)#(fA)#(mU)#(mU)#(mC)#(fC)#(mU) | (mU)#(mC)#(mU)(fG)(mU)(fU)(mU)(fU)(mA)(fA)(mU)(mU)(mG)(fC)#(mU)#(mA)-DIO |
| Pms2 | Pms2_600 | Mouse | V(mU)#(fA)#(mG)(fA)(fC)(fC)(mU)(fG)(mC)(fA)(mC)(fC)(mA)(fU)#(mU)#(fU)#(mU)#(mG)#(mG)#(fC)#(mU) | (mA)#(mA)#(mA)(fU)(mG)(fG)(mU)(fG)(mC)(fA)(mG)(mG)(mU)(fC)#(mU)#(mA)-DIO |
| Pms2 | PMS2_1041 | Human, Mouse | V(mU)#(fA)#(mA)(fU)(fU)(fU)(mG)(fC)(mC)(fU)(mU)(fU)(mU)(fA)#(mU)#(fC)#(mU)#(mG)#(mG)#(fA)#(mU) | (mG)#(mA)#(mU)(fA)(mA)(fA)(mA)(fG)(mG)(fC)(mA)(mA)(mA)(fU)#(mU)#(mA)-DIO |
| Pold1 | Pold1_666 | Mouse | V(mU)#(fA)#(mG)(fA)(fA)(fA)(mU)(fG | (mC)#(mC)#(mC)(fU)(mU)(fC)(mU)(f |

| | | | | |
|---|---|---|---|---|
| | | | )(mG)(fA)(mG)(fA)(mA)(fG)#(mG)#(fG)#(mC)#(mC)#(mA)#(fU)#(mU) | C)(mC)(fA)(mU)(mU)(mU)(fC)#(mU)#(mA)-DIO |
| Pold1 | Pold1_835 | Mouse | V(mU)#(fC)#(mA)(fA)(fC)(fC)(mA)(fG)(mU)(fU)(mG)(fC)(mA)(fU)#(mC)#(fC)#(mC)#(mA)#(mC)#(fA)#(mU) | (mG)#(mG)#(mA)(fU)(mG)(fC)(mA)(fA)(mC)(fU)(mG)(mG)(mU)(fU)#(mG)#(mA)-DIO |
| Msh6 | MSH6_533 | Human, Mouse | V(mU)#(fU)#(mG)(fA)(fA)(fC)(mC)(fU)(mG)(fU)(mA)(fU)(mA)(fU)#(mG)#(fG)#(mC)#(mU)#(mU)#(fU)#(mU) | (mC)#(mC)#(mA)(fU)(mA)(fU)(mA)(fC)(mA)(fG)(mG)(mU)(mU)(fC)#(mA)#(mA)-DIO |
| Msh6 | MSH6_2191 | Human, Mouse | V(mU)#(fU)#(mC)(fA)(fA)(fA)(mA)(fU)(mU)(fA)(mG)(fC)(mC)(fA)#(mU)#(fU)#(mG)#(mA)#(mU)#(fA)#(mU) | (mA)#(mA)#(mU)(fG)(mG)(fC)(mU)(fA)(mA)(fU)(mU)(mU)(mU)(fG)#(mA)#(mA)-DIO |
| Pold3 | Pold3_797 | Human, Mouse | V(mU)#(fU)#(mA)(fA)(fG)(fU)(mU)(fU)(mA)(fU)(mU)(fC)(mA)(fU)#(mU)#(fG)#(mC)#(mA)#(mG)#(fC)#(mU) | (mC)#(mA)#(mA)(fU)(mG)(fA)(mA)(fU)(mA)(fA)(mA)(mC)(mU)(fU)#(mA)#(mA)-DIO |
| Pold3 | Pold3_298 | Human, Mouse | V(mU)#(fA)#(mC)(fU)(fA)(fC)(mU)(fG)(mC)(fA)(mA)(fC)(mC)(fU)#(mU)#(fG)#(mU)#(mG)#(mG)#(fC)#(mU) | (mC)#(mA)#(mA)(fG)(mG)(fU)(mU)(fG)(mC)(fA)(mG)(mU)(mA)(fG)#(mU)#(mA)-DIO |
| Exo1 | Exo1_1129 | Human, Mouse | V(mU)#(fA)#(mG)(fU)(fC)(fC)(mA)(fU)(mU)(fU)(mC)(fC)(mA)(fA)#(mA)#(fC)#(mU)#(mG)#(mG)#(fU)#(mU) | (mG)#(mU)#(mU)(fU)(mG)(fG)(mA)(fA)(mA)(fU)(mG)(mG)(mA)(fC)#(mU)#(mA)-DIO |
| Exo1 | Exo1_1321 | Human, Mouse | V(mU)#(fA)#(mU)(fU)(fU)(fU)(mC)(fU)(mU)(fG)(mA)(fU)(mA)(fA)#(mC)#(fC)#(mU)#(mU)#(mU)#(fA)#(mU) | (mG)#(mG)#(mU)(fU)(mA)(fU)(mC)(fA)(mA)(fG)(mA)(mA)(mA)(fA)#(mU)#(mA)-DIO |
| Fan1 | Fan1_1550 | Human, Mouse | V(mU)#(fA)#(mA)(fU)(fU)(fU)(mA)(fC)(mG)(fU)(mU)(fG)(mA)(fA)#(mA)#(fG)#(mA)#(mG)#(mC)#(fC)#(mU) | (mC)#(mU)#(mU)(fU)(mC)(fA)(mA)(fC)(mG)(fU)(mA)(mA)(mA)(fU)#(mU)#(mA)-DIO |
| Fan1 | Fan1_1372 | Mouse | V(mU)#(fA)#(mC)(fC)(fA)(fU)(mU)(fU)(mG)(fA)(mU)(fU) | (mG)#(mC)#(mA)(fG)(mA)(fA)(mU)(fC)(mA)(fA)(mA)(m |

| | | | | |
|---|---|---|---|---|
| | | | (mC)(fU)#(mG)#(fC)#(mC)#(mU)#(mC)#(fC)#(mU) | U)(mG)(fG)#(mU)#(mA)-DIO |
| Msh2 | Msh2_1957 | Human, Mouse | V(mU)#(fC)#(mA)(fU)(fC)(fU)(mU)(fG)(mA)(fA)(mC)(fU)(mU)(fC)#(mA)#(fA)#(mC)#(mA)#(mC)#(fA)#(mU) | (mU)#(mU)#(mG)(fA)(mA)(fG)(mU)(fU)(mC)(fA)(mA)(mG)(mA)(fU)#(mG)#(mA)-DIO |
| Msh2 | Msh2_2133 | Human, Mouse | V(mU)#(fC)#(mA)(fC)(fA)(fA)(mU)(fG)(mG)(fA)(mC)(fA)(mC)(fU)#(mU)#(fC)#(mU)#(mG)#(mC)#(fU)#(mU) | (mG)#(mA)#(mA)(fG)(mU)(fG)(mU)(fC)(mC)(fA)(mU)(mU)(mG)(fU)#(mG)#(mA)-DIO |
| Mlh3 | Mlh3_1203 | Mouse | V(mU)#(fU)#(mA)(fA)(fA)(fU)(mU)(fC)(mC)(fU)(mU)(fA)(mA)(fU)#(mA)#(fU)#(mC)#(mU)#(mU)#(fC)#(mU) | (mA)#(mU)#(mA)(fU)(mU)(fA)(mA)(fG)(mG)(fA)(mA)(mU)(mU)(fU)#(mA)#(mA)-DIO |
| Mlh3 | Mlh3_1623 | Mouse | V(mU)#(fU)#(mU)(fU)(fC)(fA)(mU)(fG)(mU)(fA)(mU)(fC)(mC)(fU)#(mU)#(fC)#(mC)#(mU)#(mG)#(fA)#(mU) | (mG)#(mA)#(mA)(fG)(mG)(fA)(mU)(fA)(mC)(fA)(mU)(mG)(mA)(fA)#(mA)#(mA)-DIO |

**Table S2:** Top siRNA to target the human and mouse mismatch repair pathway evaluated *in vivo.*
*Modifications key: V- vinylphosphate, m- 2'OMe, f- 2'F (fluoro), #- phophorotioate, DIO- di-valent linker*

| Target | Antibody | Successfully used | Expected Band | Comment |
|---|---|---|---|---|
| MSH3 | **Santa Cruz sc-271079** | **Yes** | **127 kDa** | **Band around 130 kDa** |
| MSH2 | Cell Signaling 2017 | Yes | 100 kDa | Band around 100 kDa |
| MSH2 | Inv 33-7900 | No | 100 kDa | Band around 100 kDa; several other bands present |
| FAN1 | Univ Dundee | Yes | 114 kDa | Band around 110 kDa; disappears in Fan1 KO tissue |
| MSH6 | Inv MA5-42406 | Yes | 160 kDa | Band around 180-190 kDa |
| MLH1 | Abcam Ab92312 | Yes | 84 kDa | Band around 96kDa; disappears in Mlh1 KO tissue |
| MLH1 | Mybiosource MBS9605077 | No | 84 kDa | No change in bands in Mlh1 KO tissue |
| MLH1 | Santa Cruz sc-271978 | No | 84 kDa | No band at expected size |
| POLD1 | Inv MA5-35464 | Yes | 124 kDa | Band around 130 kDa |
| POLD3 | Inv PA5-96616 | Yes | 51-70 kDa | Mouse band ~56 kDa; human ~72 kDa |
| PMS2 | BD 556415 | Yes | 110 kDa | Band ~100 kDa; disappears in Pms2 KO tissue |
| PMS2 | Inv PA5-87127 | No | 110 kDa | No clear band at the expected MW |
| MLH3 | Inv PA5-75345 | No | 161 kDa | No clear band at the expected MW |
| MLH3 | Mybiosource MBS9605078 | No | 161 kDa | No clear band at the expected MW |
| MLH3 | Santa Cruz sc-25313 | No | 161 kDa | No clear band at the expected MW |
| MLH3 | Proteintech 25298-1-AP | Yes | 161 kDa | Saw band after Mlh1 IP but required significant amt of tissue |
| EXO1 | ProteinTech 16253-1-ap | No | 115 kDa | Signal in human neurons at expected MW, not in mouse brain |
| EXO1 | Abcam ab155553 | No | 115 kDa | 2-3 bands at expected MW in human neurons and mouse brain |
| PMS1 | Mybiosource MBS9134056 | No | 106 kDa | No unique band; no change in Pms1 KO tissue |

| | | | | |
|---|---|---|---|---|
| PMS1 | Thermo PA5-86724 | No | 106 kDa | No unique band; no change in Pms1 KO tissue |
| PMS1 | Thermo PA5-100752 | No | 106 kDa | No unique band; no change in Pms1 KO tissue |
| PMS1 | Novus NBP2-94782 | No | 106 kDa | No unique band; no change in Pms1 KO tissue |

**Table S3:** Mouse MMR antibodies screened and evaluated

| Target | Vendor | ID | Dilution |
|---|---|---|---|
| Msh3 | Santa Cruz | Sc-271079 | 1:500 |
| Mlh1 | Abcam | ab92312 | 1:500 |
| Exo1 | Abcam | ab155553 | 1:500 |
| Fan1 | University of Dundee | MTMR15 | 1:500 |
| Msh2 | Cell Signaling | #2017 | 1:1000 |
| Mlh3 | ProteinTech | #25298-1-AP | 1:200 |
| Pms2 | BD Pharmingen | 556415 | 1ug/ml |
| Msh6 | ThermoFisher | MA5-42406 | 1:500 |
| Pold1 | ThermoFisher | MA5-35464 | 1:500 |
| Pold3 | ThermoFisher | PA5-96616 | 1:500 |
| GAPDH | Millipore | MAB374 | 1:10000 |
| B-actin | Sigma | A5441 | 1:5000 |

**Table S4:** Western blot antibodies and dilutions to detect mouse MMR proteins

| UniProt Accession | UniProt Gene | Protein name | # of Sequences | # of Quant Values | Number of theoretical peptides (trypsin/P, 7-40) | Copy number / cell, Avg | Copy number / cell, 95% CI |
|---|---|---|---|---|---|---|---|
| Q00558 | F8a1 | 40-kDa huntingtin-associated protein | 5 | 11 | 15 | 316592 | 16653 |
| P00375 | Dhfr | Dihydrofolate reductase | 6 | 11 | 12 | 189522 | 17825 |
| Q6PEE3 | Rrm2b | Ribonucleoside-diphosphate reductase subunit M2 B | 9 | 11 | 17 | 187094 | 10205 |
| P42859 | Htt | Huntingtin | 73 | 11 | 154 | 114255 | 3410 |
| Q61164 | Ctcf | Transcriptional repressor CTCF | 4 | 11 | 25 | 109665 | 13125 |
| P17918 | Pcna | Proliferating cell nuclear antigen | 7 | 11 | 17 | 108872 | 15781 |
| Q8CGF7 | Tcerg1 | Transcription elongation regulator 1 | 21 | 11 | 41 | 79919 | 3724 |
| Q8VEE4 | Rpa1 | Replication protein A 70 kDa DNA-binding subunit | 15 | 11 | 32 | 70960 | 3719 |

| | | | | | | | |
|---|---|---|---|---|---|---|---|
| P43247 | Msh2 | DNA mismatch repair protein Msh2 | 18 | 11 | 46 | 57821 | 3657 |
| Q9WUK4 | Rfc2 | Replication factor C subunit 2 | 6 | 11 | 22 | 45917 | 3620 |
| O35654 | Pold2 | DNA polymerase delta subunit 2 | 3 | 11 | 21 | 44393 | 4906 |
| Q924H2 | Med15 | Mediator of RNA polymerase II transcription subunit 15 | 3 | 11 | 20 | 37037 | 4800 |
| Q99J62 | Rfc4 | Replication factor C subunit 4 | 6 | 11 | 19 | 32561 | 2079 |
| Q9D0F6 | Rfc5 | Replication factor C subunit 5 | 5 | 11 | 22 | 28052 | 3679 |
| Q9JKK8 | Atr | Serine/threonine-protein kinase ATR | 3 | 8 | 149 | 26479 | 11028 |
| Q9EQ28 | Pold3 | DNA polymerase delta subunit 3 | 3 | 11 | 25 | 22979 | 2714 |
| P49135 | Ercc3 | General transcription and DNA repair factor IIH helicase/translocase subunit XPB | 9 | 11 | 42 | 21628 | 994 |
| P97386 | Lig3 | DNA ligase 3 | 14 | 11 | 55 | 21476 | 807 |
| P07742 | Rrm1 | Ribonucleoside-diphosphate reductase large subunit | 13 | 11 | 44 | 19984 | 2542 |
| P52431 | Pold1 | DNA polymerase delta catalytic subunit | 9 | 11 | 64 | 16785 | 907 |
| Q9JK91 | Mlh1 | DNA mismatch repair protein Mlh1 | 4 | 11 | 49 | 13393 | 2627 |
| P54276 | Msh6 | DNA mismatch repair protein Msh6 | 18 | 11 | 74 | 11336 | 968 |
| P13705 | Msh3 | DNA mismatch repair protein Msh3 | 6 | 11 | 58 | 9793 | 877 |
| P35601 | Rfc1 | Replication factor C subunit 1 | 4 | 11 | 57 | 6458 | 793 |
| Q80TP3 | Ubr5 | E3 ubiquitin-protein ligase UBR5 | 18 | 11 | 134 | 6000 | 265 |
| Q69ZT1 | Fan1 | Fanconi-associated nuclease 1 | 1 | 6 | 49 | 5841 | 432 |
| Q8BTF7 | Lig4 | DNA ligase 4 | 2 | 2 | 44 | 3743 | 6 |
| P37913 | Lig1 | DNA ligase 1 | 2 | 5 | 50 | 3737 | 845 |
| P54279 | Pms2 | Mismatch repair endonuclease PMS2 | | | | BLQ | |

| | | | | | | |
|---|---|---|---|---|---|---|
| Q8K119 | Pms1 | PMS1 homolog 1, mismatch repair system component | | | | BLQ |
| A0A1Y7VMP7 | Mlh3 | MutL homolog 3 | | | | BLQ |
| Q9QZ11 | Exo1 | Exonuclease 1 | | | | BLQ |

**Table S5:** Peptide identification for DNA repair and handling targets

**Data S1:** Metadata annotation and proteins of interest lists

**Data S2:** Integrated preprocessing, quality control, differential analysis, visualization, and pathway enrichment

**Data S3:** Proteomic quantification including log2 fold-changes, p-values, adjusted p-values, derived percent-of-control values

**Data S4:** Proteome cluster-wise summaries exported together by spaghetti-profile plots

**Data S5:** Proteome clusters by biological process enrichment
